# Stabilising microbial communities by looped mass transfer

**DOI:** 10.1101/2021.08.19.456962

**Authors:** Shuang Li, Nafi’u Abdulkadir, Florian Schattenberg, Ulisses Nunes da Rocha, Volker Grimm, Susann Müller, Zishu Liu

**Author notes:** **Corresponding author:** Susann Müller, Helmholtz Centre for Environmental Research-UFZ, Department of Environmental Microbiology, Permoserstr. 15, 04318 Leipzig, Germany. **Email addresses:** Shuang Li. Ulisses Nunes das Rocha. Susann Müller.

## Abstract

Creating structurally and functionally stable microbiomes would be greatly beneficial to biotechnology and human health but so far has proven challenging. Here, we propose a looped mass transfer design that keeps microbiomes constant over long periods of time. The effluent of five parallel reactors that began with the same inoculum, was mixed in a reactor that represented a regional pool. Part of this pool was transferred back to the five reactors. Community dynamics were monitored and visualized by quantitative microbial flow cytometry and selected taxonomic sequencing of whole communities and sorted subcommunities. The rescue effect, known from metacommunity theory, was the main stabilizing mechanism that led to the survival of subcommunities with zero netgrowth, especially at high mass transfer rates. The looped mass transfer approach promises to overcome notorious stochastic structural fluctuations in bioreactors and has the potential to design and stabilize communities that can perform desired functions.

## Introduction

The ability to create structurally and functionally stable microbiomes would be greatly beneficial to both industrial biotechnology and human health. Stable microbiomes would be persistent in their composition and hence their functions and services^1-3^. Building and changing a microbiome as desired and maintaining it over hundreds of generations is a challenging endeavour that still needs to be undertaken. Stable microbiomes would make it much easier to use renewable and less-expensive resources and develop bio-based technologies as part of a circular bio-economy^4, 5^. Bottom-up and top-down approaches that are based on design-build-test-learn cycles^6^ to optimize artificial microbiome blueprints are promising potential avenues for the construction of human life-promoting microbiomes^7^. However, the current means of controlling microbiomes are still limited. For decades, scientists have attempted to homogenize^8^ or establish stable natural and artificial communities by shaping their niches, e.g., via substrate type and concentration, media composition, or pH, and temperature, including using machine learning and other mathematically based approaches^9-12^. Despite all these efforts, however, complex microbiomes appear to remain susceptible to large stochastic variations.

To establish stable microbiomes, a first challenge is to quantify community composition and its change. Liu et al.^9^ combined quantitative single-cell and taxonomic analyses to characterize microbial communities that comprised hundreds of species. However, when applying their approach to five parallel identically operated steady-state reactors that began from the same inoculum, they found that the five communities developed along different trajectories, leading to distinctly different compositions. One primary reason for this, as suggested by Liu et al.^9^, was the ongoing disturbance in each reactor via dilution. Dilution reduced the abundance of otherwise dominant species and caused the extinction of species with low growth rates, thereby significantly increasing the role of stochasticity. Functional redundancy supported this process. Zhou et al.^13^ also found 14 distinctly different communities that developed from the same inoculum, and referred to as ‘stochastic assembly’.

The important role of stochasticity in understanding community assembly has only recently been acknowledged in ecology^14, 15^. Stochastic variation in abundance may be ignored for large populations. For smaller ones, however, it can cause extinction^16, 17^. Nonetheless, rank-abundance data of virtually all known communities show that, except for a few dominant species, most species occur at low abundances^18-22^ and thus are strongly affected by stochasticity.

How, then, do the more or less stable communities develop which can be observed in macroecology? An answer can be found in metapopulation theory^23^, which is also incorporated in metacommunity theory^24, 25^. In metapopulations, small local populations (1) may go extinct but be replaced by recolonization, or (2) may be ‘rescued’ from extinction by immigration. In metacommunity theory, the former mechanism is referred to as “patch dynamics”^26^. The latter is referred to as ‘mass effect/transfer’^27^ or ‘rescue effect’^28^. In both cases, the regional pool that comprised all of local populations buffers and thereby stabilizes local populations.

A second challenge in stabilizing microbial communities is: How can we mimic the stabilizing effect of the regional pool? The patch dynamics mechanism requires limited dispersal to avoid the possibility that local dynamics are overly synchronized and their disparity is kept exclusive. In contrast, the rescue effect requires more substantial dispersal from the regional pool and supports the survival of local populations that would otherwise have too low an abundance and thus be at high risk of extinction. Therefore, we hypothesize that mass transfer from a regional pool could be a means to stabilize microbial communities. Mass transfers are a well-known phenomenon that occurs in the intestinal system^29^ and in wastewater treatment plants^19, 30^. Finding a simple but efficient way to implement mass transfer from a regional pool could thus be a promising avenue for understanding and building stable microbial communities with specific desired functions.

Based on the setup that was used by Liu et al.^9^, we implemented a regional pool that was connected to five parallel local communities via a loop design. Inflow from the local reactors represents emigration to the regional pool, and feedback flow represents mass transfer from the regional pool to each local community.

This metacommunity that comprised six continuously running and interconnected bioreactors was established and studied for approximately 80 generations with varying mass transfer rates. To quantify community dynamics, flow cytometry was used as the major methodology, which quantitatively interprets community structure at the single-cell level. Additionally, selected 16S rRNA gene amplicon sequencing was performed for whole communities and after the directed sorting of subcommunities. We hypothesise that (i) diversity values will become more equal with higher rates of mass transfer, (ii) an increase in mass transfer will strengthen the presence of slow growing organisms, and (iii) high mass transfer rates will increase the stability properties of constancy, resistance, and recovery^3^.

## Results

### Mass transfer stabilizes microbiomes

Two hierarchical scales (i.e. local and regional scale) were designed. For the local scale, independently inoculated and connected microbiomes were cultivated in five identically operated local reactors (i.e., local communities L1-L5; Supplementary Information S1). For the regional scale, a pool R of emigrated microorganisms from L1-L5 was formed from which dispersion back into the local scale was allowed (Fig. 1, Supplementary Information S2). We followed this setup for approximately 80 generations, and a total of 448 community samples [with a total of 35,840 subcommunities (SC)] were collected from the six bioreactors (Supplementary Information S3). Quantitative variations in the microbiomes were analysed at the single-cell level using flow cytometry (Supplementary Information S4, S5) and relative variations were analysed by 16S rRNA sequencing (Supplementary Information S6). With this setup, we sought to determine whether changes in recycling rates *RC* (Table S2.1) may be a means to stabilize microbiomes.

**Figure 1.**
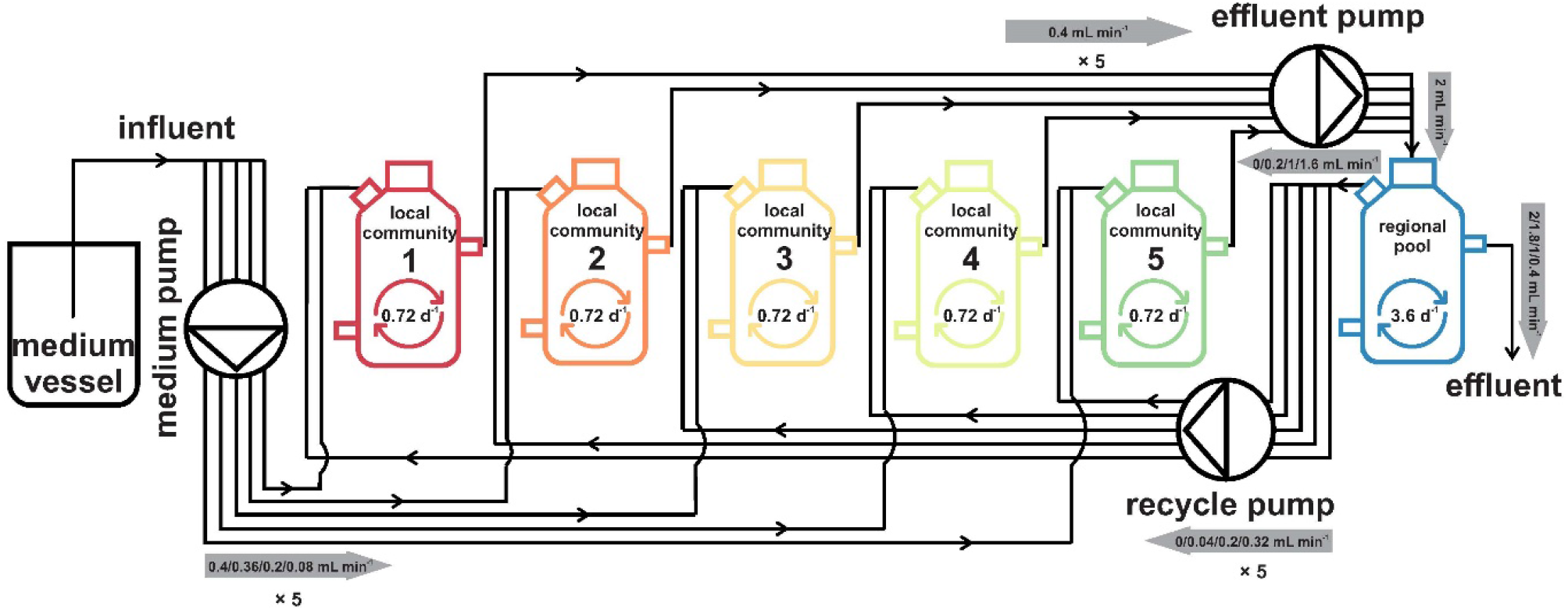
Scheme of the reactor setup. The six reactors were run under the same environmental conditions. The five local community reactors L1-L5 were run in an identical mode. The sixth reactor served as the regional pool R and was fed by the effluents of the local communities L1-L5. The dilution rate per reactor is shown within the reactor schemes. A medium pump controlled the medium flow rate from the medium vessel to local communities L1-L5. An effluent pump controlled the effluent rate from the local communities to the regional pool R. A recycle pump controlled the recycling flow rate from the regional pool R to the local communities L1-L5. The flow rates are labelled with a grey arrow that indicates the flow direction.

The experiment was divided into five phases, in which increasing recycling rates *RC* between the regional pool R and local reactors were tested in phases 2-4, whereas no exchange was allowed in phases 1 and 5. Phase 1 (Insular I) showed large variations in microbiome structures between local communities in L1-L5 despite an identical inoculum. These variations were sequentially reduced when recycling rates *RC*were increased from 10% to 80% (Fig. 2a). Additionally, temporal variation within each reactor decreased with an increase in *RC* (Fig. 2b). Fluctuations reappeared after recycling was terminated (phase 5, Insular II; Fig. 2a). An overview on the experiment is provided in video S1.

**Figure 2.**
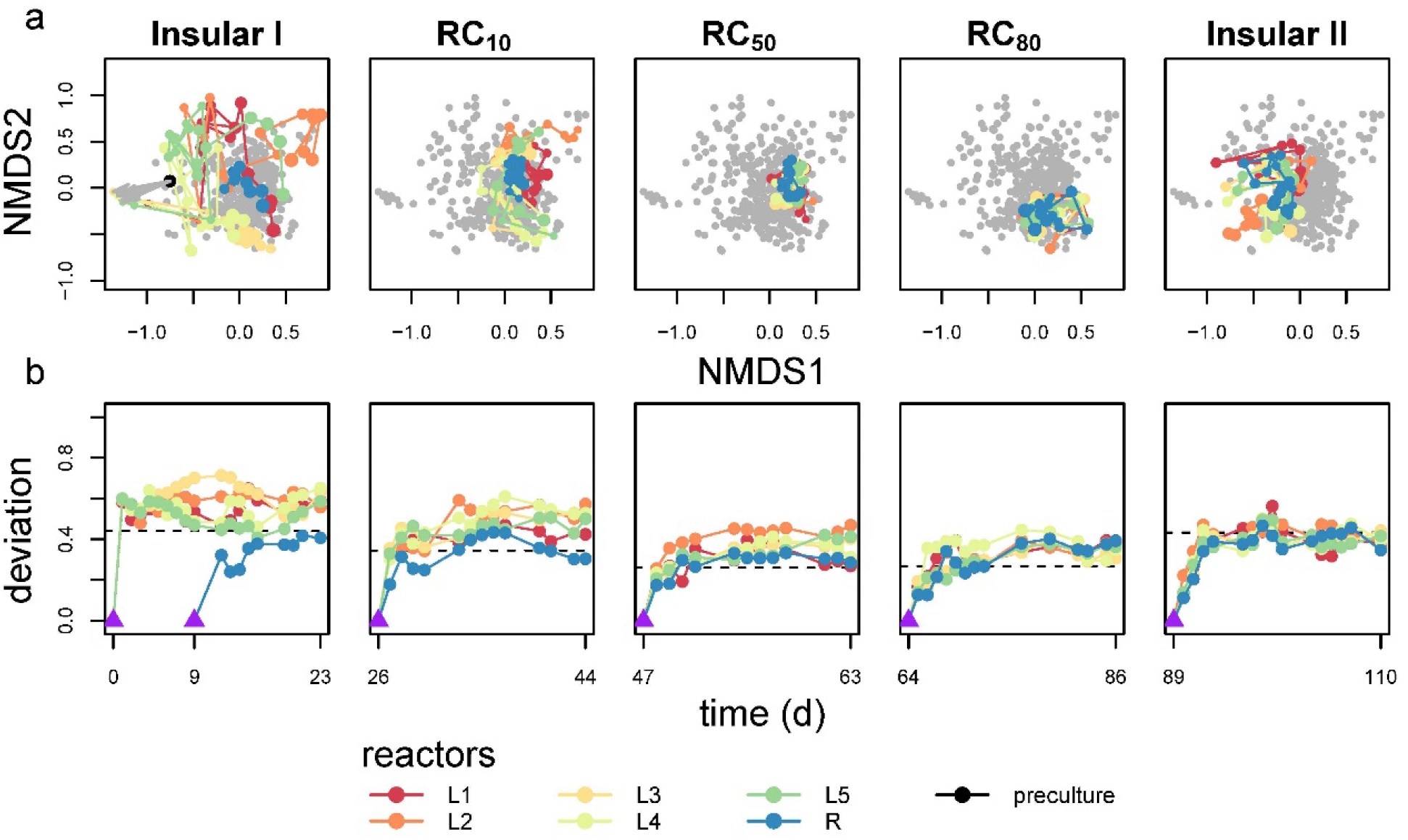
Microbiome dynamics in the five reactors L1-L5 and regional pool R. Microbiome dynamics were measured by flow cytometry. The plots represent the five different phases of the experiment. In phase 1 (Insular I), L1-L5 were isolated. In phases 2-4, the recycling flow rate from reactor R increased sequentially from 10% to 80% (RC_10_-RC_80_). Phase 5 (Insular II) was again without recycling. **(a)** The NMDS plots in the upper row show the increasing similarity of communities, both within and between reactors, with increasing recycling rate *RC*, while similarity was quickly lost when recycling from reactor R stopped. Connected time points indicate the assembly trajectory of microbiomes. Points in grey represent samples from the other phases. The NMDS plots were created using relative cell numbers of all SCs based on Bray-Curtis distance measure (try = 100, trymax = 200). **(b)** Deviation of microbiome structure from the endpoints of respective previous phases (purple triangle) based on Canberra distance. The purple triangle in the Insular I phase represents the inoculum.

The stability properties of constancy, resistance RS, and recovery RV^2^, calculated for all microbiomes L1-L5 and R, supported these findings. The five local microbiomes established the highest constancy at the highest recycling rates *RC* (Fig. S7.1a, Table S7.1). The values for resistance RS were lowest between the Insular I to RC_10_ phases (mean = 0.46 ± 0.05 for L1-L5), suggesting considerable variations in local microbiomes under these conditions. During transitions from RC_10_ to RC_50_ and from RC_50_ to RC_80_, the highest resistance RS values (mean = 0.57 ± 0.03 and 0.60 ± 0.03, respectively, for L1-L5) were found and thus represented the most stable conditions (Fig. S7.1b). The values for recovery RV were low in all phases and for all microbiomes (Table S7.2), indicating that the communities did not return to compositions of previous phases.

In summary, the data suggest that high mass transfer rates clearly lowered local and temporal variations in community composition and supported such stabilizing properties as high constancy and increasing resistance of all microbiomes, whereas recovery efforts to reinstate a previous microbiome remained unimportant.

### Mass transfer synchronizes microbiomes

The synchronization of microbiome structures between local communities L1-L5 and beyond with the regional pool was tested by calculating a range of diversity values (Supplementary Information S8). With the exception of inventory α-diversity, we found that mass transfer strongly affected these values. All local microbiomes L1-L5 did not appreciably change in their richness during the 110 days of continuous cultivation, but γ-diversity, expressed as the richness of SCs in the metacommunity, was lowest at RC_80_ and highest in the Insular I and II phases (Table 1, Figs. S8.4, S8.5), suggesting that the same SCs were dominating L1-L5 in phase 4 (RC_80_).

**Table 1.** Summary of α-diversity, γ-diversity, intra-community β-diversity, inter-community β-diversity (L vs. L), and inter-community β-diversity (L vs. R) as mean ± standard deviation (sd) per phase. The mean ± sd values were all calculated among local microbiomes L1-L5 during balanced periods if not stated otherwise. The values marked with stars behind the diversity values in these phases were significantly different from those in any other phase. \**p* ⩽ 0.05, \*\**p* ⩽ 0.01, \*\*\**p* ⩽ 0.001.

| diversity/phase | Insular I | RC<sub>10</sub> | RC<sub>50</sub> | RC<sub>80</sub> | Insular II |
| --- | --- | --- | --- | --- | --- |
| α-diversity | 15.87 ± 4.42 | 15.84 ± 2.90 | 15.48 ± 1.71 | 15.52 ± 2.63 | 17.71 ± 3.18* |
| γ-diversity | 39.73 ± 3.17*** | 31.00 ± 3.08 | 22.00 ± 2.73* | 18.80 ± 3.85* | 33.11 ± 3.62 |
| intra-community β-diversity | 8.63 ± 4.07 | 8.04 ± 3.65 | 5.56 ± 3.60 | 4.70 ± 2.52 | 10.56 ± 4.86 |
| inter-community β-diversity (L vs. L) | 19.58 ± 5.29*** | 13.58 ± 4.09 | 5.43 ± 3.36*** | 2.58 ± 1.65*** | 12.24 ± 5.20 |
| inter-community β-diversity (L vs. R) | 17.50 ± 4.23*** | 12.20 ± 3.58** | 6.50 ± 3.03*** | 2.43 ± 1.82*** | 9.73 ± 4.54** |

Temporal intra-community β-diversity values highlight structure variations per community. The highest values were found on day 3 for each of L1-L5 (35.8 ± 3.42), indicating rapid changes in L1-L5 during the adaptation period in the Insular I phase (Fig. 3a). For balanced periods within each phase, the intra-community β-diversity values for the Insular I and II phases (8.63 ± 4.07 and 10.56 ± 4.86, respectively) and especially for RC_50_ and RC_80_ (5.56 ± 3.60 and 4.70 ± 2.52, respectively) were significantly lower (Table 1).

**Figure 3.**
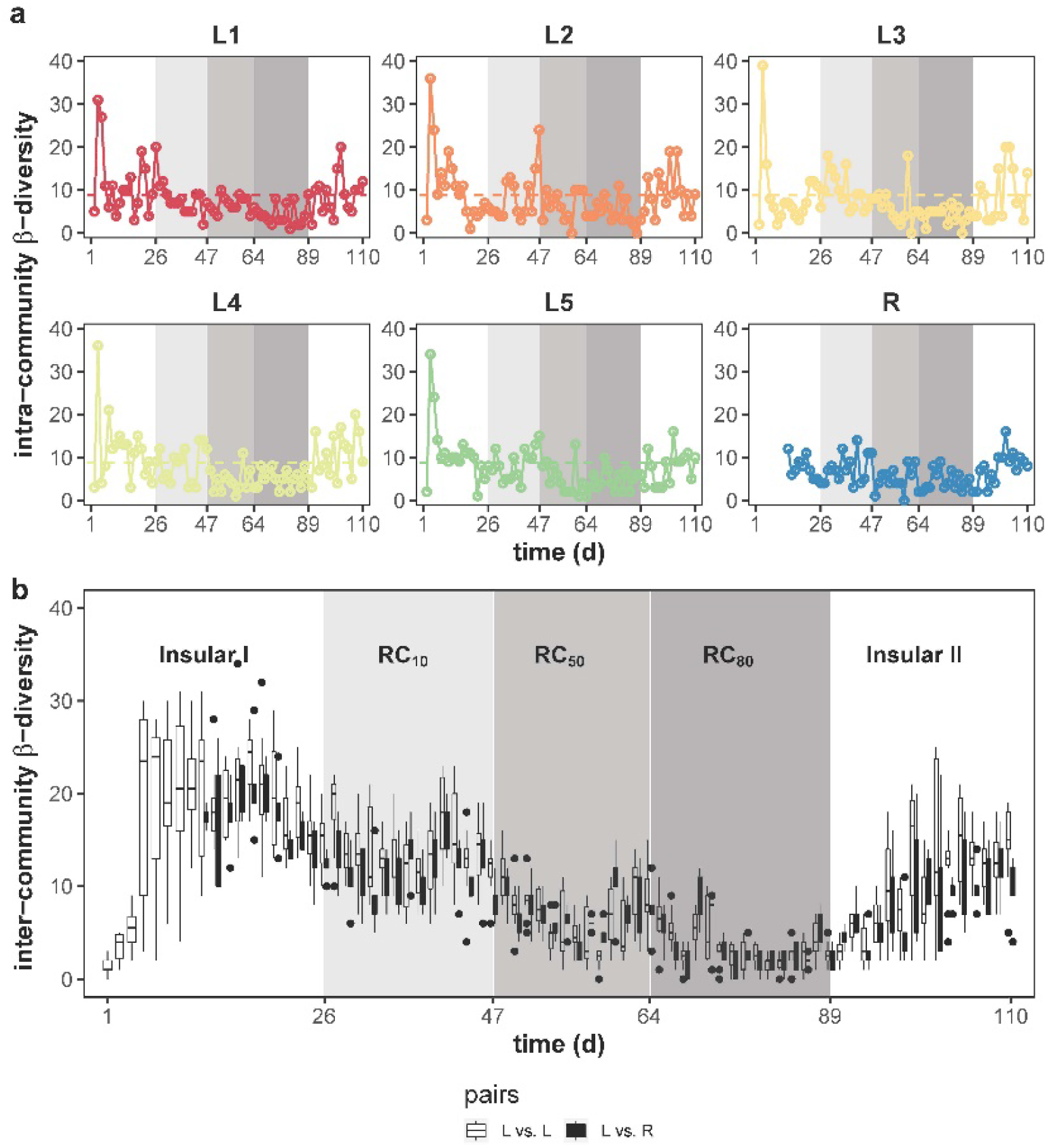
Intra- and inter-community β-diversity characteristics of local microbiomes L1-L5 and the regional pool R. **(a)** Variation of intra-community β-diversity over time. The number of unique dominant SCs was determined in pairwise successive samples within a microbiome. The threshold value was 8.76 and used to distinguish drift events from intrinsic cellular fluctuations during balanced growth condition (horizontal dashed line). The intra-community β-diversity values were reduced and showed fewer drift events under RC_50_ and RC_80_ conditions compared with other phases. **(b)** Variation of inter-community β-diversity over time. Numbers of SCs were determined that were not shared in pairwise samples at the same time point between local microbiomes L vs. L (open symbols) and L vs. R (closed symbols). An increase in recycling rates *RC* lowered inter-community β-diversity to a high degree. The shaded areas represent different phases with changes in *RC*.

Temporal intra-community β-diversity variations were also a means to find stochastic structural changes (i.e., drifts) in L1-L5 that occurred during the balanced periods under *RC*. If variations surpass a certain threshold, defined by the mean intra-community β-diversity value that was determined for the balanced periods of the first phases of all L1-L5 (8.76, dashed lines in Fig. 3a), then they were considered drifts. Drifts were observed more frequently in the insular phases (Fig. 3a, Tables S8.1, S8.2) and were lowest for RC_80_. Additionally, this synchrony at RC_80_ was even less deranged when looking at 16S rRNA sequencing data of sorted SCs (Fig. S9.3). Some of the drifts were apparently caused by different physiological cell states of dominant SCs but not by genotypic taxon variation (e.g., *Azospirillum* switching between six SCs with different growth states; Supplementary Information S9). Stochastic changes in community structures thus appear to be significantly suppressed by mass transfer.

Inter-community β-diversity was used to compare variations between L1-L5 and R at each time point. This diversity parameter was also highly sensitive to the recycling flow rate *RC* and significantly declined from the Insular I phase (19.58 ± 5.29) to RC_80_ (2.58 ± 1.65) and increased again in the Insular II phase (12.24 ± 5.20; Table 1, Fig. 3b). The taxonomic composition of L1-L5 and R (genus level) was clearly different between the phases, with the exception of RC_50_ and RC_80_ (Fig. S9.1), thus supporting the synchrony in microbiome compositions that was shown by the cytometric data.

Furthermore, the degree of exchange of particular SCs by other SCs, which is described by the term ‘turnover’ β_SIM_ was also high for the Insular I and II phases but only half as much for the RC_80_ (i.e., multisite intra-community β-diversity; 0.34 ± 0.02; Table S8.3). In contrast, ‘nestedness’ β_NES_, a value that describes the local and temporal persistence of SCs in communities^31^, was highest at RC_80_ (0.14 ± 0.02; Fig. S8.6a, Table S8.3). The type of the persistent SCs is presented in Table S8.4. Thus, the strong compositional synchronization of the local microbiomes at RC_80_ was further supported by the determination of β_SIM_ and β_NES_ values, as the high turnover of SCs was observed only in the Insular I and II phases, but an increasing nestedness at RC_80_ (Table S8.4).

Thus, although α-diversity richness values of local communities did not change, γ-diversity and inter-community β-diversity were lowest at RC_80_. At the end of RC_80_ (day 86), only 13 out of 80 SCs were dominating the final microbiomes in L1-L5 and R, with 54.7-65.6% of all cells in high synchrony. The intra-community β-diversity indicated the lowest variation and lowest drift events at RC_80_, which was also proven by whole-community 16S rRNA gene sequencing (Fig. S9.1), for community trend analysis (Fig. S9.2), and drift events (Fig. S9.3). The 16S rRNA gene sequencing data supported and reinforced the cytometric data.

### Mass transfer allows the survival of cells with a low or zero netgrowth rate in microbiomes

Subcommunities were found to survive in the reactors even when their netgrowth rate µ′ was lower than the prevailing dilution rate *D* = 0.72 d^-1^.

The following ideal values for the mass transfer rate, *M*, were expected for the proposed experimental setup (Figs. S10.1, S10.2) when conditions assumed balanced situations (i.e., constant cell numbers) and when the cell numbers of the inflow (i.e., regional pool) were equal to cell numbers in the local microbiome. *M* should change with the recycling rate *RC*: *M* = 0.072 d^-1^ (RC_10_), *M* = 0.36 d^-1^ (RC_50_), and *M* = 0.576 d^-1^ (RC_80_; Eq. S10.8.). Corresponding to µ′ = *D* ™ *M* (Eq. S10.9), the netgrowth rate can theoretically be calculated to be 0.648 d^-1^ (RC_10_), 0.36 d^-1^ (RC_50_) and 0.144 d^-1^ (RC_80_; Fig. S11.1a). The actual *M* values in the experiment closely followed the theoretical calculations (Table S10.1), whereas the netgrowth rate µ′ showed a few variations because of fluctuations in cell numbers (Table S10.1, Fig. S11.1c). These results demonstrate that at high *RC* and high *M* values, the netgrowth rate µ′ decreased sharply.

Similar to µ′ for the whole community, the netgrowth rates of the subcommunities, µ′ *SC*_*x*_, can be calculated (Eqs. S11.1-11.5). µ′ *SC*_*x*_ decreased with increasing *RC*, partly to zero (Fig. 4, blue points), and the number of SCs in all local microbiomes with µ′ *SC*_*x*_ ⩾ 0 decreased from 305 out of 305 SCs in the Insular I phase to 124 out of 155 SCs at RC_80_ and increased back to 280 in the Insular II phase (Fig. S11.1d). The number of SCs with µ′ *SC*_*x*_ ⩾ 0 per local microbiome at RC_80_ was similar to those in the regional pool, R (23 SCs), which was much lower than the other phases (Fig. 4).

Subcommunities remained in the reactor setup at RC_80_ because of netgrowth in at least one of the five local microbiomes (Fig. 4, red points for L1-L5) and were rescued by their redistribution via the regional pool R. Some low-growing or non-growing SCs have also increased their abundance in the regional pool R (Fig. 4, red points for R). Some of the rescued SCs were cytometrically sorted and taxonomically sequenced. For example, at RC_80_, *Azospirillum* (G33), *Azospira* (G18), and *Ochrobactrum* (G1; Fig. S9.5) indicated netgrowth in all or most of the L1-L5 (Fig. 4, Supplementary Information S9), whereas an unassigned genus from PeM15 (order, G12 and G14) accumulated in R, and cells of *Sphingobacteriaceae* (family, G4) and *Pseudacidovorax* (G2, G11; Fig. S9.5) grew in few of the microbiomes L1-L5 and R and all were rescued by redistribution using R. These SCs were mainly mono-dominant and could be assumed to be superior local competitors under RC_80_ and R. Some other SCs that were also among the growing SCs (G5, G9, G25; Fig. S9.5, Table S9.2) were multi-dominant. Generally, fewer SCs responded to mass transfer at RC_80_ but did so with higher relative abundance per SC (Fig. S12.2). When performing correlations between absolute cell number per SC (Fig. S8.2) and *RC*, five SCs were revealed that all also showed netgrowth or rescue characteristics (G5, G12, G13, G14, G33; Fig. S12.1). Among the growing and rescued SCs were those that were nested. At RC_80_, flow cytometrically determined nested SCs were G2, G4, G5, G9, G12, and G14 (Table S8.4), and the 16S rRNA analysis determined *Pseudacidovorax* (G2), PeM15 (order, G12, G14), and multi-dominant G5 and G9 for these SCs (Fig. S9.5).

**Figure 4.**
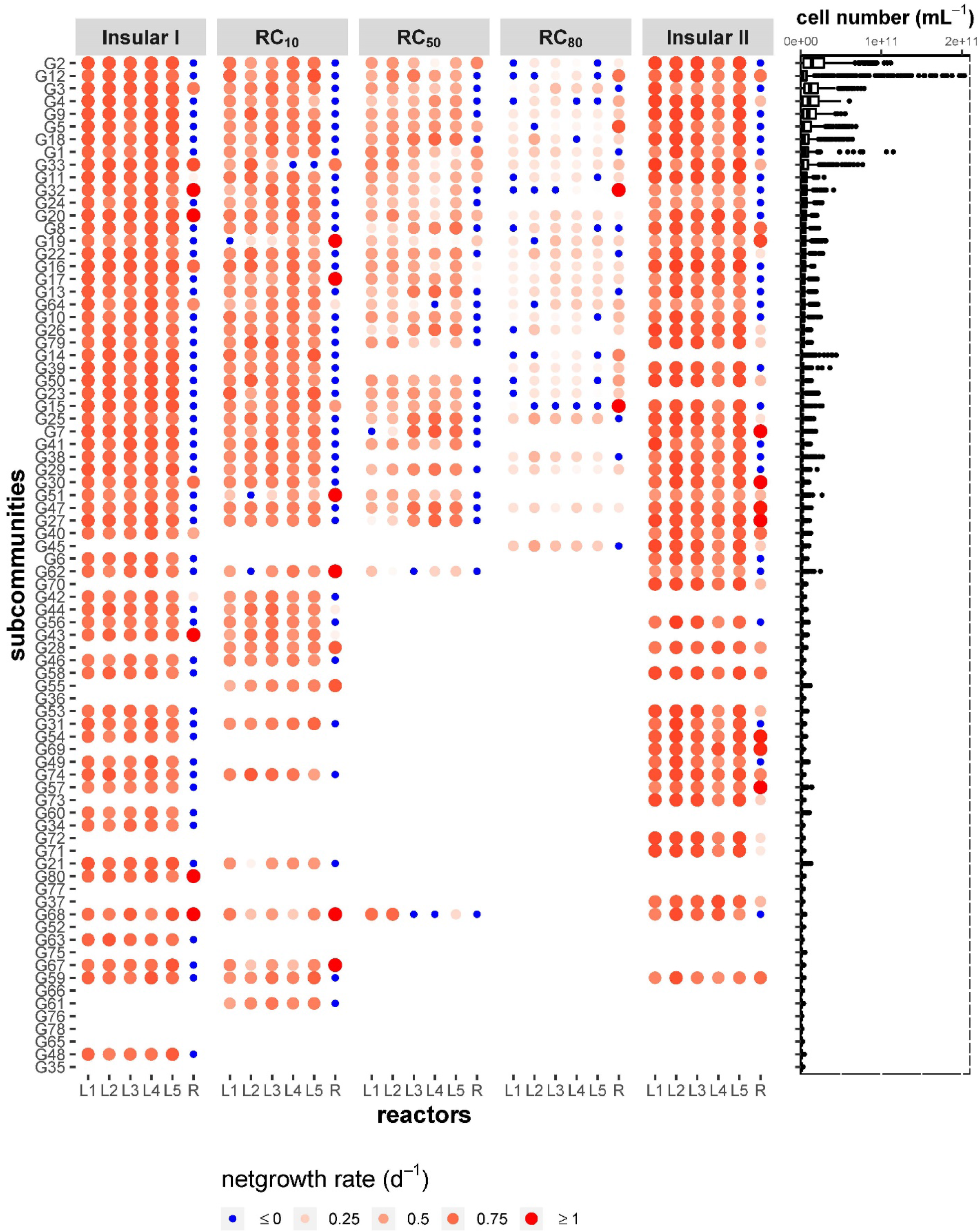
Cell numbers of SCs and netgrowth rates µ′ *SC*_*x*_ (x = 1-80), determined for the balanced periods of the different phases in the local communities L1-L5 and the regional pool (R). On the left, the netgrowth rate µ′ *SC*_*x*_ (d^-1^) is presented only for dominant SCs with relative abundances > 1.25 in at least one sample during the corresponding periods in L1-L5 and R. The colour gradient and size of the red circles represent the value of positive µ′ *SC*_*x*_. Darker colours and larger circles indicate higher µ′ *SC*_*x*_ values for positive netgrowth. All blue circles represent zero netgrowth µ′ *SC*_*x*_. The SCs are ranked in descending order according to their mean cell number (cells mL^-1^) for all days and all reactors. Cell numbers of x = 80 SCs of a total of 421 samples are shown as a boxplot on the right. Outliers of deviated cell number values are shown as points.

Thus, the mass transfer supported genera that showed netgrowth µ′ *SC*_*x*_ in at least one or a few of the local communities and were rescued by redistribution to all five communities through the regional pool R, or accumulated in R and were also redistributed via R to the local communities. Subcommunities that exhibited netgrowth were also nested. These mechanism ensured the survival of cells, even if they did not grow or grew slowlier in local microbiomes than the actual dilution rate *D* would allow.

## Discussion

Stochasticity and dispersal are essential processes in shaping (meta)community structure and function in natural environments. Both random drift events, which influence the birth and death of organisms, and the extent to which organisms disperse are key factors in how communities are affected at local and temporal scales.

The absence of dispersal results in communities that are isolated from each other, making them susceptible to stochastic variation^9, 10, 13^. Low dispersal is common in macroecology and contributes to the presence and coexistence of multiple species within a metacommunity between patches^32, 33^. In contrast, high dispersal is known to increase similarity between localities and thus the risk of global extinction^34-36^. High dispersal has a quantitative effect on local community formation in accordance to the mass-effect paradigm, in which community composition in sink habitats tends to resemble that of the source habitat^37, 38^. Frequently, the abundance of species in source communities and their transfer rates influence the composition of sink habitats^39^, and lost individuals are replaced by members of source communities^40^. This underscores the assumption that a regional pool that floods local communities with high rates of mass transfer can lead to synchrony and stability.

Our study demonstrated that the invented loop-designed mass transfer stabilized the microbiomes and increased synchrony between the six localities. The term ‘stability’ is described by various properties, out of which constancy, resistance, and recovery are the most essential ones^1^. We found greater constancy and resistance values in L1-L5 at RC_50_ and RC_80_, reflecting stability of the community to stochastic assembly processes and perturbation events (i.e., dilution rate). The source-sink relationship between L1-L5 and the regional pool R created a reduction of the species pool but at high cell abundance that provided less space for neutral forces. Long-term constancy was established when transfer rates at RC_50_ and RC_80_ prevented the extinction of local species through the rescue effect and when regional equality was achieved according to Chesson^41^. The increase in resistance that was observed in the present study could also be the result of the gradual increase in mass transfer rates (i.e., the RC_10_ phase selected already partially nested SCs that eventually dominated at RC_50_ and RC_80_, such as G2; Table S8.4). Recovery by definition describes the ability of a community to return to the constancy space after a temporary disturbance^1, 2^. We found that recovery values were always low because the microbiomes were intentionally steered towards a low-diverse but unchanging state through the use of the loop-designed metacommunity setup. Nonetheless, recovery values were always positive, despite permanent changes in mass transfer rates and because of reinforcement of the rescue effect, which, in addition to the high constancy and resistance values, demonstrates the high stability properties of the metacommunity at RC_50_ and RC_80_.

Usually, mass transfer occurs between ecologically distinct niches in natural and engineered systems. Even when mass transfer is high, different environments have specific community structures (e.g., discrete bacterioplankton communities in a lake and its inlets^25^ or microbial communities in connected transects of wastewater treatment plants^19, 30^). In our metacommunity, identical localities that operated in the same way were also expected to develop disparity^2^, but these were overcome by the loop design. The looped mass transfer caused synchrony between L1-L5 and R and minimized the ability to form independent and disparate microenvironments. Separate local niches that originally formed during the first 26 days of the experimental setup elapsed. After, the microbiomes in L1-L5 and R formed a mutual niche where the same core microbiomes always dominated (Fig. S12.2). These metacommunity niche-shaping effects were major events that caused the acute synchrony of the L1-L1 and R microbiomes and the transitional loss of cell types that were previously dominant in the insular phase.

The high mass transfer rates we applied were able to nearly eradicate local neutral and discrete species-sorting forces, allowing only a few members of the microbiome to grow synchronously at all locations. Notably, relief from mass transfer allowed the renewal of earlier diversity levels (Insular I phase vs. Insular II phase), indicating that rare species were still present to allow diversity recovery.

We found that high mass transfer rates affected diversity values and reinforced specific cell types. Looped high mass transfer did not alter richness in terms of number of dominant SCs or individual cell production per reactor much (PC; Table S10.1). But it reduced β-diversity between and within microbiomes and resulted in the loss of local niches, which in turn reduced γ-diversity (Table 1). As a consequence of the lower number of cell types, high mass transfer resulted in lower stochastic drift events (Table S8.1) and thus narrower established constancy spaces (Fig. 2b). Fewer or even no drift events were found in the balanced growth periods, indicating that neutral forces in the metacommunity were low, differing from insular environments^9, 10^. Random birth and death events were limited by the continuous inflow of source organisms from the regional pool R and by the increase in biomass with increasing mass transfer (Fig. S3.1). High biomass is known to lower susceptibility to demographic drift or disturbance^42^.

Another argument in favour of the power of mass transfer to prevent variation was the low degree of the exchange, β_SIM_ (turnover), of SCs within and between microbiomes at high mass transfer rates, which was different for insular situations (Fig. S8.6, Table S8.3). Conditions of strong mass transfer have been described to moderate the turnover of community structures and support the nestedness of species (e.g., in the process of biotic homogenization^43^ or post-glacial recolonization processes of northern biotopes^44^). Most SCs that benefitted from mass transfer were also those that showed persistence, β_NES_ (nestedness; Table S8.4). Similar to core species, they could also play an important role in maintaining community traits. Core species are common and dominant (e.g., in gut microbiomes^45^, in benthic octocoral associations in the form of symbiosis^46^, and species in macroecology that cope with climate warming, which largely determines temporal stability of the total biomass in alpine grassland communities^47^). In our loop design, dominant SCs indicated an increased number of correlations despite nutrient limitation that was caused by high cell density during RC_50_ and RC_80_ in L1-L5 and especially in the regional pool R (Table S13.1). Genera in nested SCs were able to survive in R (Table S8.4) because they could potentially cope with the low carbon and ammonium resources, such as *Sphingobacteriaceae* (Fig. S9.1) ^48-50^ or PeM15^51^. Nested *Azospirillum* (G33), *Azospira* (G18) and *Pseudacidovorax* (G2, G11; Fig. S9.5) are nitrogen fixers. They may have displaced other genera because of ammonium self-sufficiency^52-54^. Thus, the regional pool R might have acted as a ‘hotspot’sphin for genus and function selection, which underscores the possibility of selecting desired functions by modifying the conditions of the regional pool.

We also found that mass transfer supported slow-growing organisms through the rescue effect. Mass transfer is a source-sink relationship and contributes to the spread and survival of species in sinks that would otherwise go extinct^55^. Source-sink relationships are generally very strong to ensure coexistence, and modelled environmental variations were found to not affect them^56-58^. The rescue of species at a sink site is reflected in various biotechnological processes where biotechnologists have successfully used this source-sink principle for bio-augmentation to keep a desired species in a system^59, 60^.

Most studies, however, ignore or cannot distinguish between the niche-specific competitive hierarchy of microbiome members in sources or sinks and the effects of emigration and immigration on these relationships. Recent studies have begun to track sink members within microbiomes by calculating their netgrowth based on the relative abundances of 16S rRNA gene sequencing data and bulk biomass to approximate their contribution to sink biomass formation^61, 62^. Our study goes a step further by calculating the netgrowth of all cell types that migrated back and forth within the loop-designed source-sink metacommunity, based on data from individual cells.

We found that slow-growing or almost non-growing cell types, that would typically be washed-out under continuous feeding conditions^9^, remained part of the microbiome through the rescue effect (Fig. 4). At the extreme, at RC_80_, when the metacommunity was fully mixed and when the final microbiome was established, the growth of some SCs in L1-L5 was especially low or nonexistent because of an increase in biomass and decrease in nutrient resources. However, these SCs overcame their non-growth in one or two reactors by growing in one or more of the other five local reactors and, subsequently, by redistribution via R, which enhanced their persistence in the metacommunity. Similar rescue effects were modeled for a macroecology background in relation to a meta-food-web, suggesting that biodiversity can be buffered under global change^63^. The source-sink relationship from local communities L1-L5 to the regional pool R and its reverse loop supported the growth of SCs in L1-L5 rather than in nutrient-limited R. Unknown is why cells were nested in R and rescued through R. The few SCs (e.g., multi-dominant G5 and PeM15 in G12, G14) showed small cell morphologies. It is conceivable that they might have been able to successfully utilize the limited resources and simultaneously take advantage of the high-exchange-rate (*D* = 3.6 d^-1^) qualities of R.

Our setup was thus able to protect slow-growing or even non-growing microorganisms for at least 80 generations, which otherwise would not have survived without mass transfer and the rescue effect. The loop-designed metacommunity may thus be a tool to protect and preserve functionally valuable microorganisms even if their growth rate is slow and lower than the prevailing dilution rate.

In summary, looped mass transfer is a means of stabilizing microbial communities over long periods of time. The degree of stabilization can be selected via the mass transfer rate *RC*. Mass transfer i) reduced local and temporal variations, and the stochastic behaviour that is normally observed in insular setups was reduced. All microbiomes showed high constancy and increasing resistance at high mass transfer rates. Mass transfer ii) synchronized structures of the microbiomes, resulting in the lowest inter-community β-diversity at the highest mass transfer. The variation of β-diversity within communities ceased, and the persistence of particular SCs was highest at high mass transfer. High turnover of community structures was observed only when no mass transfer occurred. An increase in mass transfer iii) increased cell numbers, thereby decreasing netgrowth rates µ′. Subcommunities that showed no growth (µ′ *SC*_*x*_ = 0) in one locality were rescued by growth at another locality and by their redistribution via the loop design. Thus, lost SCs, whose growth rate was below the dilution rate and that would normally go extinct, were fostered and replaced by members of the source community. The regional pool itself also served as a rescue site through the redistribution of SCs that accumulated specifically in R.

The local reactor conditions that were used in this study ultimately selected our microbiome. It is conceivable to test other local and especially regional conditions that support other cell types in natural communities in the future and thus design other stabilized communities. In particular, when certain medically or biotechnologically relevant functions are desired that require organisms with different physiological properties, including different growth rates, our loop design provides a solution for long-term stabilization and thus the reliable functioning of microbiomes.

## Materials and Methods

The metacommunity consisted of five local communities (L1-L5) that were operated in parallel and identically in continuous flow reactors (Fig. 1). To establish mass transfer between L1-L5, effluents from all five bioreactors were combined in a sixth bioreactor (i.e., the regional pool R). After mixing, a part of the regional pool R was returned to L1-L5 via a recycling loop.

In accordance to Liu et al.^9^, the cultivation of L1-L5 began simultaneously with an identical inoculum that originated from a full-scale wastewater treatment plant (Supplementary Information S1). To study the effects of mass transfer, recycling flow rates *RC* were applied with 10% (RC_10_), 50% (RC_50_) and 80% (RC_80_) of the original medium feeding rate of 0.4 mL min^-1^ (Supplementary Information S2).

The experiment was run for 110 days. Five phases were distinguished in accordance to to their recycling flow rate *RC*: phase 1 with no recycling (Insular I, starting on day 0), phases 2-4 (with recycling flow rates RC_10_, RC_50_, and RC_80_, starting on days 26, 47, and 64, respectively), and phase 5 (Insular II, starting on day 89), again without recycling. Within each phase, the first 7 days (5-times volume exchange of a reactor) were defined as the adaptation period, and the subsequent days were a balanced period during which abiotic parameters were expected to be more or less constant (Supplementary Information S3).

A total of 448 samples were collected from the six bioreactors, (i.e., 76 samples from each of the five local communities L1-L5 and 68 samples from the regional pool R; Supplementary Information S3, Dataset S3). The harvested cells were stabilized, stored at -20°C, and stained with 4’,6-diamidino-2-phenylindole (DAPI) for flow cytometric measurement (Supplementary Information S4). The samples were analysed with a MoFlo Legacy Cell Sorter (Beckman Coulter, Brea, CA, USA). To ensure reliability of the cell handling procedures, a microbial cytometric mock community (mCMC)^64^ was used, which was handled identically to the reactor samples on each measurement day. The instrument was optically aligned daily using 0.5 μm and 1 μm UV mono-disperse fluorescent beads, and the same beads were inserted into each sample as an internal reference to monitor the instrument’s stability (Supplementary Information S4). All raw data are available in the FlowRepository (https://flowrepository.org/; accession no. FR-FCM-Z3MU).

Single-cell data were collected in logarithmically scaled 2D-dot plots of DAPI fluorescence vs. forward scatter (FSC) for cell size-related information. A cell gate that excluded beads and noise was defined, which included 200,000 virtual cells for each measurement. According to apparent cell clusters in the 2D-dot plots of all measured samples (n = 448), a gate-template with 80 gates was defined (G1-G80). The cell gate, the cell numbers per gate of the gate-template, and the measured total cell number (CN) were used to determine both the relative and absolute cell numbers per gate over time (Figs. S8.1-S8.3). In this study, we refer to the cells from a gate as a subcommunity (SC). Data from a total of 35,840 SCs (including 80 SC of the inoculum) were obtained (Supplementary Information S4, Dataset S4). Dominant SCs were defined as having an average proportion of cell numbers higher than 1.25%.

For cell counting, live cells were stained with SYTO^®^9 and counted flow cytometrically with the CyFlow^®^Space (Sysmex Partec GmbH, Görlitz, Germany) by using the True Volumetric Absolute Counting mode and FloMax 2.4 (Sysmex Partec; Supplementary Information S5).

A total of 500,000 cells of each selected SC were sorted, and the cell pellets were stored at -20°C for subsequent DNA isolation (Supplementary Information S4). DNA from both whole-community samples and sorted cells was extracted (Supplementary Information S6). 16S rRNA gene amplicon sequencing was performed by using Pro341F^65^ and Pro805R^66^ primers for the V3-V4 region. To ensure quality of the sequencing run and analysis, a sequencing mock community (ZymoBIOMICS™ Microbial Community Standard, Zymo Research, Irvine, CA, USA) was included in the sequencing project (Fig. S6.1). The community and SC samples were resolved at the genus level (Figs. S9.1, S9.5). All raw data are available in the NCBI Sequence read archive (https://www.ncbi.nlm.nih.gov/bioproject/PRJNA756026/; accession no. PRJNA756026). All tools and scripts that were used for the statistical analyses are provided in Supplementary Information S6, step 6.

Changes in microbiome structures were visualized by dissimilarity analysis (Bray-Curtis index) based on relative cell abundance per SC (Supplementary Information S8, step 1). The temporal variation in community structure was quantified by calculating Canberra distances between vectors of subsequent relative cell numbers in each dominant SC (Supplementary Information S8, step 2). Effluents between local communities L1-L5 and the regional pool R were compared by permutational analysis of variance (PERMANOVA) to test whether L1-L5 alone structured the R community (Supplementary Information S8, step 2).

Intra-and inter-community β-diversity were calculated to quantify temporal and regional variations of the local microbiomes L1-L5 and regional pool R^9^. Community inherent stochastic drift events were highlighted based on intra-community β-diversity values. Changes in α-and γ-diversity values were also documented (Supplementary Information S8, step 3).

To quantify the increasing synchrony in SC emergence in the reactors, community composition variation was partitioned into species replacement and species loss to determine turnover and nestedness using a method that was developed for microbial community flow cytometric data. The turnover β_SIM_ and nestedness β_NES_ components of Sørensen dissimilarity were calculated^30^ (Supplementary Information S8, step 4).

To understand the effect of mass transfer on diversity patterns and synchrony, we calculated the netgrowth rate of both the entire microbiomes (µ^′^, Supplementary Information S10) and every dominant SCs in local microbiomes L1-L5 (µ^′^*SC*_*x*_; Supplementary Information S11). The netgrowth rates were calculated from cell numbers in the inflow, cell numbers lost by the effluent, and the growth of cells in a time interval Δ*t* (d) (Supplementary Information S10: Eqs. S10.1-S10.10; Supplementary Information S11: Eqs. S11.1-S11.5).

To verify synchrony with increasing recycling flow rates *RC*, stability properties^2^ were calculated for the whole communities (Supplementary Information S8), and highlighted for SCs that were promoted by mass transfer (Supplementary Information S12).

### Data evaluation

The absolute and relative cell abundance per SC (G1-G80) within the cell gate was computed with FlowJo v10 (FlowJo LLC, Oregon, USA) and cell number counted with and FloMax v2.4 (Sysmex Partec GmbH, Görlitz, Germany). Unless otherwise noted, all calculations and analyses were performed using RStudio v1.2.1335 (Boston, MA, USA) with R v3.6.3 (R Core Team, 2020). The dissimilarity analysis was supported by the R packages ‘vegan’ v2.5.6^67^ and ‘flowCyBar’^68^ (http://bioconductor.org/packages/flowCyBar/). The correlation analysis was supported by the R packages ‘Hmisc’ v4.4.0^69^. The quantification of stability properties followed the R-script of Liu et al.^2^. Detailed descriptions of the statistical handling are provided in the Supplementary Information: stability test (Supplementary Information S7), community composition analysis (Supplementary Information S8), sequencing-data-based analysis (Supplementary Information S9), netgrowth rates (Supplementary Information S10 and S11, Datasets S10 and S11), and correlation analysis (Supplementary Information S12 and S13), respectively. The Wilcoxon test was conducted to determine significance (*p* < 0.05) of the difference in α-, β-, γ-diversity values, numbers of correlation and stability property values between pairwise phases. The PERMANOVA test was supported by R packages ‘vegan’ v2.5.6^67^. The graphical work was supported by the R packages ‘ggplot2’ v3.3.0^70^.

## Data availability

All flow cytometric raw data are available in the FlowRepository (https://flowrepository.org/; accession no. FR-FCM-Z3MU). All 16S rRNA amplicon sequencing data are available in the NCBI Sequence Read Archive (https://www.ncbi.nlm.nih.gov/sra) under the accession number: PRJNA756026 (https://www.ncbi.nlm.nih.gov/bioproject/PRJNA756026). R-scripts and materials for data analysis and relevant graphical work are available in GitHub (https://github.com/ufzshuangli/mc_masstransfer).

## Acknowledgement of funding

This work was funded by the European Regional Development Funds (EFRE—Europe Funds Saxony, grant 100192205) and the Helmholtz Association, Helmholtz-Centre for Environmental Research – UFZ in the frame of the Integrated Platform Electro-Biorefineries & Biosyntheses. The work was also supported by the Chinese Scholarship Council, the Petroleum Technology Development Fund and the German Academic Exchange Service (ID 57401043).

## Author contribution statement

S.L. performed the loop-designed mass transfer experiment, evaluated abiotic and cytometric data, performed the data analysis, and wrote the manuscript. N.A. measured live cell number, evaluated these data and curated the 16S rRNA amplicon sequencing data. F.S. did the DNA extraction and the 16S rRNA amplicon sequencing. U.N.R. supervised the 16S rRNA amplicon sequencing and revised the manuscript. V.G. helped with ecological theory and wrote the manuscript. S.M. designed the experiment, supervised the work, and wrote the manuscript. Z.L. designed the experiment, performed flow cytometric analysis and cell sorting, evaluated the cytometric data and revised the manuscript. All authors contributed critically to the draft and gave final approval for publication.

## Competing interests

The authors have no conflicts of interest to declare.

## Notes

### Competing Interest Statement

The authors have declared no competing interest.

