## Supplementary Information for "Stabilising microbial communities by looped mass transfer"

### Content

S1: Bioreactor setup ... 3
S2: Experimental setup ... 3
S3: Overview of biotic and abiotic parameters measured in this study ... 5
S4: Flow cytometric analysis of community structure ... 8
Preparation of cell samples ... 8
Step 1: Cell sample fixation ... 8
Step 2: DNA staining with DAPI ... 8
Cytometric analysis ... 9
Step 1: Instrumental setup ... 9
Step 2: Measuring community samples ... 9
Step 3: Creation of the gate template to determine the community structure ... 10
Step 4: Cell sorting ... 10
S5: Cell counting by flow cytometry ... 11
Preparation of cells for the determination of cell numbers ... 11
Step 1: Cell sample dilution ... 11
Step 2: DNA staining with SYTO®9 ... 11
Counting cell number ... 11
Step 1: Instrumental setup ... 11
Step 2: Determination of total cell number ... 12
S6: DNA extraction and 16S rRNA gene amplicon sequencing ... 13
Step 1: DNA extraction ... 13
Step 2: DNA quality testing and library preparation for Illumina MiSeq sequencing ... 14
Step 3: Processing of raw sequencing data, denoising and selection of 16S rRNA gene amplicon sequencing variants ... 14
Step 4: Taxonomic classification ... 16
Step 5: Removing contaminants and setting relative abundance thresholds based on positive and negative controls. ... 16
Step 6: Visualisation of the amplicon sequencing data. ... 17
S7: Stability of communities under the influence of RC$_{10}$, RC$_{50}$ and RC$_{80}$ ... 17
Step 1: Constancy spaces for determining changes in microbial community composition caused by increasing $RC$ ... 18
Step 2: Resistance RS and recovery RV for determining changes in microbial community composition caused by increasing $RC$ ... 19
S8: Analysis of microbial community composition ... 20
Step 1: Visualisation of microbial community composition ... 20
Step 2: Comparison of effluents between local communities L1-L5 and the regional pool R ... 25
Step 3: α-, γ-, β-diversity values of microbial communities L1-L5 and R ... 25
Step 4: Partitioning of β-diversity to reveal turnover and nestedness of SCs ... 28
S9: Taxonomic composition of whole communities and sorted SCs ... 31
Step 1: Taxonomic composition of communities ... 31
Step 2: Taxonomic assignment of sorted SCs ... 36
S10: Calculation of the netgrowth rate μ’ ... 41
Calculation of the recycling rate $RC$ ... 42
Relationship between netgrowth rate μ′ and mass transfer rate $M$ ... 43
Produced cell numbers in local communities L1-L5 ... 45
S11: Quantification of the netgrowth rate per subcommunity $\mu'SCx$ ... 46
Calculation of $\mu'SCx$ on the basis of experimental data ... 48
S12: Effect of the recycling rate $RC$ on the presence of SC ... 50
S13: Relationship between biotic and abiotic parameters ... 52
References ... 53

#### S1: Bioreactor setup

The bioreactors were constructed and handled as described previously[1]. In short, a microbial community from an activated sludge basin of a wastewater treatment plant (Eilenburg, Saxonia, Germany, 51°27′39.4′′N, 12°36′17.5′′E) was pre-cultivated in a medium mixture of peptone medium and synthetic wastewater [v:v = 2%:98%; 0.198 g $L^{-1}$ peptone (from meat), 0.2 g $L^{-1}$ meat extract, 0.219 g $L^{-1}$ yeast extract, 0.1 g $L^{-1}$ glucose, 0.49 g $L^{-1}$ Na-propionate (filtered), 0.0059 g $L^{-1}$ $CaCl_2.2H_2O$, 0.0294 g $L^{-1}$ KCl, 0.06 g $L^{-1}$ NaCl, 0.04 g $L^{-1}$ $K_2HPO_4$, 0.2156 g $L^{-1}$ $KH_2PO_4$ and 0.0196 g $L^{-1}$ $MgSO_4.7H_2O$; chemicals were purchased from: Merck KGaA (Darmstadt, Germany), SERVA Electrophoresis GmbH (Heidelberg, Germany) and Carl Roth GmbH (Karlsruhe, Germany)]. This pre-cultivation was started by mixing 10 mL activated sludge samples with 100 mL medium mixture in a 500 mL Erlenmeyer flask, and the cultivation was carried out on a rotary shaker at 125 rpm and 30°C for 24 h (Incubator Hood TH 25; Edmund Bühler GmbH, Hechingen, Germany). Five bioreactors were then setup in parallel and the same volume of the preculture was used for inoculation for each 1 L bioreactor filled with the medium mixture to a final volume of 800 mL (initial $OD_{600,\lambda=5mm} = 0.057 \pm 0.003$). Effluents of each of the five reactors were collected by a sixth bioreactor, and this reactor was operated in exactly the same way as the other five reactors, only without the addition of fresh medium (Fig. 1).

All six connected reactors were run at 27°C (thermostat with Incubator Hood TH 25) and 350 rpm using a multipoint magnetic stirrer (Thermo Electron LED GmbH, Langenselbold, Germany) and stirrer bars (45 × 8 mm, Labsolute®; Th. Geyer GmbH, Renningen, Germany), and at an aeration rate of 150 mL $min^{-1}$ with compressed sterile filtered ambient air controlled by a rotor gas flowmeter (six measuring channels; Analyt-MTC GmbH, Müllheim, Germany). The fluidic system of the bioreactors was controlled through a set of microprocessor-controlled dispensing pumps IPC-N 12 (Ismatec®; Cole-Parmer GmbH, Wertheim, Germany), and the continuously running mode was maintained over time.

#### S2: Experimental setup

The local communities L1-L5 and the regional pool R together formed a metacommunity (Fig. S2.1). The rates of effluents and influents among the local communities L1-L5 and also for the regional pool R were controlled manually. The flow rate of the total influent (medium plus recycling flow) into each of the local communities L1-L5 was set at a constant 0.4 mL $min^{-1}$, a dilution rate of 0.72 $d^{-1}$ and a hydraulic retention time of 33.3 h. The setup for the regional pool R differed from that of local communities L1-L5. The influent of the regional pool R was the sum of effluents from the five local communities L1-L5 without additional nutrients. This setup caused a fivefold higher dilution rate than for the local communities. Owing to $5D$ (i.e., 3.6 $d^{-1}$), the cells entered and left the regional pool R at a fivefold higher rate than for the five local communities L1-L5.

The six reactors were run for 110 days. The days were sub-grouped into five phases according to increasing recycling flow rates (Table S2.1). The first phase was the Insular I phase, in which no exchange with the other reactors was allowed. The reactor serving as the regional pool R was run as a sink for local community effluents starting at day 9, which were not recycled back to the local communities L1-L5. The second phase $RC_{10}$ (i.e. recycling rate $RC$ 10%) started at day 26 in which the regional pool R and the local communities L1-L5 were interconnected. The third phase $RC_{50}$ (i.e. recycling rate $RC$ 50%) was started at day 47 and the fourth phase $RC_{80}$ (i.e. recycling rate $RC$ 80%) at day 64. In phases 2-4, the inflow from the regional pool R, increasing from 0.04, 0.2 and 0.32 mL $min^{-1}$, was compensated for by decreasing the amounts of medium. Therefore, the medium flow rates were lowered to 0.36, 0.2 and 0.08 mL $min^{-1}$ (Table S2.1). To maintain the nutrient load rate for the local communities L1-L5 under the recycling flow rate conditions, the nutrients in the medium were concentrated 1.1 times, 2 times and 5 times, accordingly. In the fifth Insular II phase starting at day 89, all recycling from the regional pool R to local communities L1-L5 was stopped and conditions returned back to how they were in the first phase. Within each of the five phases, the first 7 days were defined as an adaptation period in which the medium volume of the reactors was exchanged five times. Afterwards, balanced growth conditions were assumed. A total of 448 samples were collected from the six bioreactors in 110 days, with 76 samples from each of the local communities L1-L5 and 68 samples from the regional pool R. The number of samples, recycling flow rate settings and time intervals per phase are summarised in Table S2.1.

**Table S2.1** Summary of sample numbers, recycling rate ($RC$) setting and time intervals per phase

| phase | number of samples per reactor | medium flow rate (mL $min^{-1}$) | recycling flow rate (mL $min^{-1}$) | medium factor | adaptation period (d) | balanced period (d) |
|---|---|---|---|---|---|---|
| Insular I | 18 (10 for R) | 0.4 | 0 | 1 | 0-8 | 8-26 |
| $RC_{10}$ | 14 | 0.36 | 0.04 | 1.1 | 26-33 | 33-47 |
| $RC_{50}$ | 13 | 0.2 | 0.2 | 2 | 47-54 | 54-64 |
| $RC_{80}$ | 15 | 0.08 | 0.32 | 5 | 64-71 | 71-89 |
| Insular II | 16 | 0.4 | 0 | 1 | 89-96 | 96-110 |

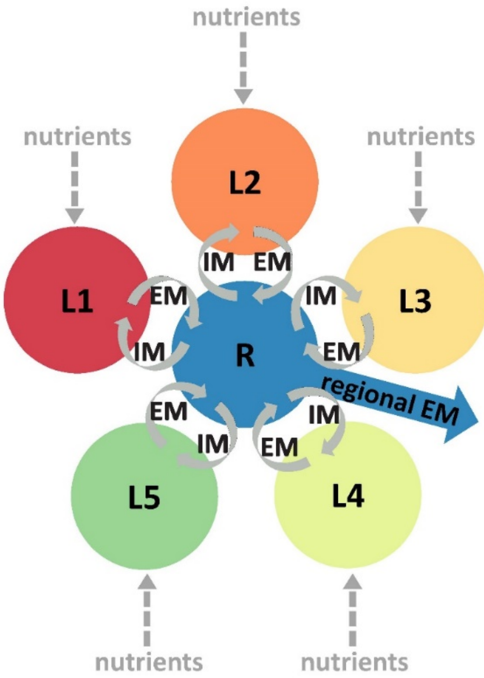


**Figure S2.1** Conceptual setup of the metacommunity investigated in this study. Local communities L1-L5 assembled in identical localities, and all were connected to a community in the regional pool R. Microbial immigration (IM) and emigration (EM) of local communities occurred between each of the local communities and the regional pool R. In addition, cells were emigrated via the regional pool R.

#### S3: Overview of biotic and abiotic parameters measured in this study

For each sample, 7 mL of cell culture was taken, of which 0.2 mL was used for counting total cell number (CN), 1 mL for DNA extraction (centrifuged at 21,000×g for 10 min and 4°C, and pellet stored at -20°C), and 5.8 mL for the measurements of optical density (OD), pH and electrical conductivity (EC), as well as for the flow cytometric analysis of the cells. In addition, 5 mL was collected two to five times a week per reactor to analyse the chemical oxygen demand of the supernatant (CODs) and of the total sample (CODt), ammonia nitrogen (NH4, supernatant) and total phosphate, calculated as phosphor (PHOt, total sample). The dry weight (DW) was measured once a week using a 20 mL sample.

The optical density ($OD_{600, \lambda=5\ mm}$, Ultraspec 1100pro; Amersham Biosciences, Little Chalfont, UK), pH (EL 20; Mettler-Toledo, Greifensee, Switzerland) and electrical conductivity (EC, inolab Cond7110; WTW, Germany) were measured daily in all reactors. Additionally, COD, NH4 and PHOt were measured daily in the first 7 days (i.e., adaptation period) and once every 3 or 4 days during the balanced period per phase. CODs and CODt (DIN ISO 15705:2002) and NH4 (DIN 38406-E5) were analysed with NANOCOLOR® test tubes (Macherey-Nagel GmbH, Düren, Germany) following the manufacturer's instructions. CODb (chemical oxygen demand of biomass) was calculated by subtracting CODs from CODt. PHOt was measured by phosphomolybdate blue spectrophotometry (DIN 38405-D11). Briefly, after a pre-heating treatment (120°C, 30 min), 500 µL diluted supernatant was treated with 800 µL reagent [stock solution: 125 mL $H_2O$, 25 mL 9N $H_2SO_4$, 25 mL $Mo_7O_{24}^{-6}$ solution (1.65 g $(NH_4)_6Mo_7O_{24}\cdot 4H_2O$ in 25 mL $H_2O$), and 25 mL Fe (II) solution (3.89 g $(NH_4)_2Fe(SO_4)_2\cdot 6H_2O$ in 25 mL $H_2O$)] for 10 minutes. The dry weight (DW) of the biomass was measured once a week during the balanced growth conditions of each phase. Briefly, the cell solution was centrifuged at 5,000×g for 10 min at 4°C (Centrifuge 5804R; Eppendorf, Hamburg, Germany). The supernatant was discarded and the residual pellet was transferred to a 2 mL tube and centrifuged at 20,000×g for 10 min at 4°C (Heraeus Fresco 21 centrifuge; Thermo Scientific, Langenselbold, Germany). Finally, the pellet was dried at 50°C for 4 days. In addition to CODb and DW, biomass was also evaluated by counting the total number of cells $mL^{-1}$ (CN) by flow cytometry (Supplementary Information 5). All other supernatants measured in this study were obtained by centrifuging the samples at 3,200×g for 10 min and 4°C. The OD, pH, EC and PHOt measurements were performed in triplicate, while COD and NH4 were measured in duplicates. A graphical overview of all parameters is shown in Fig. S3.1. All data of the parameters are listed in Dataset S3.

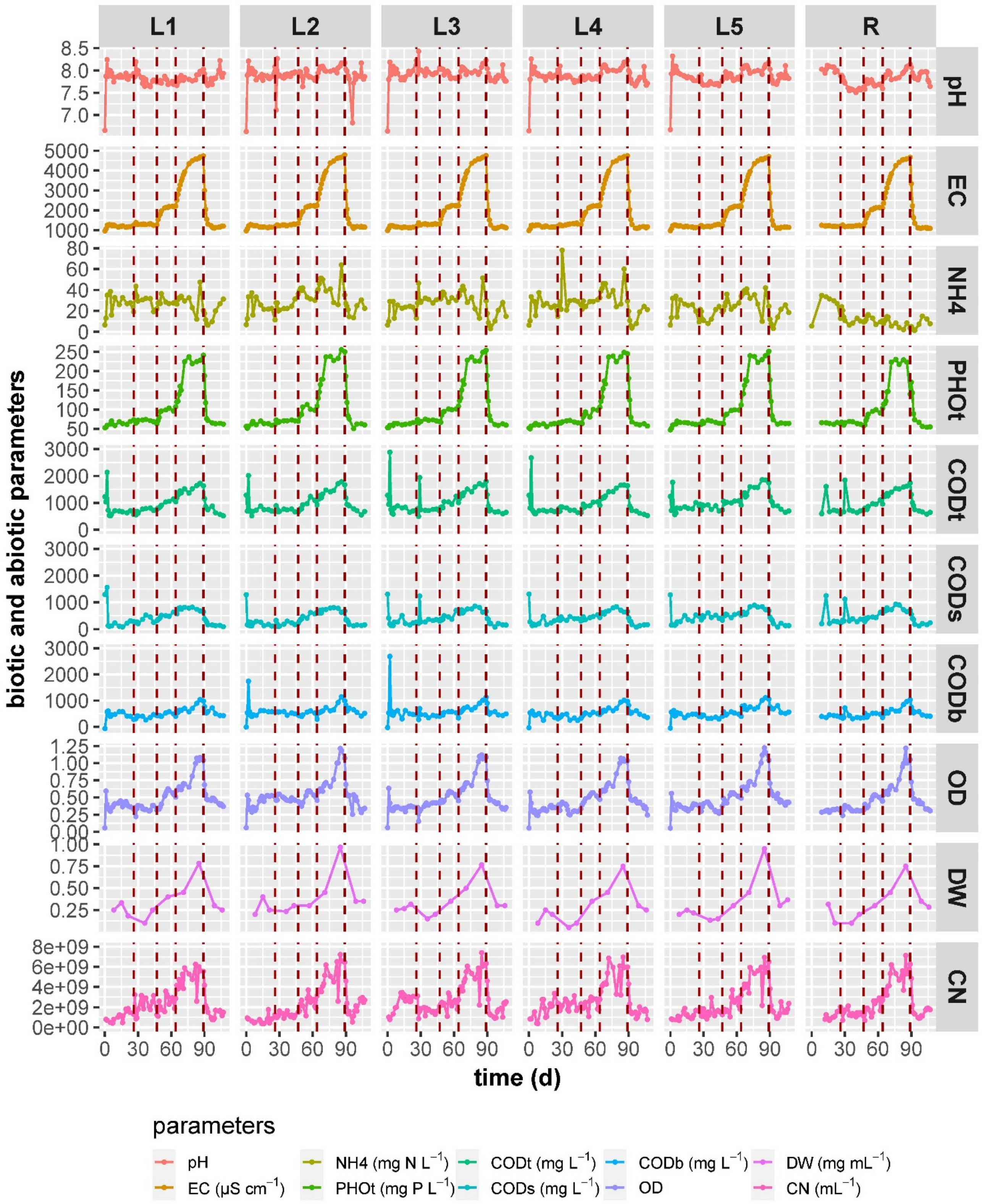


**Figure S3.1** Overview of the bulk biotic and abiotic parameters measured for the local communities L1-L5 and the regional pool R. Parameters were pH, EC (electrical conductivity, μS cm$^{-1}$), NH4 (ammonium, mg N L$^{-1}$), PHOt (phosphate of total sample, mg P L$^{-1}$), CODt (chemical oxygen demand of total sample, mg L$^{-}$

$^{1}$), CODs (chemical oxygen demand of supernatant, mg $L^{-1}$), CODb (chemical oxygen demand of biomass, mg $L^{-1}$), $OD_{600,\lambda=5mm}$ (optical density), DW (dry weight, g $L^{-1}$) and CN (cell number, $mL^{-1}$). The dashed red lines indicate the times at which the next phase begins.

With the increase of $RC_{10}$ to $RC_{80}$ the values for the abiotic parameter EC and phosphate also increased, while the values for ammonium and pH remained constant in local communities L1-L5. The biotic parameter showed increased values for biomass (CODb, OD, dry weight and cell number; comparisons between successive phases, Wilcoxon test: $p \leqslant 0.01$). The regional pool R showed similar trends, with the exception of ammonium, which decreased. In all reactors, the highest biomass values were found in $RC_{80}$ and the lowest in Insular I phase and Insular II phase (Fig. S3.1, Dataset S3).

#### S4: Flow cytometric analysis of community structure

##### Preparation of cell samples

###### Step 1: Cell sample fixation

For fixation, 2 × 2.5 mL samples were taken from each reactor and placed in glass tubes. Supernatants were removed after centrifugation at 3,200×g for 10 min at 4°C. For every tube, the cells in the pellet were suspended in 2 mL paraformaldehyde solution [PFA, 2% in phosphate-buffered saline (PBS, 6 mM $Na_2HPO_4$, 1.8 mM $NaH_2PO_4$, 145 mM NaCl, pH 7)] and incubated for 30 min at room temperature (RT). Afterwards, the cells were centrifuged again (3,200×g, 10 min, 4°C), resuspended in 4 mL 70% ethanol, and then stored at -20°C.

###### Step 2: DNA staining with DAPI

An aliquot of the fixed sample was taken into a glass tube and washed twice with PBS (3,200*g, 10 min, 4°C), and then adjusted with PBS to an $OD_{700,\lambda=5\,mm}$ of 0.035. Two mL of the adjusted cell solution was centrifuged (3,200×g, 10 min, 4°C), and the pellet was resuspended in 1 mL solution A (0.11 M citric acid and 4.1 mM Tween 20 in bidistilled water) and incubated at RT for 20 min, the first 10 min in an ultrasonication bath (35 kHz; Merck Eurolab, Darmstadt, Germany). Solution A was discarded after centrifugation (3,200×g, 10 min, 4°C). The cells were resuspended in 2 mL solution B [0.24 µM DAPI (4',6-diamidino-2-phenylindole, Lot. 118M4025V; Sigma-Aldrich, St. Louis, MO, USA) in phosphate buffer (289 mM $Na_2HPO_4$ and 128 mM $NaH_2PO_4$ in bidistilled water)] and incubated overnight at RT and in the dark before flow cytometric measurement.

#### Cytometric analysis

##### Step 1: Instrumental setup

The cells were flow cytometrically measured in a MoFlo Legacy Cell Sorter using the software Summit v4.3 (Beckman Coulter, Brea, CA, USA). The instrument was equipped with a 488 nm argon laser (400 mW; Coherent, Santa Clara, CA, USA) and a 355 nm UV laser (150 mW, Xcyte CY-355-150; Lumentum, Milpitas, CA, USA). The 488 nm laser light was used for detection of the forward scatter (FSC, 488/10 nm band pass) and the side scatter (SSC, 488/10 nm band pass, trigger signal). The DAPI fluorescence was measured at signal channel FL4 (450/65 nm band pass) after excitation with the UV laser beam.

The fluidic system was run at constant sheath pressure of 56.0 psi with a 70 µm nozzle. The sample pressure was adjusted within the range of 55.8-56.2 psi, in order to obtain a stable event measurement of around 3,500 events per second. The sheath fluid was composed of 10-fold sheath buffer (19 mM $KH_2PO_4$, 38 mM KCl, 166 mM $Na_2HPO_4$ and 1.39 M NaCl with 0.13 µm filtrated Millipore water) and further diluted with filtrated Millipore water to a 0.2-fold working solution (for cell sorting: 0.5-fold working solution).

For the daily optical calibration of the cytometer in the linear range, 1 µm blue fluorescent FluoSpheres (F-8815; Molecular Probes, Eugene, OR, USA) and 2 µm yellow-green fluorescent FluoSpheres (F8827; ThermoFisher Scientific, Waltham, MA, USA) were used. For calibration in the logarithmic range, 0.5 µm and 1.0 µm UV Fluoresbrite Microspheres (18339 and 17458, respectively; Polysciences, Warrington, PA, USA) were used.

To ensure the reliability and comparability of the cell fixation and staining procedures, a microbial cytometric mock community (mCMC)[2] was used each day. The cells of the mCMC were handled identically to the staining protocol in the section S4 (steps 1 & 2: Preparation of cell samples). The use of the mCMC guaranteed a high resolution by ensuring optimal optical settings of the flow cytometer and comparable cytometric measurements of bioreactor samples, even if they were measured over months.

##### Step 2: Measuring community samples

Prior to measurement, DAPI-stained cells were filtered to remove larger particles by using a nylon filter (CellTrics® 50 µm; Sysmex Partec GmbH, Görlitz, Germany) and were spiked with 0.5 µm and 1 µm UV Fluoresbrite Microspheres (18339 and 17458, respectively; Polysciences). The microspheres served as internal standards to monitor instrument stability and to allow the correct comparison of samples (Fig. S4.1a). Cell data were collected in logarithmically scaled 2D-dot plots according to DAPI fluorescence for DNA content and forward scatter (FSC) for cell sizerelated information. A cell gate was defined, which comprised 200,000 virtual cells for each measurement (cell gate; Fig. S4.1a).

##### Step 3: Creation of the gate-template to determine the community structure

According to the measured samples, apparent cell clusters in 2D-dot plots (FSC and DAPI fluorescence) were gated sample per sample[3] and all defined gates were combined together to create the  gate template. In this study, the gate template included 80 gates (G1 to G80; Fig. S4.1b), and we defined the cell population in each of the gates as a subcommunity (SC). The relative cell abundance per SC (G1-G80) within the cell gate was computed using FlowJo™ v10 (FlowJo LLC, Ashland, OR, USA) automatically. A list of the data on relative cell abundance per gate per sample is given in the Dataset S4.

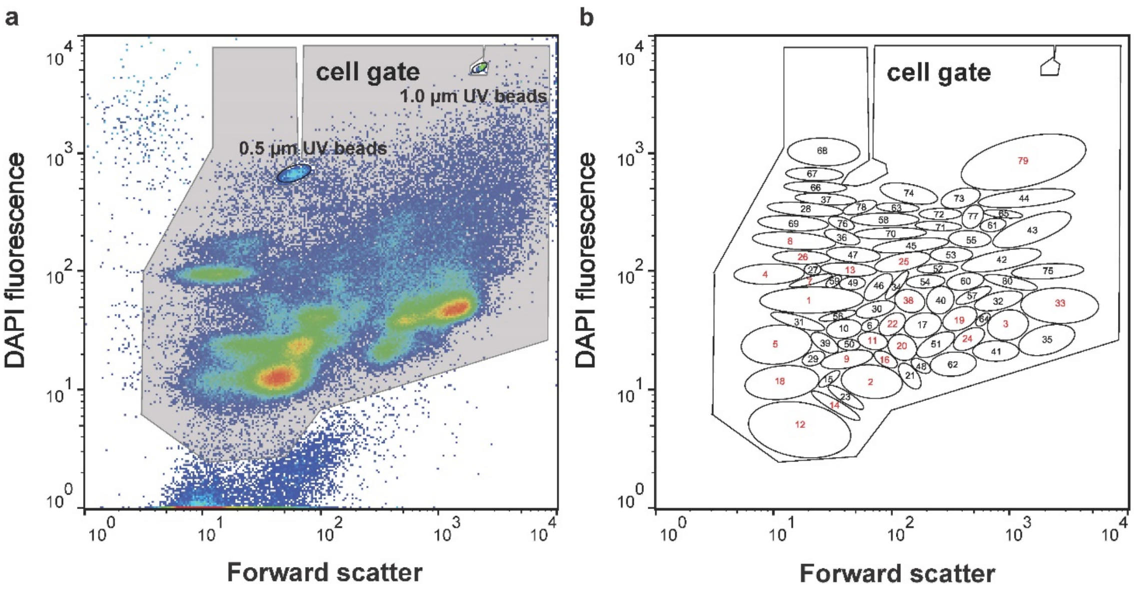


**Figure S4.1** Gating strategy. First, a cell gate was defined for the measurement of 200,000 cells. Calibration beads were excluded. Afterwards, the gate template was created by defining 80 gates (i.e., G1-G80), based on forward scatter (Forward scatter) and DAPI fluorescence (DAPI fluorescence). The cell numbers per gate and sample were used for evaluations in further data evaluation pipelines. Sorted gates are marked by red numbers.

##### Step 4: Cell sorting

To determine the taxonomic affiliation of selected SCs, cell sorting was performed. The cell sorting procedure was carried out in accordance with the work of Cichocki et al.[2]. Briefly, the positions of gates to be sorted were defined and assigned using the Summit software (V4.3; Beckman Coulter,

Brea, CA, USA). The cell sorting was performed using the four-way-sort option and the ‘1.0 Drop Pure’ sort mode. A total of 500,000 cells of each selected gate were sorted into a 1.5 mL Eppendorf tube at an event rate of not more than 1,500 events per second. Sorted cells were harvested from the sheath buffer by centrifugation (20,000×g, 6°C, 25 min), and the cell pellets were stored at -20°C for subsequent DNA isolation.

#### S5: Cell counting by flow cytometry

##### Preparation of cells for the determination of cell numbers

###### Step 1: Cell sample dilution

Live cells (0.2 mL) were sampled from the bioreactors and directly diluted in three standardised steps (all in all 1/500-fold) to about $10^7$ cells per mL using 0.85% saline solution.

###### Step 2: DNA staining with SYTO®9

SYTO®9 (Lot. 2088729; ThermoFisher Scientific, Eugene, OR, USA) is a cell-permeant nucleic acid stain, which was used in this study to stain fresh cell samples to distinguish cells from medium particles. A 35 µM stock solution of SYTO® 9 was prepared daily and stored on ice. The final cell solution contained 950 µL diluted cell sample and 50 µL 35 µM SYTO®9, with a final concentration of 1.75 µM SYTO®9. The cell solution was mixed and incubated for 15 min at RT before cell counting. The diluted and stained cell samples were measured on the same day.

##### Counting cell numbers

###### Step 1: Instrumental setup

The cell counting was performed with the flow cytometer CyFlow®Space (Sysmex Partec GmbH, Görlitz, Germany) using the True Volumetric Absolute Counting mode, which counts cells in a fixed volume (0.2 mL). This device was equipped with a 488 nm argon laser (50mW; Sapphire, Coherent, Santa Clara, CA, USA). The fluorescence of the stained cells was measured using the filters 536/40 nm band pass for green fluorescence and 610/30 nm band pass for red fluorescence. For the daily optical calibration of the flow cytometer in the linear range, 0.5 µm yellow-green fluorescent FluoSpheres (F8827; ThermoFisher Scientific, Waltham, MA, USA) and 1.0 µm yellow-green fluorescent FluoSpheres (F13081; ThermoFisher Scientific, Waltham, MA, USA) were used. These latter FluoSpheres were also added to each sample to ensure comparability of measurements.

##### Step 2: Determination of total cell number

Before determining the total cell number (CN, $mL^{-1}$), the cell suspension was filtered to remove larger particles by using a nylon filter (CellTrics® 50 µm; Sysmex Partec GmbH, Görlitz, Germany). The measuring rate was adjusted to below 1500 events per second. Dependent on the cell concentration, 100-200 µL of the cell solution was added to 1-1.1 mL Millipore water to maintain the measuring rate. A cell gate was defined on the basis of SYTO®9 green and red fluorescence (Fig. S5.1) to differentiate the cells from instrumental and background noise. Total cell numbers were counted automatically using the software FloMax (V2.4; Sysmex Partec GmbH, Germany). Based on the cell numbers in diluted samples and the actual dilution factor, the CNs in local communities L1-L5 and the regional pool R were calculated. In total, 421 values (71 per local community and 66 per regional pool) were measured, as shown in Dataset S3 and visualised in Fig. S5.2. In Table S5.1, the mean values and the standard deviation of cell numbers per phase (balanced period) are shown for the local communities L1-L5 and the regional pool R, respectively. The CN per SC was calculated by multiplying relative cell numbers per SC (Dataset S4) by cell number per community (Dataset S5).

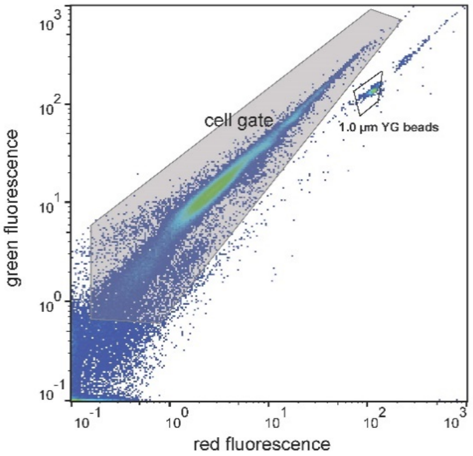


**Figure S5.1**. Cell gate for cell counting. The cell gate was set apart from instrumental noise, background noise and calibration beads.

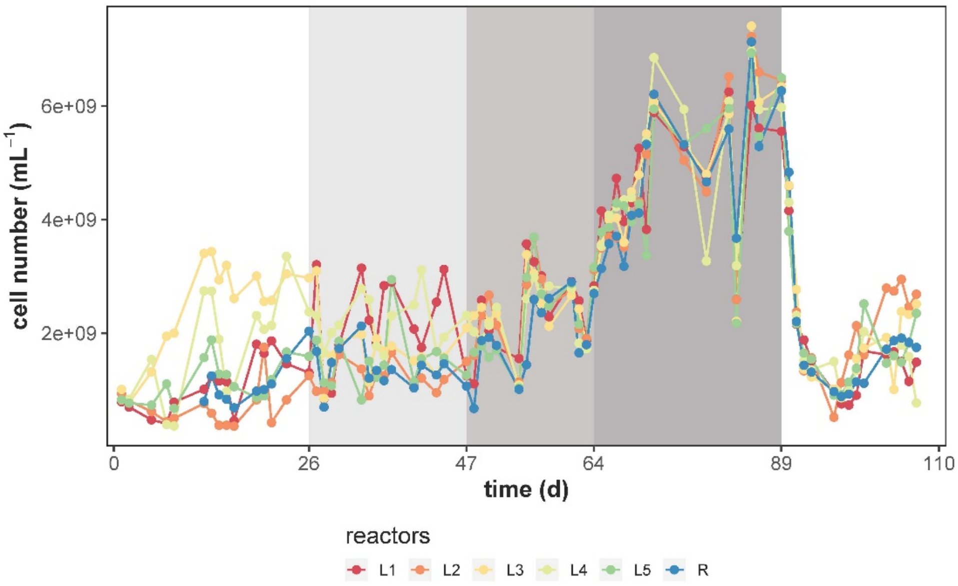


**Figure S5.2** Analysis of cell numbers (CN, $mL^{-1}$) in the local communities L1-L5 and the regional pool R. The shaded areas represent different phases with changed $RC$.

**Table S5.1** Summary of mean values and standard deviation (sd) of cell numbers ($\times 10^9$ cells $mL^{-1}$) per balanced period and phase of the local communities (L1-L5) and the regional pool (R).

| reactor | Insular I | | $RC_{10}$ | | $RC_{50}$ | | $RC_{80}$ | | Insular II | |
|---|---|---|---|---|---|---|---|---|---|---|
| | mean | sd | mean | sd | mean | sd | mean | sd | mean | sd |
| local communities (L1-L5) | 1.60 | 0.93 | 1.82 | 0.65 | 2.46 | 0.71 | 5.29 | 1.39 | 1.58 | 0.62 |
| regional pool (R) | 1.02 | 0.26 | 1.38 | 0.31 | 2.05 | 0.67 | 5.40 | 1.02 | 1.41 | 0.44 |

#### S6: DNA extraction and 16S rRNA gene amplicon sequencing

The 16S rRNA amplicon gene sequencing analysis was performed as follows:

##### Step 1: DNA extraction

The DNA was extracted in accordance with a protocol from Cichocki et al.[2]. In short, 70 µL of Chelex 100 solution [Chelex 100 sodium form, Sigma-Aldrich, CAS no. 11139-85-8, preparation of 10% (wt/vol) with molecular-biology-grade water] was added to each frozen cell pellet from whole communities or sorted cells of SCs (Supplementary Information 4, Cytometric analysis, step 4). Whole-community pellets were thawed, $OD_{700,\lambda=5mm}$ adjusted to 0.01 with sterile PBS and centrifuged at 20,000×g for 25 min at 4°C prior to the Chelex addition. Negative controls (14 controls, sheath buffer and 300 µL of Chelex solution) and a positive control (ZymoBIOMICS™

Microbial Community Standard; Zymo Research Europe GmbH, Freiburg, Germany) were included in the analysis. Each of the solutions was vortexed for 10 s, incubated at 95°C for 45 min and centrifuged at 7,000×g at 4°C for 5 min. Fifty µL of the supernatant, which contained the extracted DNA, was transferred to a new pre-chilled tube. Samples were stored at -20°C before further use.

##### Step 2: DNA quality testing and library preparation for Illumina MiSeq sequencing

DNA yield was determined using Quibit 3.0 (Thermo Fisher Scientific, Waltham, MA, USA). DNA extracts with more than 0.1 ng of DNA per µL were amplified for 25 PCR cycles and those with less than 0.1 ng per µL were amplified for 35 PCR cycles. Table S6.1 indicates the number of cycles used for PCR amplification of the different samples. 16S rRNA gene amplicon sequencing was performed on the V3-V4 region of the 16S rRNA gene region using the primers Pro341F 5'-CCTACGGGNBGCASCAG-3'[4] and Pro805R 5'-GACTACNVGGGTATCTAATCC-3'[5]. These primers and the barcoded primers used for the library were synthesised by Eurofins (Eurofins Scientific, Luxembourg City, Luxembourg). PCR amplification was performed in accordance with the procedure of Liu et al.[1] and checked for the presence of single band PCR products by gel electrophoresis (1.5% agarose). For each PCR batch, a negative control without any DNA was amplified for up to 35 cycles and checked by gel electrophoresis (1.5% agarose) to ensure that no contamination was present. In the absence of contamination, the samples were purified, quantified and equimolarly pooled in accordance with the work of Liu et al.[1]. The pooled amplicon samples were sequenced with MiSeq (Illumina®, San Diego, CA, USA).

##### Step 3: Processing of raw sequencing data, denoising and selection of 16S rRNA gene amplicon sequencing variants

The raw sequence reads from Illumina Miseq were checked and separated according to their number of PCR cycles (Table S6.1) by using manifest files as indicated by the QIIME2 v2020.2, following the instructions of the developers[6]. The paired-end sequence reads fastq files were demultiplexed using q2-demux. Subsequently, the primers were trimmed and low-quality reads were removed as defined by QIIME2[6]. Denoising and selection of amplicon sequencing variants were performed separately for samples amplified for 25 or 35 PCR cycles using DADA2, in accordance with the instructions of the developers[7].

**Table S6.1**. Metadata of different libraries from whole communities and sorted gates amplified based on 25 and 35 PCR cycles. Run: the sequencing round in which the group of samples was sequenced; Type:

the different sorted gates (G) and the whole-community (WC) samples; Identity: which samples were sorted or kept as a whole-community; PCR cycles: the number of cycles for which the different samples were amplified during PCR.

| sample ID | reactor | days | run | type | identity | PCR-Cycles | sample ID | reactor | days | run | type | identity | PCR-Cycles |
|---|---|---|---|---|---|---|---|---|---|---|---|---|---|
| Samp01 | R | 85 | Run1 | G5 | sorted | 35 | Samp80 | R | 97 | Run2 | WC | Community-88 | 25 |
| Samp02 | R | 85 | Run1 | G12 | sorted | 35 | Samp81 | L1 | 100 | Run2 | WC | Community-89 | 25 |
| Samp03 | L2 | 86 | Run1 | G18 | sorted | 35 | Samp82 | L3 | 100 | Run2 | WC | Community-90 | 25 |
| Samp04 | L3 | 61 | Run1 | G7 | sorted | 35 | Samp83 | L1 | 107 | Run2 | WC | Community-91 | 25 |
| Samp05 | L3 | 61 | Run1 | G12 | sorted | 35 | Samp84 | L2 | 107 | Run2 | WC | Community-92 | 25 |
| Samp06 | L3 | 63 | Run1 | G8 | sorted | 35 | Samp85 | L3 | 107 | Run2 | WC | Community-93 | 25 |
| Samp07 | L1 | 100 | Run1 | G7 | sorted | 35 | Samp86 | L4 | 107 | Run2 | WC | Community-94 | 25 |
| Samp08 | L2 | 47 | Run1 | G16 | sorted | 35 | Samp87 | L5 | 107 | Run2 | WC | Community-95 | 25 |
| Samp09 | L2 | 47 | Run1 | G22 | sorted | 35 | Samp88 | L2 | 86 | Run1 | G25 | sorted | 35 |
| Samp10 | L1 | 34 | Run1 | G4 | sorted | 35 | Samp89 | L3 | 61 | Run1 | G18 | sorted | 35 |
| Samp11 | L1 | 99 | Run1 | G8 | sorted | 35 | Samp90 | L3 | 63 | Run1 | G4 | sorted | 35 |
| Samp12 | L5 | 44 | Run1 | G7 | sorted | 35 | Samp91 | L3 | 63 | Run1 | G9 | sorted | 35 |
| Samp13 | L3 | 100 | Run1 | G5 | sorted | 35 | Samp99 | L1 | 0 | Run1 | WC | Community_01 | 35 |
| Samp14 | L1 | 71 | Run1 | G4 | sorted | 35 | Samp100 | L1 | 1 | Run1 | WC | Community_02 | 25 |
| Samp15 | L2 | 44 | Run1 | G1 | sorted | 35 | Samp101 | L2 | 1 | Run1 | WC | Community_03 | 25 |
| Samp16 | L2 | 44 | Run1 | G24 | sorted | 35 | Samp102 | L3 | 1 | Run1 | WC | Community_04 | 25 |
| Samp17 | L2 | 44 | Run1 | G26 | sorted | 35 | Samp103 | L4 | 1 | Run1 | WC | Community_05 | 25 |
| Samp18 | L4 | 86 | Run1 | G13 | sorted | 35 | Samp104 | L5 | 1 | Run1 | WC | Community_06 | 25 |
| Samp19 | L5 | 71 | Run1 | G2 | sorted | 35 | Samp105 | L1 | 8 | Run1 | WC | Community_07 | 25 |
| Samp20 | L5 | 71 | Run1 | G4 | sorted | 35 | Samp106 | L2 | 8 | Run1 | WC | Community_08 | 25 |
| Samp21 | L1 | 99 | Run1 | G20 | sorted | 35 | Samp107 | L3 | 8 | Run1 | WC | Community_09 | 25 |
| Samp22 | R | 85 | Run1 | G14 | sorted | 35 | Samp108 | L4 | 8 | Run1 | WC | Community_10 | 25 |
| Samp23 | L1 | 85 | Run1 | G5 | sorted | 35 | Samp109 | L5 | 8 | Run1 | WC | Community_11 | 25 |
| Samp24 | L1 | 85 | Run1 | G12 | sorted | 35 | Samp110 | R | 9 | Run1 | WC | Community_12 | 25 |
| Samp25 | L1 | 85 | Run1 | G14 | sorted | 35 | Samp111 | L1 | 26 | Run1 | WC | Community_13 | 25 |
| Samp26 | L3 | 85 | Run1 | G5 | sorted | 35 | Samp112 | L2 | 26 | Run1 | WC | Community_14 | 25 |
| Samp27 | L3 | 85 | Run1 | G12 | sorted | 35 | Samp113 | L3 | 26 | Run1 | WC | Community_15 | 25 |
| Samp28 | L3 | 85 | Run1 | G14 | sorted | 35 | Samp114 | L4 | 26 | Run1 | WC | Community_16 | 25 |
| Samp29 | L2 | 86 | Run1 | G9 | sorted | 35 | Samp115 | L5 | 26 | Run1 | WC | Community_17 | 25 |
| Samp34 | L3 | 61 | Run1 | G5 | sorted | 35 | Samp116 | R | 26 | Run1 | WC | Community_18 | 25 |
| Samp35 | L3 | 99 | Run1 | G12 | sorted | 35 | Samp117 | L1 | 27 | Run1 | WC | Community_19 | 25 |
| Samp36 | L1 | 34 | Run1 | G19 | sorted | 35 | Samp118 | L2 | 27 | Run1 | WC | Community_20 | 25 |
| Samp37 | R | 85 | Run1 | G33 | sorted | 35 | Samp119 | L3 | 27 | Run1 | WC | Community_21 | 25 |
| Samp38 | L1 | 85 | Run1 | G33 | sorted | 35 | Samp120 | L4 | 27 | Run1 | WC | Community_22 | 25 |
| Samp39 | L3 | 85 | Run1 | G33 | sorted | 35 | Samp121 | L5 | 27 | Run1 | WC | Community_23 | 25 |
| Samp40 | L2 | 86 | Run1 | G1 | sorted | 35 | Samp122 | R | 27 | Run1 | WC | Community_24 | 25 |
| Samp41 | L5 | 55 | Run2 | WC | Community_49 | 25 | Samp123 | L1 | 34 | Run1 | WC | Community_25 | 25 |
| Samp42 | R | 55 | Run2 | WC | Community_50 | 25 | Samp124 | L2 | 34 | Run1 | WC | Community_26 | 25 |
| Samp43 | L3 | 61 | Run2 | WC | Community_51 | 25 | Samp125 | L3 | 34 | Run1 | WC | Community_27 | 25 |
| Samp44 | L5 | 61 | Run2 | WC | Community_52 | 25 | Samp126 | L4 | 34 | Run1 | WC | Community_28 | 25 |
| Samp45 | L1 | 64 | Run2 | WC | Community-53 | 25 | Samp127 | L5 | 34 | Run1 | WC | Community_29 | 25 |
| Samp46 | L2 | 64 | Run2 | WC | Community-54 | 25 | Samp128 | R | 34 | Run1 | WC | Community_30 | 25 |
| Samp47 | L3 | 64 | Run2 | WC | Community-55 | 25 | Samp129 | L2 | 44 | Run1 | WC | Community_31 | 25 |
| Samp48 | L4 | 64 | Run2 | WC | Community-56 | 25 | Samp130 | L5 | 44 | Run1 | WC | Community_32 | 25 |
| Samp49 | L5 | 64 | Run2 | WC | Community-57 | 25 | Samp131 | L1 | 47 | Run1 | WC | Community_33 | 25 |
| Samp50 | R | 64 | Run2 | WC | Community-58 | 25 | Samp132 | L2 | 47 | Run1 | WC | Community_34 | 25 |
| Samp51 | L1 | 65 | Run2 | WC | Community-59 | 25 | Samp133 | L3 | 47 | Run1 | WC | Community_35 | 25 |
| Samp52 | L2 | 65 | Run2 | WC | Community-60 | 25 | Samp134 | L4 | 47 | Run1 | WC | Community_36 | 25 |
| Samp53 | L3 | 65 | Run2 | WC | Community-61 | 25 | Samp135 | L5 | 47 | Run1 | WC | Community_37 | 25 |
| Samp54 | L4 | 65 | Run2 | WC | Community-62 | 25 | Samp136 | R | 47 | Run1 | WC | Community_38 | 25 |
| Samp55 | L5 | 65 | Run2 | WC | Community-63 | 25 | Samp137 | L1 | 48 | Run1 | WC | Community_39 | 25 |
| Samp56 | R | 65 | Run2 | WC | Community-64 | 25 | Samp138 | L2 | 48 | Run1 | WC | Community_40 | 25 |
| Samp57 | L1 | 72 | Run2 | WC | Community-65 | 25 | Samp139 | L3 | 48 | Run1 | WC | Community_41 | 25 |
| Samp58 | L2 | 72 | Run2 | WC | Community-66 | 25 | Samp140 | L4 | 48 | Run1 | WC | Community_42 | 25 |
| Samp59 | L3 | 72 | Run2 | WC | Community-67 | 25 | Samp141 | L5 | 48 | Run1 | WC | Community_43 | 25 |
| Samp60 | L4 | 72 | Run2 | WC | Community-68 | 25 | Samp142 | R | 48 | Run1 | WC | Community_44 | 25 |
| Samp61 | L5 | 72 | Run2 | WC | Community-69 | 25 | Samp143 | L1 | 55 | Run1 | WC | Community_45 | 25 |
| Samp62 | R | 72 | Run2 | WC | Community-70 | 25 | Samp144 | L2 | 55 | Run1 | WC | Community_46 | 25 |
| Samp63 | L1 | 89 | Run2 | WC | Community-71 | 25 | Samp145 | L3 | 55 | Run1 | WC | Community_47 | 25 |
| Samp64 | L2 | 89 | Run2 | WC | Community-72 | 25 | Samp146 | L4 | 55 | Run1 | WC | Community_48 | 25 |
| Samp65 | L3 | 89 | Run2 | WC | Community-73 | 25 | Samp154 | L1 | 99 | Run2 | G38 | sorted | 35 |
| Samp66 | L4 | 89 | Run2 | WC | Community-74 | 25 | Samp155 | L1 | 63 | Run2 | G2 | sorted | 35 |
| Samp67 | L5 | 89 | Run2 | WC | Community-75 | 25 | Samp156 | L1 | 63 | Run2 | G3 | sorted | 35 |
| Samp68 | R | 89 | Run2 | WC | Community-76 | 25 | Samp157 | L1 | 63 | Run2 | G11 | sorted | 35 |
| Samp69 | L1 | 90 | Run2 | WC | Community-77 | 25 | Samp158 | L1 | 100 | Run2 | G3 | sorted | 35 |
| Samp70 | L2 | 90 | Run2 | WC | Community-78 | 25 | Samp159 | L1 | 100 | Run2 | G4 | sorted | 35 |
| Samp71 | L3 | 90 | Run2 | WC | Community-79 | 25 | Samp160 | L1 | 100 | Run2 | G24 | sorted | 35 |
| Samp72 | L4 | 90 | Run2 | WC | Community-80 | 25 | Samp161 | L2 | 47 | Run2 | G33 | sorted | 35 |
| Samp73 | L5 | 90 | Run2 | WC | Community-81 | 25 | Samp162 | L2 | 47 | Run2 | G79 | sorted | 35 |
| Samp74 | R | 90 | Run2 | WC | Community-82 | 25 | Samp163 | L2 | 26 | Run2 | G33 | sorted | 35 |
| Samp75 | L1 | 97 | Run2 | WC | Community-83 | 25 | Samp164 | L5 | 47 | Run2 | G3 | sorted | 35 |
| Samp76 | L2 | 97 | Run2 | WC | Community-84 | 25 | Samp165 | L3 | 100 | Run2 | G18 | sorted | 35 |
| Samp77 | L3 | 97 | Run2 | WC | Community-85 | 25 | Samp166 | L3 | 58 | Run2 | G19 | sorted | 35 |
| Samp78 | L4 | 97 | Run2 | WC | Community-86 | 25 | Samp167 | L3 | 58 | Run2 | G24 | sorted | 35 |
| Samp79 | L5 | 97 | Run2 | WC | Community-87 | 25 | Samp168 | R | 107 | Run2 | WC | Community_96 | 25 |

##### Step 4: Taxonomic classification

To perform taxonomic classification of the amplicon sequence variants (ASVs), feature classifiers were trained with the q2-feature-classifier provided by the QIIME2 using the instructions from the developers[6]. To this end, the SILVA database v132[8] was used as a reference database and the primers (Pro341F and Pro805R) were used to trim the reference database. After the taxonomic classification, samples amplified for 25 or 35 PCR cycles were merged using q2-feature-table merge[6].

##### Step 5: Removing contaminants and setting relative abundance thresholds based on positive and negative controls.

A two-fold approach was used to determine which ASVs should be removed based on negative (two sets) and positive controls (one set). For the first set of negative controls, 14 DNA extractions of the sheath buffer were sequenced to account for potential contamination during the DNA extraction. The second set of negative controls consisted of one extraction using the Chelex 100 solution as template for the PCR. All ASVs present in the sheath buffer DNA extracts or Chelex 100 solution were removed from the analysis. As a positive control, the ZymoBIOMICS™ Microbial Community Standard (Zymo Research Europe GmbH Freiburg, Germany) was sequenced in triplicate. Then, the ASVs from the eight different species were identified as present in this community using BLAST[9]. BLAST used as the q2-feature-classifier was unable to determine the phylogeny of all ASVs present in the ZymoBIOMICS™ Microbial Community Standard at the species level. The highest relative abundance of ASVs that did not belong to the ZymoBIOMICS™ Microbial Community Standard ASVs was 0.1%. Therefore, every ASV with a relative abundance of 0.1% or lower was removed from the 16S rRNA gene amplicon sequencing dataset.

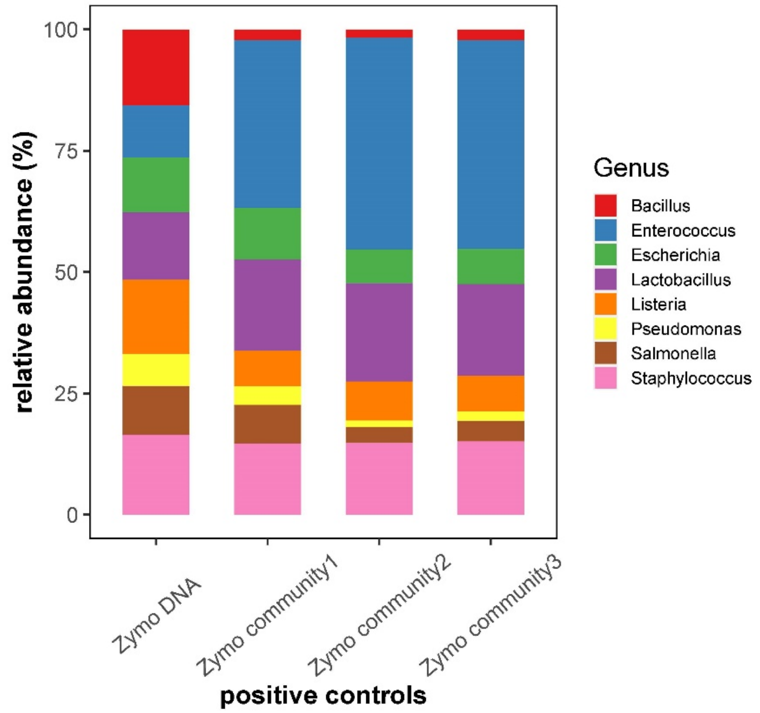


**Figure S6.1** Positive control for 16S rRNA gene amplicon sequencing. The ZymoBIOMICS™ Microbial Community Standard ASVs were used for this purpose (left column: data provided by the company) and three sets were run in parallel in accordance with the protocol described in this paper. The highest relative abundance of other strains not belonging to the standard was 0.1%. Therefore, every ASV with a relative abundance of 0.1% or lower was removed from the 16S rRNA gene amplicon sequencing dataset.

##### Step 6: Visualization of the amplicon sequencing data.

We used the phyloseq R package to visualise the taxonomic composition of the different samples used for 16S rRNA gene amplicon sequencing, in accordance with the instructions of the developers[10].

#### S7: Stability of communities under the influence of $RC_{10}$, $RC_{50}$ and $RC_{80}$

The stability properties of constancy, resistance RS and recovery RV were calculated for all SCs per community per phase and reactor in accordance with previous studies[1, 11]. Community stability was tested during Insular I phase, $RC_{10}$, $RC_{50}$, $RC_{80}$ and Insular II phase based on the Canberra distance measure.

#### Step 1: Constancy spaces for determining changes in microbial community composition caused by increasing $RC$

Constancy space reveals the fluctuation of community structure without the influence of external disturbance, thus indicating the inherent community variation. To investigate whether the shift in recycling rates $RC$ caused essential changes in microbial community compositions, in if so, to what extent, the deviations of the community structures of L1-L5 and of R from either the inoculum or start and end of the recycling phases were calculated using the Canberra distance (Fig. 2b; R-script v1.0, GitHub: https://github.com/fcentler/EcologicalStabilityPropertiesComputation). When the deviation of the community structure leaves the constancy space, the community is defined as unstable[1, 11]. The constancy space can be calculated in accordance with the procedure described by Liu et al.[11]. Briefly, the relative cell abundance data of all SCs of all samples from the balanced periods per phase and per reactor were included in the calculation of Canberra distances. This resulted, first, in an average sampling point, which described the mean of all samples. Second, Canberra distances from all samples to the average sampling point were calculated. Third, the constancy space was defined by determining the maximum Canberra distance per phase, which was possible due to the undisturbed balanced phases from which the samples were taken from. The mean value of the constancy space per phase was calculated for each of the five local communities L1-L5 (dashed lines in Fig. 2b).

A small value between 0 and 1 mirrors only small intrinsic fluctuations of a community and thus high constancy. Increased recycling rates resulted in a smaller constancy space of community structures of L1-L5, while no recycling allowed for higher fluctuations and a larger constancy space. The mean values of constancy spaces for the local communities L1-L5 were 0.44, 0.34, 0.26 and 0.27 for phases 1 to 4, respectively. In the Insular II phase, when recycling rates were stopped, the constancy values were similarly high as in Insular I phase (0.43, Table S7.1). The phase-to-phase variation was confirmed by Wilcoxon test ($p = 0.008$, 0.008, 0.841 and 0.008 respectively), with the exception of that between $RC_{50}$ and $RC_{80}$.

**Table S7.1** Constancy space calculated based on Canberra distance.

| reactor | Insular I | $RC_{10}$ | $RC_{50}$ | $RC_{80}$ | Insular II |
|---|---|---|---|---|---|
| L1 | 0.44 | 0.33 | 0.26 | 0.27 | 0.50 |
| L2 | 0.46 | 0.34 | 0.25 | 0.32 | 0.48 |
| L3 | 0.41 | 0.33 | 0.28 | 0.27 | 0.37 |
| L4 | 0.46 | 0.34 | 0.28 | 0.24 | 0.39 |
| L5 | 0.43 | 0.37 | 0.22 | 0.25 | 0.41 |
| mean ± sd | 0.44 ± 0.02 | 0.34 ± 0.02 | 0.26 ± 0.02 | 0.27 ± 0.03 | 0.43 ± 0.06 |
| R | 0.28 | 0.25 | 0.22 | 0.26 | 0.39 |

##### Step 2: Resistance RS and recovery RV for determining changes in microbial community composition caused by increasing $RC$

The values for resistance RS and recovery RV were used to determine whether microbiomes change in composition between the phases, that is, Insular I phase to $RC_{10}$, $RC_{10}$ to $RC_{50}$, $RC_{50}$ to $RC_{80}$ and $RC_{80}$ to Insular II phase. The values were calculated by the degree of deviation from the starting point (resistance RS) and the ability of the microbiome to return to the starting point (recovery RV). The deviation from the inoculum (day 0) or from the respective endpoints of previous balanced phases (days 26, 47, 64, and 89, respectively; Fig. 2b) were used to calculate the deviations that occurred. These phase-to-phase variations were confirmed by significance testing (Wilcoxon test, $p$ = 0.0079, 0.1508 and 0.008 respectively; Fig. S7.1). The RS value was lowest during the shift from Insular I phase to $RC_{10}$ (mean 0.46 ± 0.05 for L1-L5) suggesting the greatest changes in community composition under these conditions. The shifts $RC_{10}$ to $RC_{50}$ and $RC_{50}$ to $RC_{80}$ showed the highest RS (mean 0.57 ± 0.03 and 0.60 ± 0.03, respectively, for L1-L5) and therefore indicated small changes of microbiome structures. We did not observe much changes in RV in all phases and for all communities (Table S7.1).

**Table. S7.2** Resistance (RS) and recovery (RV) for the shifts between phases in L1-L5 and R based on Canberra distance.

| reactor | Insular I to $RC_{10}$ | | $RC_{10}$ to $RC_{50}$ | | $RC_{50}$ to $RC_{80}$ | | $RC_{80}$ to Insular II | |
|---|---|---|---|---|---|---|---|---|
| | RS | RV | RS | RV | RS | RV | RS | RV |
| L1 | 0.52 | 0.06 | 0.57 | 0.23 | 0.61 | 0.03 | 0.44 | 0.19 |
| L2 | 0.41 | 0.02 | 0.53 | 0 | 0.61 | 0.01 | 0.52 | 0.11 |
| L3 | 0.47 | 0.02 | 0.6 | 0 | 0.6 | 0.14 | 0.52 | 0.04 |
| L4 | 0.39 | 0.07 | 0.59 | 0.14 | 0.55 | 0.12 | 0.5 | 0.12 |
| L5 | 0.49 | 0.01 | 0.58 | 0.01 | 0.62 | 0.02 | 0.5 | 0.09 |
| mean ± sd | 0.46 ± 0.05 | 0.04 ±0.03 | 0.57 ± 0.03 | 0.08 ± 0.10 | 0.60 ± 0.03 | 0.06 ± 0.06 | 0.50 ± 0.03 | 0.11 ± 0.05 |
| R | 0.57 | 0.18 | 0.67 | 0.08 | 0.6 | 0.01 | 0.53 | 0.15 |

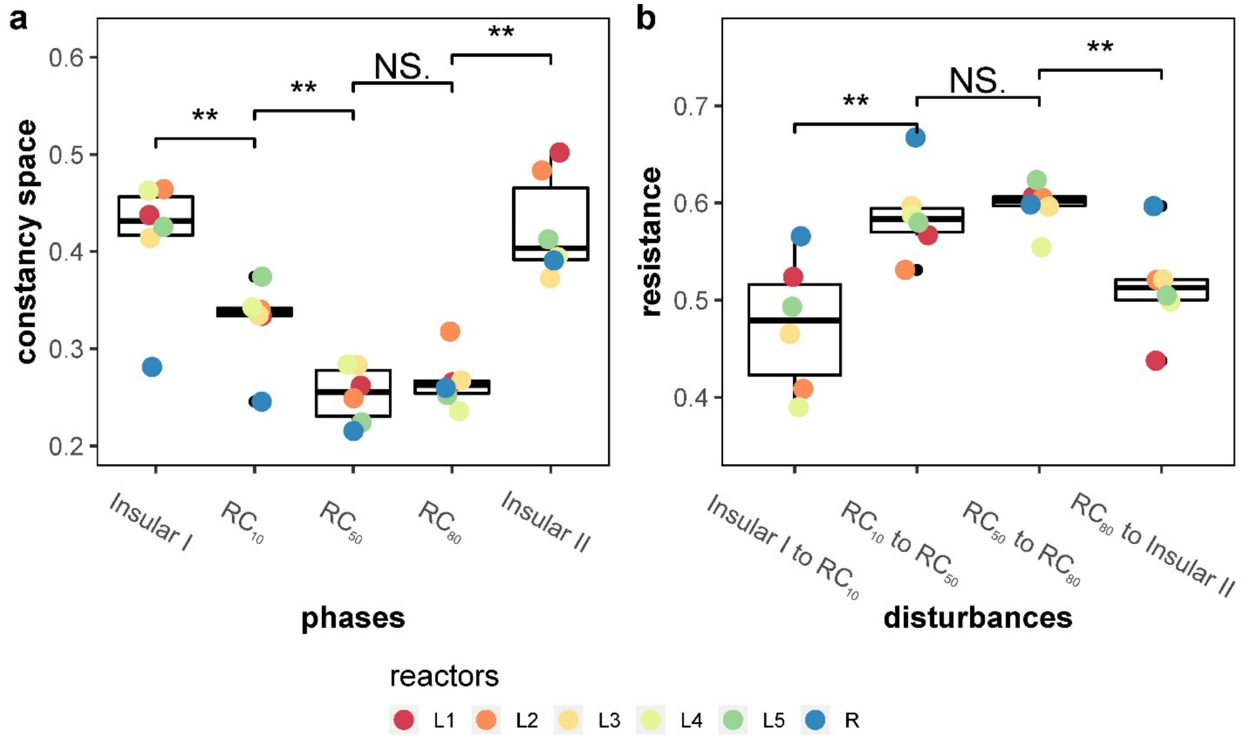


**Figure S7.1** The stability properties of **a)** constancy and **b)** resistance RS were calculated for the local microbiomes L1-L5 and the regional pool R. A small constancy space marks a high constancy and high resistance values mark a high resistance behaviour. The stars indicate a significant difference in stability properties of local communities L1-L5 between two successive phases (Wilcoxon; $*p \leqslant 0.05$ and $**p \leqslant 0.01$).

#### S8: Analysis of microbial community composition

##### Step 1: Visualisation of microbial community composition

The microbial community composition (i.e. fingerprint) was determined by flow cytometry. The variations in the composition of the six communities were shown by the relative cell abundance and absolute cell abundance variations for the dominant SCs (Figs. S8.1, S8.2; dominant SC: relative cell abundance >1/number of gates = 1.25% or absolute cell abundance > cell number of total community/number of gates).

The fate of all dominant SCs through time (Fig. S8.3) was shown based on relative cell abundance. Using the R package ‘vegan’[12], the dissimilarity of the samples was shown as NMDS plots based on the Bray-Curtis distance measure calculated based on relative cell abundance in Fig. 2a.

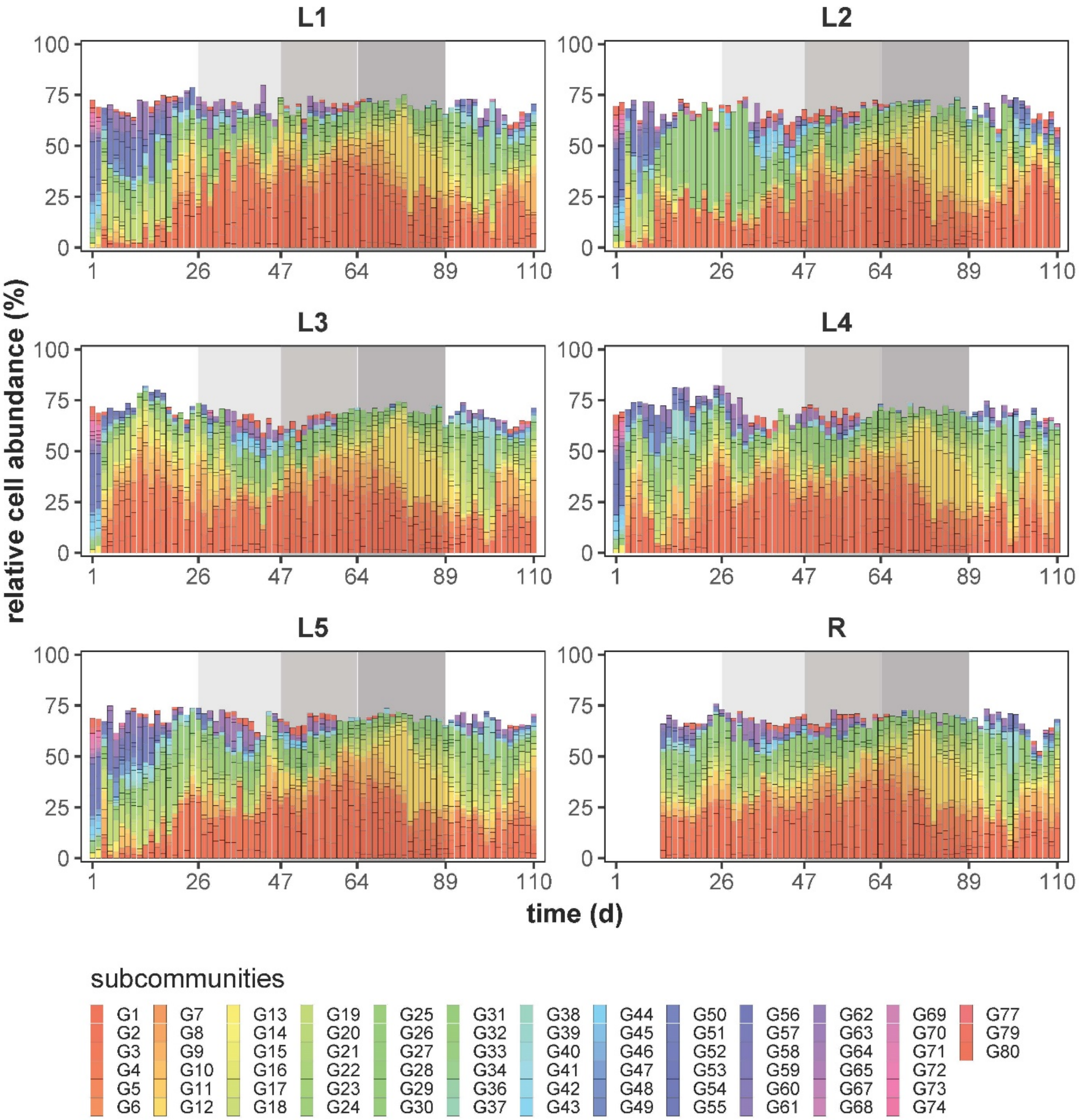


**Figure S8.1** The relative cell abundance distribution per dominant SC and reactors (relative abundance > 1/number of gates = 1.25% in the corresponding community sample) over time. Each SC is represented by a different colour. The relative abundance of each SC is given in %. The shaded areas represent different phases with changed $RC$.

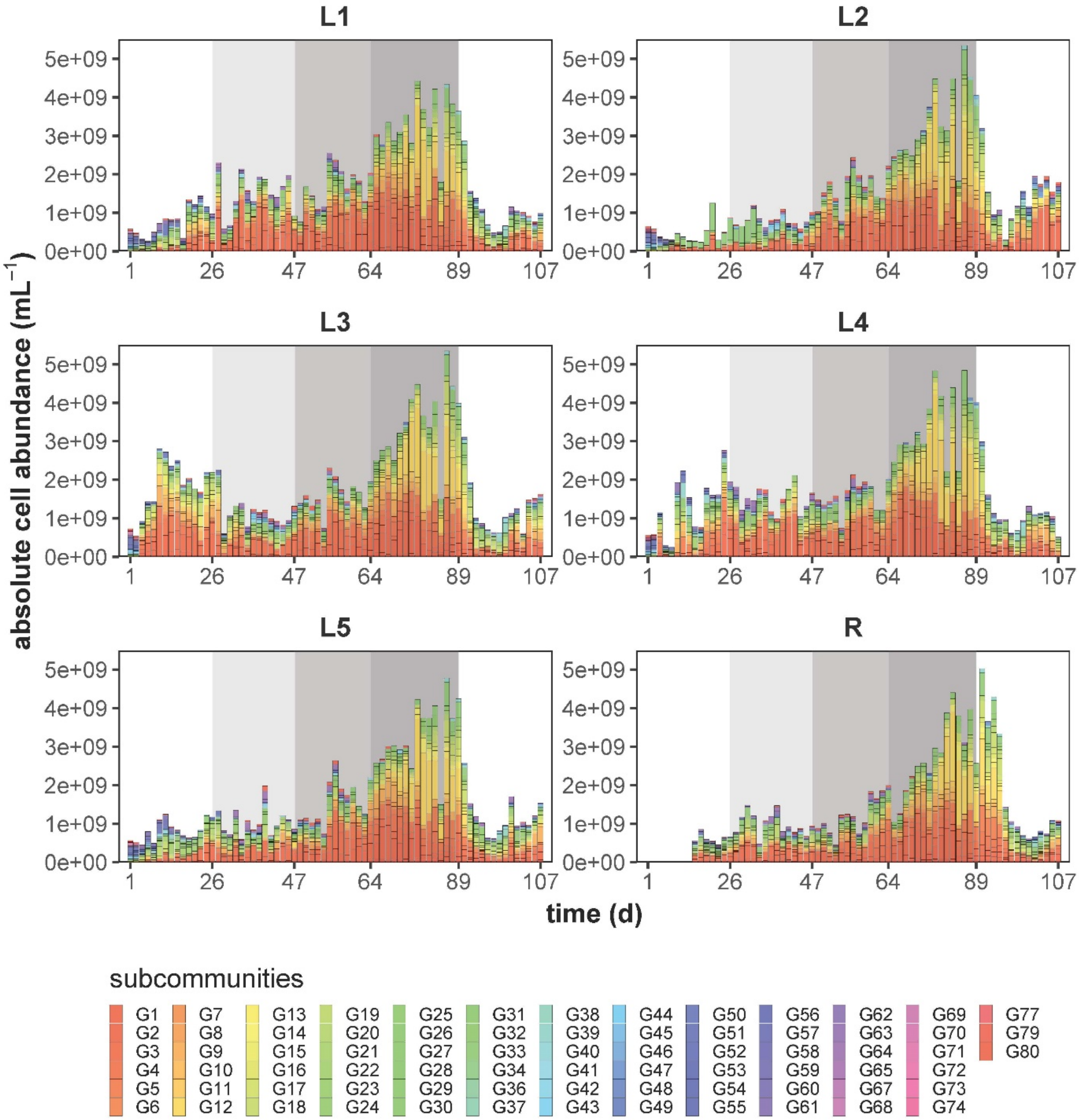


**Figure S8.2** The absolute cell abundance distribution per dominant SCs and reactors (absolute cell abundance > cell number of total community/number of gates) over time. Each SC is represented by a different colour. The absolute cell abundance of each SC is given in cells mL$^{-1}$. The shaded areas represent different phases with changed $RC$.

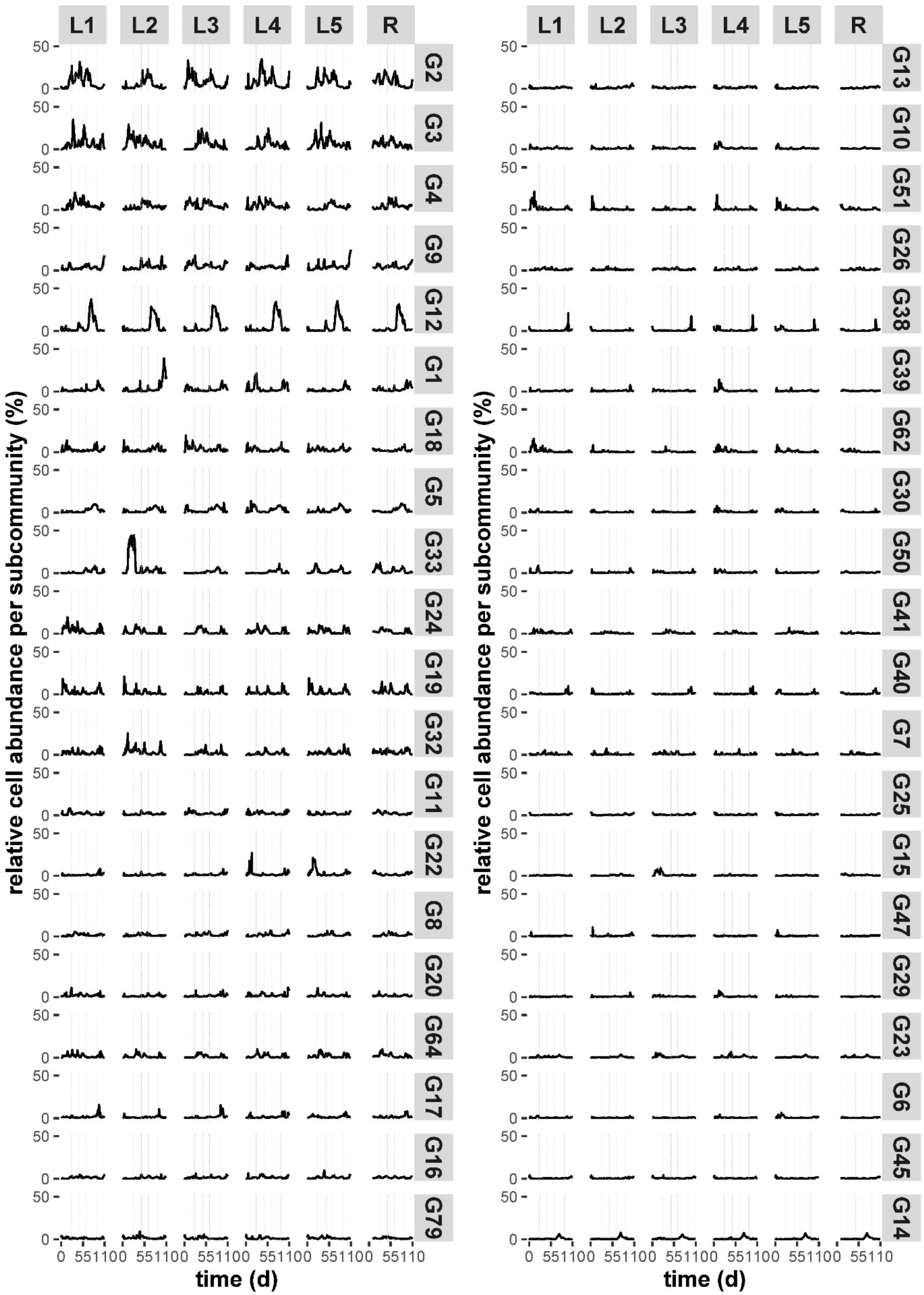
L1
L2
L3
L4
L5
R
G2
G3
G4
G9
G12
G1
G18
G5
G33
G24
G19
G32
G11
G22
G8
G20
G64
G17
G16
G79
G13
G10
G51
G26
G38
G39
G62
G30
G50
G41
G40
G7
G25
G15
G47
G29
G23
G6
G45
G14
relative cell abundance per subcommunity (%)
time (d)

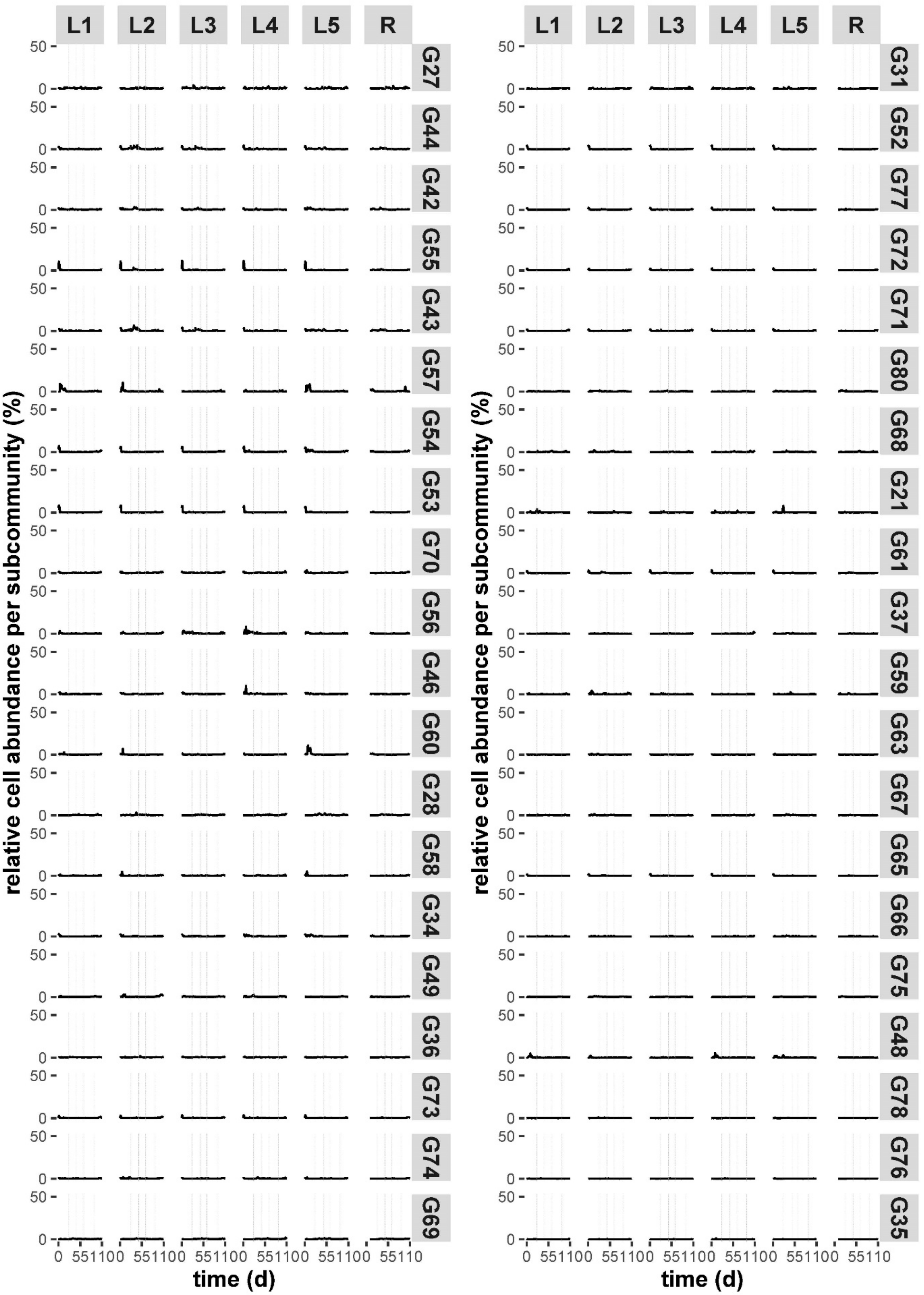
L1
L2
L3
L4
L5
R
G27
G44
G42
G55
G43
G57
G54
G53
G70
G56
G46
G60
G28
G58
G34
G49
G36
G73
G74
G69
G31
G52
G77
G72
G71
G80
G68
G21
G61
G37
G59
G63
G67
G65
G66
G75
G48
G78
G76
G35
50
0
0 551100 551100 551100 551100 551100 55110
relative cell abundance per subcommunity (%)
time (d)

**Figure S8.3** The fate of SCs according to relative cell abundance. SCs are ranked in descending order of mean relative cell abundance over the whole experiment. The dashed lines indicate the different phases with changed $RC$.

##### Step 2: Comparison of effluents between local communities L1-L5 and the regional pool R

A PERMANOVA (permutational multivariate analysis of variance) was applied using the R package 'vegan'[11] to determine the difference between the effluents from the local communities L1-L5 and the regional pool R. We wanted to know whether the microbial composition of R was similar to the sum of effluents from L1-L5. Briefly, samples from day 12 to day 23 (Insular I phase) were used. For each day, the values of absolute cell abundance per SC from L1-L5 were summed up and then normalised to relative cell abundance to simulate the sum of L1-L5 effluents. The sum of L1-L5 effluents was compared with the R community of the same days (permutations = 9999, method = 'bray'). The results showed that the composition of the R community was divergent from that of L1-L5 (PERMANOVA, F = 8.452, $p$ = 0.0002, $R^2$ = 0.346). Thus, the composition of the R community was not only determined by the immigration from L1-L5, but also by its unique operational parameters (e.g., dilution time, no nutrient feeding).

##### Step 3: α-, γ- and β-diversity values of microbial communities L1-L5 and R

The α-diversity was determined by counting the numbers of dominant SCs (relative cell abundance > 1/number of gates = 1.25%) per local community L1-L5 and the regional pool R.

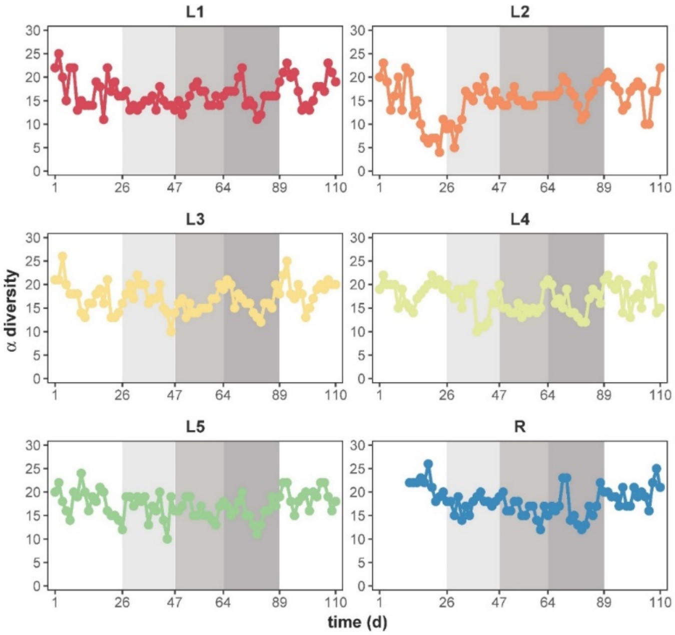


**Figure S8.4** α-diversity of the composition of the five local communities L1-L5 and the regional pool R. The α-diversity was determined by counting the numbers of dominant SCs (relative cell abundance > 1/number of gates = 1.25%) in each community. The colour indicates the respective community as specified in Fig. 1. The shaded areas represent different phases with changed $RC$.

The γ-diversity was determined by counting the numbers of SCs among a total of 80 SCs that were dominant (relative cell abundance > 1/number of gates = 1.25%) in at least one local community within the whole metacommunity, which was expressed as the richness of SCs in the metacommunity. The γ-diversity showed remarkable phase-to-phase variation (Wilcoxon test, $p = 3.416\times10^{-4}$, $6.258\times10^{-4}$, 0.045, $3.605\times10^{-4}$, respectively; Fig. S8.5). The γ-diversity declined from 39.73 ± 3.17 SCs in Insular I phase to 18.8 ± 3.85 SCs in $RC_{80}$ and increased again to 33.11 ± 3.62 SCs in Insular II phase (Table 1). Thus, the γ-diversity decreased with increasing $RC$ values, but recovered to a level when the recycling flow was stopped.

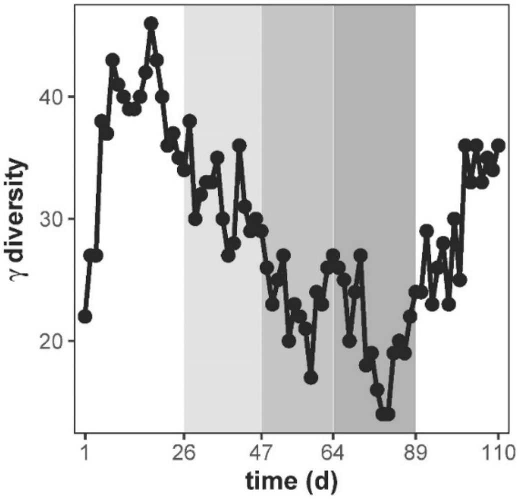


**Figure S8.5** γ-diversity of the metacommunity composed of five local communities (L1-L5). The γ-diversity was determined by counting the numbers of SCs among a total of 80 SCs that were dominant (relative cell abundance > 1/number of gates = 1.25%) in at least one local community. The shaded areas represent different phases with changed $RC$.

The β-diversity was computed as the number of unique dominant SCs, which were not shared by pairwise samples. Intra-community β-diversity compared samples from successive time points per reactor (Fig. 2a), while inter-community β-diversity compared samples from the same time point but from different reactors (Fig. 2b). Significant differences of intra-community β-diversity between $RC_{50}$ and zero or low recycling phases were found ($p = 5.505\times10^{-5}$ for Insular I, $8.016\times10^{-4}$ for $RC_{10}$, $1.625\times10^{-7}$ for Insular II), as well as between $RC_{80}$ and these phases respectively ($p = 3.737×10^{-8}$ for Insular I, $1.753×10^{-6}$ for $RC_{10}$, $1.983×10^{-11}$ for Insular II). Wilcoxon test also showed significant differences of inter-community β-diversity between successive phases for L vs. L ($p = 3.848×10^{-14}$, $2.2×10^{-16}$, $1.033×10^{-8}$, $2.2×10^{-16}$) and L vs. R ($p = 3.203×10^{-8}$, $5.086×10^{-10}$, $4.838×10^{-10}$, $3.203×10^{-8}$, $2.2×10^{-16}$). A summary of α-, γ- and β-diversity values is shown in Table 1.

Drift events were calculated from balanced periods when the temporal community composition variation (i.e. intra-community β-diversity) was higher than a defined threshold (Fig. 2a and Table S8.1). The threshold was set to 8.76 and was calculated as the mean value for intra-community β-diversity of L1-L5 of the Insular I phase during the balanced periods. To investigate which microorganisms were involved in the drift events, we sorted the relevant SCs that massively changed their relative cell abundance. After cell sorting, 16S rRNA sequencing analysis was performed (Supplementary Information S9). We studied three drift events: drift1 ($RC_{10}$) in L2 and L5 at day 47, drift2 ($RC_{50}$) in L3 and L5 at day 61 and drift3 (Insular II) in L1 and L3 at day 100. For comparison of community states between before and after the drifts, day 44, day 58 and day 99 were taken as before-drift community states, respectively, for drift1, 2 and 3 (Table S8.2).

**Table S8.1** Drifts occurring on certain days in local communities L1-L5. The values showed the intra-community β-diversity between two successive time points. The drifts were identified by intra-community β-diversity values being above a threshold of $> 8.76$ (i.e. mean intra-community β-diversity value during the balanced periods of Insular I phase of all five local communities). A dash (-) means that no drift occurred in the respective communities at the time. The drifts in bold were chosen for cell sorting (Supplementary Information S9).

| time (d) | Insular I | | | | | | | | | | | | $RC_{10}$ | | | | | | | |
|---|---|---|---|---|---|---|---|---|---|---|---|---|---|---|---|---|---|---|---|---|
| | *8* | *9* | *12* | *13* | *14* | *15* | *16* | *19* | *20* | *21* | *23* | *26* | *34* | *35* | *36* | *40* | *41* | *43* | *44* | ***47*** |
| L1 | - | - | 10 | 10 | 13 | - | - | 19 | 15 | - | 9 | 20 | - | - | - | - | 9 | 9 | - | - |
| L2 | 19 | 15 | 11 | 9 | 11 | - | - | - | - | - | - | - | 13 | 11 | - | - | 11 | - | 15 | **24** |
| L3 | - | - | - | - | - | - | - | - | 12 | 12 | 11 | - | - | 16 | - | 9 | 9 | - | - | - |
| L4 | 13 | 15 | 13 | 13 | - | 11 | 15 | 12 | - | 9 | - | 9 | 9 | 10 | 12 | - | - | 14 | 14 | 12 |
| L5 | 9 | 10 | 10 | 9 | 13 | 11 | 11 | 10 | - | - | - | - | - | 10 | - | 12 | 10 | 10 | 13 | **15** |

| time (d) | $RC_{50}$ | | | | $RC_{80}$ | | Insular II | | | | | | | | | |
|---|---|---|---|---|---|---|---|---|---|---|---|---|---|---|---|---|
| | *58* | ***61*** | *62* | *63* | *76* | *89* | *97* | *98* | *99* | ***100*** | *103* | *104* | *105* | *106* | *107* | *110* |
| L1 | 9 | - | - | - | - | 9 | - | - | 15 | **20** | 9 | - | - | 10 | 10 | 12 |
| L2 | - | 10 | 10 | 10 | 11 | - | 11 | - | 19 | 9 | 19 | 10 | - | 9 | - | 9 |
| L3 | - | **18** | - | - | - | - | - | 15 | - | **20** | 20 | 15 | - | - | - | 14 |
| L4 | - | 11 | - | - | - | - | - | 15 | - | 17 | 13 | 12 | - | 20 | 16 | 9 |
| L5 | - | **13** | - | - | - | - | - | 9 | - | 16 | 9 | 9 | 10 | 11 | - | 10 |

**Table S8.2** Relative cell abundance of SCs (%) before and after drift1 (day 47, $RC_{10}$) in L2 and L5, drift2 (day 61, $RC_{50}$) in L3 and L5, and drift3 (day 100, Insular II) in L1 and L3. The pink background of squares marks the dominance of the SCs (relative cell abundance > 1/number of gates = 1.25%). A dash (-) means that the SC was not involved in the drift in the corresponding reactor. The SCs in bold were chosen for cell sorting (Supplementary Information S9).

| | reactor | state | **G1** | **G3** | G4 | **G7** | G8 | G10 | G11 | G13 | **G16** | G18 | G19 | G20 | **G22** | G23 | **G24** | **G26** | G27 | G32 | **G33** | G41 | G42 | G43 | G47 | G50 | G59 | G62 | G64 | **G79** | |
|---|---|---|---|---|---|---|---|---|---|---|---|---|---|---|---|---|---|---|---|---|---|---|---|---|---|---|---|---|---|---|---|
| drift 1 (day 47) | L2 | before | 12.6 | - | 1.0 | 2.8 | 1.2 | 1.6 | 0.6 | 2.2 | 0.5 | 1.4 | - | 0.5 | 0.8 | - | 4.0 | 4.8 | - | 1.1 | 0.3 | 2.1 | 1.7 | - | 3.3 | 0.2 | 1.5 | 1.8 | 2.3 | 0.9 | |
| | | after | 0.3 | - | 2.5 | 0.2 | 1.8 | 0.6 | 6.0 | 0.4 | 5.0 | 0.6 | - | 2.2 | 5.1 | - | 0.4 | 0.9 | - | 1.7 | 8.0 | 0.9 | 0.7 | - | 0.5 | 2.7 | 0.0 | 0.0 | 0.1 | 5.1 | |
| | L5 | before | - | 1.2 | - | 6.3 | - | - | 2.8 | 2.5 | - | - | 0.6 | 2.9 | 1.3 | 1.3 | - | 2.7 | 2.3 | 0.6 | - | 0.5 | - | 0.1 | - | - | 2.5 | - | - | 0.9 | |
| | | after | - | 9.1 | - | 1.1 | - | - | 1.1 | 0.6 | - | - | 1.5 | 1.0 | 0.4 | 1.1 | - | 1.0 | 0.4 | 2.7 | - | 1.6 | - | 1.4 | - | - | 0.3 | - | - | 5.1 | |
| drift 2 (day 61) | reactor | time (d) | **G5** | **G7** | **G12** | G13 | G16 | G17 | **G18** | **G19** | G20 | G22 | **G24** | G26 | G27 | G32 | G41 | G51 | G64 | G79 | | | | | | | | | | | |
| | L3 | before | 0.9 | 0.5 | 0.6 | 0.4 | 1.2 | 1.5 | 0.8 | 5.2 | 1.0 | 1.2 | 2.5 | 1.2 | 0.50 | 3.3 | 1.9 | 1.3 | 2.7 | 2.0 | | | | | | | | | | | |
| | | after | 2.2 | 4.1 | 2.2 | 1.5 | 1.9 | 0.9 | 2.3 | 0.6 | 1.9 | 1.3 | 0.4 | 3.7 | 1.6 | 0.2 | 0.8 | 0.3 | 0.4 | 0.6 | | | | | | | | | | | |
| | L5 | before | 0.6 | 0.5 | - | - | 0.8 | - | 0.6 | 5.5 | 0.7 | - | 3.3 | - | 0.6 | 5.2 | 2.0 | 1.3 | 3.7 | 2.7 | | | | | | | | | | | |
| | | after | 2.4 | 2.6 | - | - | 1.8 | - | 1.4 | 0.4 | 1.5 | - | 0.3 | - | 2.2 | 1.1 | 1.0 | 0.2 | 0.4 | 0.8 | | | | | | | | | | | |
| drift 3 (day 100) | reactor | time (d) | G2 | **G3** | **G4** | **G5** | **G7** | G8 | G11 | **G12** | G13 | G15 | G17 | **G18** | **G20** | G22 | **G24** | G26 | G28 | G30 | G31 | G32 | **G38** | G40 | G41 | G51 | G56 | G57 | G62 | G64 | G68 |
| | L1 | before | - | 0.0 | 0.3 | 0.6 | 0.2 | - | 4.3 | - | 1.0 | - | 6.8 | - | 7.0 | 8.4 | 0.9 | 0.6 | 1.6 | 1.6 | - | - | 21.0 | 9.6 | 0.0 | 1.1 | - | - | 0.0 | 0.1 | 1.3 |
| | | after | - | 11.4 | 3.6 | 3.2 | 3.3 | - | 0.5 | - | 1.8 | - | 1.2 | - | 0.5 | 1.1 | 10.9 | 1.6 | 0.8 | 0.2 | - | - | 0.4 | 0.3 | 1.9 | 2.9 | - | - | 2.3 | 7.4 | 0.3 |
| | L3 | before | 0.6 | 0.0 | 0.3 | 1.2 | - | 1.9 | - | 2.2 | - | 0.2 | 4.0 | 0.6 | 3.5 | - | 0.7 | - | - | 2.6 | 0.2 | 2.1 | 17.5 | 7.3 | 0.0 | - | 0.4 | 1.7 | - | 0.0 | - |
| | | after | 2.8 | 11.2 | 2.4 | 11.5 | - | 1.2 | - | 0.7 | - | 1.5 | 0.9 | 4.1 | 0.6 | - | 5.2 | - | - | 0.5 | 2.2 | 0.9 | 0.8 | 0.5 | 2.1 | - | 1.8 | 0.1 | - | 3.3 | - |

#### Step 4: Partitioning of β-diversity to reveal turnover and nestedness of SCs

The community composition variation was partitioned into species replacement and species loss to determine turnover and nestedness values using a method developed for microbial community flow cytometric data[13]. The method was used to study the temporal pattern of intra- and inter-community β-diversity loss when the recycling rate $RC$ was increased. Using multiple-site measures in the R package 'betapart', the turnover $\beta_{SIM}$ and nestedness $\beta_{NES}$ components of Sørensen dissimilarity were calculated[14, 15] (Fig. S8.6 and Table S8.3). Multisite intra-community β-diversity was calculated over all samples from the balanced period per phase per community. Multisite inter-community β-diversity was calculated over five local communities (L1-L5) per time point. The lowest intra-community β-diversity values for turnover $\beta_{SIM}$ were always at $RC_{80}$ with a mean value of 0.34 ± 0.02 and the highest values for nestedness $\beta_{NES}$ were also always at $RC_{80}$ with a mean value of 0.14 ± 0.02. The inter-community β-diversity values for turnover $\beta_{SIM}$ were also lowest at $RC_{80}$ (0.11 ± 0.04) during the balanced periods, while the values for nestedness $\beta_{NES}$ remained roughly the same (Table S8.3).

We also investigated which SCs were nested. Nested SCs were determined if they remained dominant at all time points over the balanced period in the respective reactor and phase (relative cell abundance > 1.25, Table S8.4). Generally, fewer nested SCs were observed in insular phases than in RC phases. In insular phases, nested but different SCs were found in the various local communities, while in RC phases, some of the SCs were nested in all reactors L1-L5 per phase or over all phases, that is, G2 in $RC_{10}$; G2, G9, G3, G4 and G11 in $RC_{50}$; and G2, G9, G4, G5, G12 and G14 in $RC_{80}$.

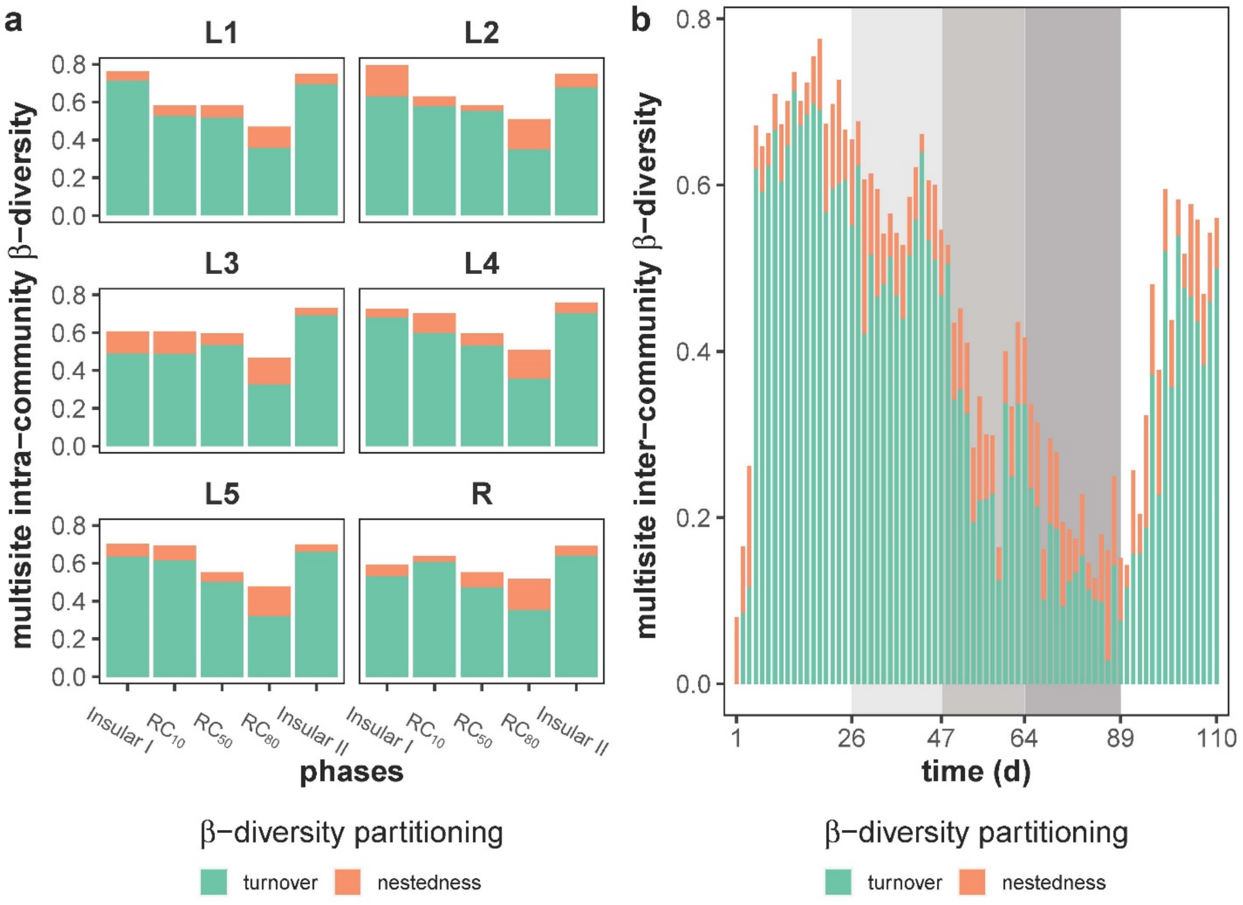


**Figure S8.6** Partitioning of β-diversity into turnover (green) and nestedness (orange) components. a) Multisite intra-community β-diversity was calculated over all samples from the balanced period per phase per community and b) multisite inter-community β-diversity was calculated over five local communities (L1-L5) per time point. The shaded areas represent different phases with changed $RC$.

**Table S8.3** Summary of turnover and nestedness components of multisite intra- and inter-community comparisons (Liu & Müller[13]) as mean ± standard deviation (sd) per phase. The mean ± sd values were all calculated among local communities L1-L5 during balanced periods if not clarified otherwise. Asterisks indicate that the diversity values in these phases were significantly different from those in any other phases, at *$p \leqslant 0.05$, **$p \leqslant 0.01$ or ***$p \leqslant 0.001$.

| comparison | components | Insular I | RC$_{10}$ | RC$_{50}$ | RC$_{80}$ | Insular II |
|---|---|---|---|---|---|---|
| multisite intra-community | turnover | 0.63 ± 0.09 | 0.56 ± 0.05 | 0.53 ± 0.02 | 0.34 ± 0.02** | 0.68 ± 0.02 |
| | nestedness | 0.09 ± 0.05 | 0.08 ± 0.03 | 0.05 ± 0.01 | 0.14 ± 0.02 | 0.05 ± 0.01 |
| multisite inter-community | turnover | 0.64 ± 0.05*** | 0.52 ± 0.06* | 0.24 ± 0.07** | 0.11 ± 0.04** | 0.43 ± 0.09* |
| | nestedness | 0.07 ± 0.03 | 0.07 ± 0.02 | 0.08 ± 0.03 | 0.07 ± 0.04 | 0.09 ± 0.03 |

**Table S8.4** List of nested subcommunities (SCs) in respective reactors and phases. Nested SCs were determined if they remained dominant (relative cell abundance > 1.25) at all time points over the balanced period in the respective reactor and phase. Nested SCs are marked with the number '1' and a pink background. The SCs were ranked from left to right according to the descending order of frequencies of being nested (numbers of pink squares).

| reactor | phase | G2 | G9 | G3 | G4 | G11 | G8 | G18 | G22 | G5 | G12 | G14 | G24 | G1 | G33 | G64 | G15 | G32 | G13 | G17 | G19 | G20 | G26 | G38 | G43 |
|---|---|---|---|---|---|---|---|---|---|---|---|---|---|---|---|---|---|---|---|---|---|---|---|---|---|
| L1 | Insular I | 0 | 0 | 0 | 0 | 0 | 0 | 0 | 0 | 0 | 0 | 0 | 0 | 0 | 0 | 0 | 0 | 0 | 0 | 0 | 0 | 0 | 0 | 0 | 0 |
| L2 | Insular I | 0 | 0 | 1 | 0 | 0 | 0 | 0 | 0 | 0 | 0 | 0 | 0 | 0 | 0 | 0 | 0 | 1 | 0 | 0 | 0 | 0 | 0 | 0 | 0 |
| L3 | Insular I | 1 | 1 | 0 | 1 | 1 | 0 | 1 | 0 | 0 | 0 | 0 | 0 | 0 | 0 | 0 | 1 | 0 | 0 | 0 | 0 | 0 | 0 | 0 | 0 |
| L4 | Insular I | 0 | 0 | 0 | 0 | 0 | 0 | 0 | 1 | 0 | 0 | 0 | 0 | 0 | 0 | 0 | 0 | 0 | 0 | 0 | 0 | 0 | 0 | 0 | 0 |
| L5 | Insular I | 0 | 0 | 0 | 0 | 0 | 0 | 0 | 1 | 0 | 0 | 0 | 0 | 0 | 0 | 0 | 0 | 0 | 0 | 1 | 0 | 0 | 0 | 1 | 0 |
| R | Insular I | 1 | 1 | 0 | 0 | 1 | 0 | 1 | 1 | 0 | 0 | 0 | 1 | 0 | 0 | 0 | 1 | 0 | 0 | 0 | 0 | 0 | 0 | 0 | 0 |
| L1 | RC$_{10}$ | 1 | 0 | 1 | 1 | 0 | 1 | 0 | 0 | 0 | 0 | 0 | 1 | 0 | 0 | 0 | 0 | 0 | 0 | 0 | 0 | 0 | 0 | 0 | 0 |
| L2 | RC$_{10}$ | 1 | 0 | 1 | 0 | 0 | 0 | 0 | 0 | 0 | 0 | 0 | 1 | 0 | 0 | 1 | 0 | 0 | 0 | 0 | 0 | 0 | 0 | 0 | 1 |
| L3 | RC$_{10}$ | 1 | 1 | 1 | 0 | 0 | 0 | 1 | 0 | 0 | 0 | 0 | 1 | 0 | 0 | 1 | 0 | 0 | 0 | 0 | 0 | 0 | 0 | 0 | 0 |
| L4 | RC$_{10}$ | 1 | 0 | 0 | 0 | 1 | 0 | 0 | 0 | 0 | 0 | 0 | 0 | 0 | 0 | 0 | 0 | 0 | 0 | 0 | 0 | 0 | 0 | 0 | 0 |
| L5 | RC$_{10}$ | 1 | 0 | 0 | 0 | 0 | 0 | 0 | 0 | 0 | 0 | 0 | 0 | 0 | 0 | 0 | 0 | 0 | 0 | 0 | 0 | 1 | 0 | 0 | 0 |
| R | RC$_{10}$ | 1 | 0 | 1 | 0 | 0 | 0 | 0 | 0 | 0 | 0 | 0 | 1 | 0 | 0 | 1 | 0 | 0 | 0 | 0 | 0 | 0 | 0 | 0 | 0 |
| L1 | RC$_{50}$ | 1 | 1 | 1 | 1 | 1 | 1 | 0 | 0 | 0 | 0 | 0 | 0 | 0 | 0 | 0 | 0 | 0 | 0 | 0 | 0 | 0 | 0 | 0 | 0 |
| L2 | RC$_{50}$ | 1 | 1 | 1 | 1 | 1 | 0 | 0 | 0 | 0 | 0 | 0 | 0 | 0 | 0 | 0 | 0 | 0 | 0 | 0 | 0 | 0 | 0 | 0 | 0 |
| L3 | RC$_{50}$ | 1 | 1 | 1 | 1 | 1 | 1 | 0 | 0 | 0 | 0 | 0 | 0 | 0 | 0 | 0 | 0 | 0 | 0 | 0 | 0 | 0 | 0 | 0 | 0 |
| L4 | RC$_{50}$ | 1 | 1 | 1 | 1 | 1 | 1 | 0 | 0 | 0 | 0 | 0 | 0 | 0 | 0 | 0 | 0 | 0 | 0 | 0 | 0 | 0 | 0 | 0 | 0 |
| L5 | RC$_{50}$ | 1 | 1 | 1 | 1 | 1 | 1 | 0 | 0 | 0 | 0 | 0 | 0 | 0 | 0 | 0 | 0 | 0 | 0 | 0 | 0 | 0 | 1 | 0 | 0 |
| R | RC$_{50}$ | 1 | 1 | 1 | 1 | 1 | 1 | 0 | 0 | 0 | 0 | 0 | 0 | 0 | 0 | 0 | 0 | 1 | 0 | 0 | 1 | 0 | 0 | 0 | 0 |
| L1 | RC$_{80}$ | 1 | 1 | 1 | 1 | 0 | 0 | 1 | 0 | 1 | 1 | 1 | 0 | 0 | 0 | 0 | 0 | 0 | 0 | 0 | 0 | 0 | 0 | 0 | 0 |
| L2 | RC$_{80}$ | 1 | 1 | 0 | 1 | 0 | 0 | 0 | 0 | 1 | 1 | 1 | 0 | 0 | 0 | 0 | 0 | 0 | 0 | 0 | 0 | 0 | 0 | 0 | 0 |
| L3 | RC$_{80}$ | 1 | 1 | 1 | 1 | 0 | 0 | 1 | 0 | 1 | 1 | 1 | 0 | 0 | 1 | 0 | 0 | 0 | 0 | 0 | 0 | 0 | 0 | 0 | 0 |
| L4 | RC$_{80}$ | 1 | 1 | 0 | 1 | 0 | 0 | 0 | 0 | 1 | 1 | 1 | 0 | 0 | 1 | 0 | 0 | 0 | 0 | 0 | 0 | 0 | 0 | 0 | 0 |
| L5 | RC$_{80}$ | 1 | 1 | 1 | 1 | 0 | 0 | 1 | 0 | 1 | 1 | 1 | 0 | 0 | 1 | 0 | 0 | 0 | 0 | 0 | 0 | 0 | 0 | 0 | 0 |
| R | RC$_{80}$ | 1 | 1 | 1 | 1 | 0 | 0 | 1 | 0 | 1 | 1 | 1 | 0 | 0 | 1 | 0 | 0 | 0 | 0 | 0 | 0 | 0 | 0 | 0 | 0 |
| L1 | Insular II | 0 | 0 | 0 | 0 | 0 | 0 | 0 | 0 | 0 | 0 | 0 | 0 | 1 | 0 | 0 | 0 | 0 | 0 | 0 | 0 | 0 | 0 | 0 | 0 |
| L2 | Insular II | 0 | 1 | 0 | 0 | 0 | 0 | 0 | 0 | 0 | 0 | 0 | 0 | 1 | 0 | 0 | 0 | 0 | 1 | 0 | 0 | 0 | 0 | 0 | 0 |
| L3 | Insular II | 0 | 1 | 0 | 0 | 0 | 0 | 0 | 1 | 0 | 0 | 0 | 0 | 0 | 0 | 0 | 0 | 0 | 0 | 0 | 0 | 0 | 0 | 0 | 0 |
| L4 | Insular II | 0 | 1 | 0 | 0 | 0 | 0 | 0 | 1 | 0 | 0 | 0 | 0 | 0 | 0 | 0 | 0 | 0 | 0 | 0 | 0 | 0 | 0 | 0 | 0 |
| L5 | Insular II | 0 | 1 | 0 | 0 | 0 | 1 | 0 | 1 | 0 | 0 | 0 | 0 | 1 | 0 | 0 | 0 | 0 | 0 | 0 | 0 | 0 | 0 | 0 | 0 |
| R | Insular II | 0 | 1 | 0 | 0 | 0 | 1 | 0 | 1 | 0 | 0 | 0 | 0 | 1 | 0 | 0 | 0 | 0 | 0 | 0 | 0 | 0 | 0 | 0 | 0 |

#### S9: Taxonomic composition of whole communities and sorted SCs

##### Step 1: Taxonomic composition of communities

A total of 96 whole-community samples were chosen for taxonomic analysis with two aims: 1) to confirm the variation in flow-cytometric-analysis-based community composition caused by mass transfer, for which samples of inoculum (day 0), for the 2$^{nd}$ and the 8$^{th}$ days as well as the final day for each phase were chosen for all local communities L1-L5 and the regional pool R (starting at day 9); and 2) to confirm the drifts identified by flow-cytometric-analysis-based intra-community variation, for which samples of three time points at which drifts occurred in at least two local communities were chosen (Tables S8.1, S8.2), namely, $RC_{10}$: drift at day 47 in L2 and L5, $RC_{50}$: drift at day 61 in L3 and L5, and Insular II: drift at day 100 in L1 and L3. The samples of day 44 in L2 and L5, day 55 in L3 and L5, and day 97 in L1 and L3 were taken for comparison of community states. The sampling time points for each reactor are listed in Table S9.1.

**Table S9.1** The time points (day) of samples for whole-community taxonomic analysis are listed per reactor and phase. The time points in bold are those used to study the impact of drifts on community variation. The shaded areas represent different phases with changed $RC$. The time points of days 26, 47, 64 and 89 were the days on which the phases ended as well as those on which the next phases started; therefore, drifts at day 47 happened at phase $RC_{10}$, while drifts at day 61 and day 100 happened at $RC_{50}$ and Insular II.

| reactor/ phase | inoculum | Insular I | $RC_{10}$ | $RC_{50}$ | $RC_{80}$ | Insular II |
|---|---|---|---|---|---|---|
| L1 | | 1, 8, 26 | 27, 34, 47 | 48, 55, 64 | 65, 72, 89 | 90, 97, **100**, 107 |
| L2 | | 1, 8, 26 | 27, 34, 44, **47** | 48, 55, 64 | 65, 72, 89 | 90, 97, 107 |
| L3 | 0 | 1, 8, 26 | 27, 34, 47 | 48, 55, **61**, 64 | 65, 72, 89 | 90, 97, **100**, 107 |
| L4 | | 1, 8, 26 | 27, 34, 47 | 48, 55, 64 | 65, 72, 89 | 90, 97, 107 |
| L5 | | 1, 8, 26 | 27, 34, 44, **47** | 48, 55, **61**, 64 | 65, 72, 89 | 90, 97, 107 |
| R | - | 9, 26 | 27, 34, 47 | 48, 55, 64 | 65, 72, 89 | 90, 97, 107 |

The taxonomic composition of communities at genus level is shown in Fig. S9.1 (the relative abundance table of communities at species level is shown in Dataset S9.1). A total of 18 classes and 159 genera were identified over all whole-community samples. Generally, the communities were dominated by the classes Bacteroidia, Gammaproteobacteria and Alphaproteobacteria, which are typical members of activated sludge[16, 17]. At phases Insular I, $RC_{10}$ and Insular II, communities harboured more diverse genera, namely, *Leadbetterella*, *Azospirillum*, *Pseudacidovorax*, *Brevundimonas* and *Pedobacter*. Although the taxonomic compositions of the local communities L1-L5 and the regional pool R were fairly comparable within each of the five phases, the compositions clearly differed between the phases. Therefore, the 16S rRNA gene sequencing data supported the findings obtained by flow cytometry. The 16S rRNA gene sequencing data showed for the $RC_{50}$ and $RC_{80}$ high-rate recycling phases high synchrony in community compositions (Fig. S9.1). An unassigned genus from *Sphingobacteriaceae* (family) accounted for about half of the genera in $RC_{50}$ and $RC_{80}$, and *Leadbetterella* and *Azospirillum* persisted but at lower relative abundances (Fig. S9.1). Members of the *Sphingobacteriaceae* have been reported to degrade a variety of organic chemicals [18-20], indicating their ability to use complex 'leftover' substrates. Members of *Azospirillum* are known for the ability to fix nitrogen[21] and thus could help overcome nitrogen deficiency specifically in $RC_{80}$. These major genera indicated the key role of nutrient availability in community building and it appears that mass transfer supported this dominance.

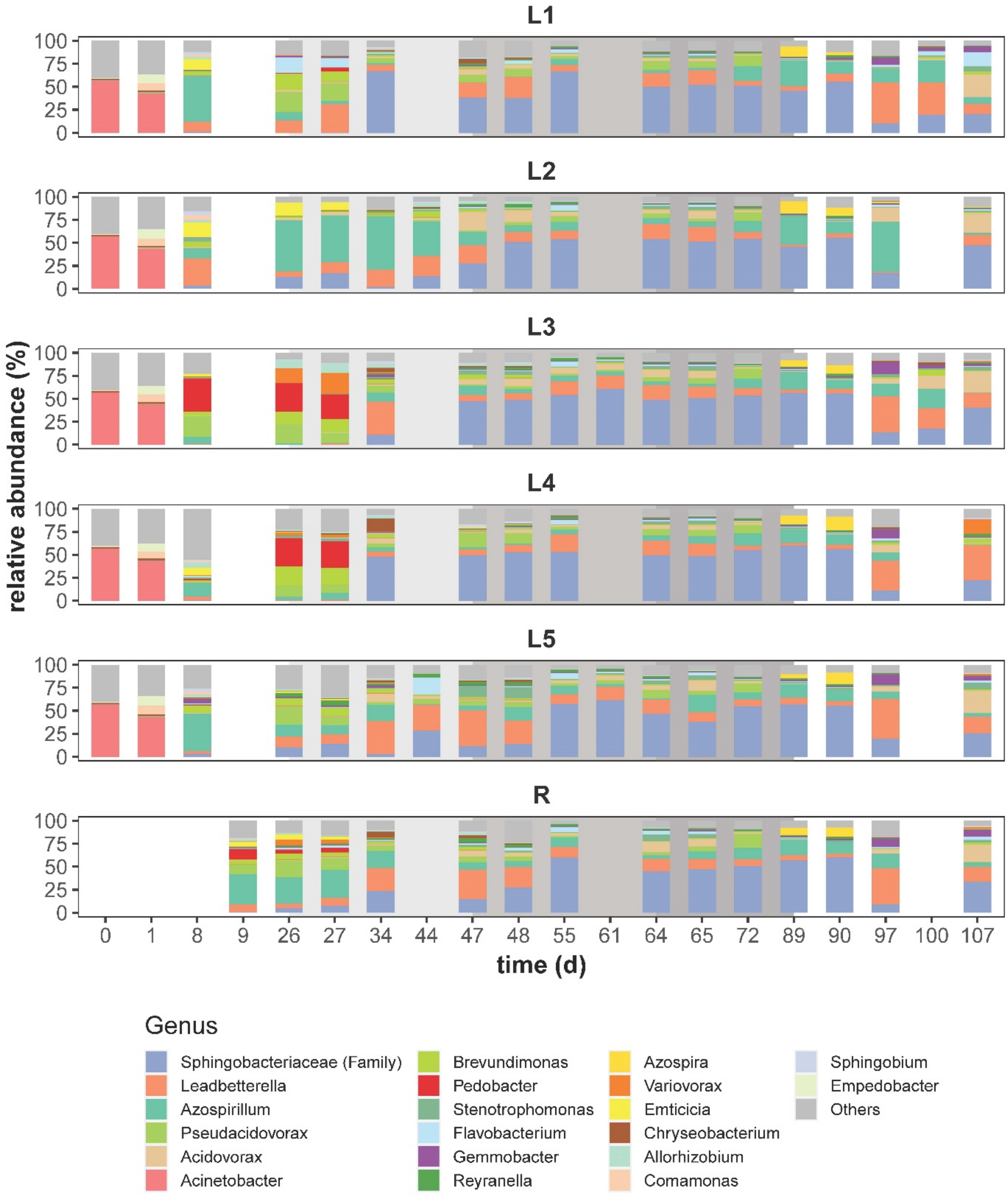


**Figure S9.1** The taxonomic composition in relative abundance (%) of communities over time at the genus level. A total of 159 genera were identified. The top 20 most abundant genera are colour-coded and stacked from bottom to top per column in decreasing order of mean relative abundance over all samples. An unassigned genus from the family *Sphingobacteriaceae* is labelled by this family. The relative abundances of genera other than the top 20 are summed up and represented as ‘Others’. The shaded areas represent different phases with changed $RC$.

The NMDS analysis based on relative abundances of all 159 genera (Fig. S9.2) was supported by the R package ‘vegan’[12] and visualised identically to the procedure used for the flow cytometric single-cell data shown in Fig. 2a. The trend of taxonomic composition of communities confirmed the trend revealed by flow cytometric fingerprints. The most extreme change was observed during the adaptation period of Insular I phase from day 1 to day 8, and all local communities L1-L5 evolved in similar directions at the end of Insular I phase. As $RC$ increased from 10 % to 80 %, the communities became increasingly convergent on both regional and temporal scales. When recycling ended, community divergence increased slightly again compared with the levels at $RC_{50}$ and $RC_{80}$. Thus, the data obtained by 16S gene sequencing clearly evidenced the influence of mass transfer on community composition demonstrated by the flow cytometric data.

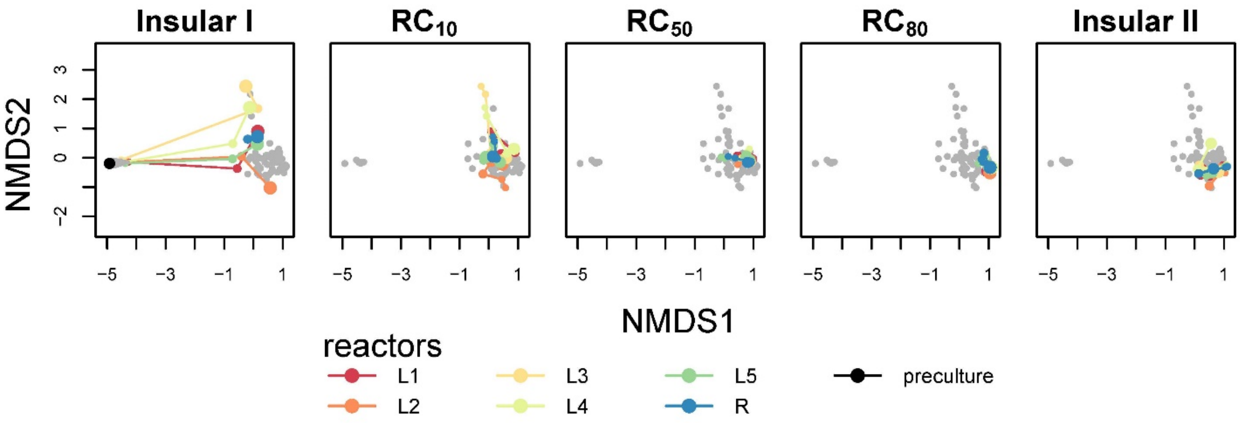


**Figure S9.2** NMDS analysis using relative abundance of all genera (total of 159) based on Bray-Curtis dissimilarities (try = 100, trymax = 200). Successive and connected time points indicate the assembly trajectory of communities. Points in grey represent samples from the other phases.

Additional whole-community samples were analysed at days 47, 61 and 100 to clarify changes in community composition occurring due to drift events previously determined by intra-community β-diversity based on flow cytometric data (Table S9.1). For comparison, the intra-community β-diversity based on the taxonomic composition of communities at the genus level was calculated following the workflow used for flow cytometric data (Supplementary Information S8, step 3). First, all genera with relative abundance > 0.63 (1/159, 1/total number of genera) were determined as ‘dominant genera’. Second, the intra-community β-diversity was computed as the number of

unique dominant genera, which were not shared by pairwise samples from successive time points per community. Third, owing to limited time points sampled for taxonomic analysis, the comparison could not be performed for short time intervals as was done for the much higher available sample numbers of flow cytometric data. Instead, the pairwise comparison could only be undertaken across long time intervals (varying from 1 to 18 days). To create a comparison with these data, the intra-community β-diversity based on SCs was therefore recomputed using the same pairwise samples at chosen time points (Table S9.1, unfilled circles in Fig. S9.3). We found that the variations of intra-community β-diversity based on the two analyses were similar and that the intra-community β-diversity values for both analyses decreased as $RC$ increased, pointing to the fact that increasing $RC$ is a means of preventing stochastic events.
The drifts events were calculated in the balanced periods of all phases. The values based on SCs were often higher than the corresponding values based on genus level, and the latter were so low at some time points that no drift could be detected. These findings showed that some drifts in SCs clearly detected by flow cytometric analysis were not revealed at the genus level. Thus, the β-diversity of SCs was more sensitive as indicator of community variation than the genus level. The background for these findings is that cells of the same genus may occupy different gates (SCs) in the gate template, whereas sequencing does not take into account physiological cell properties.

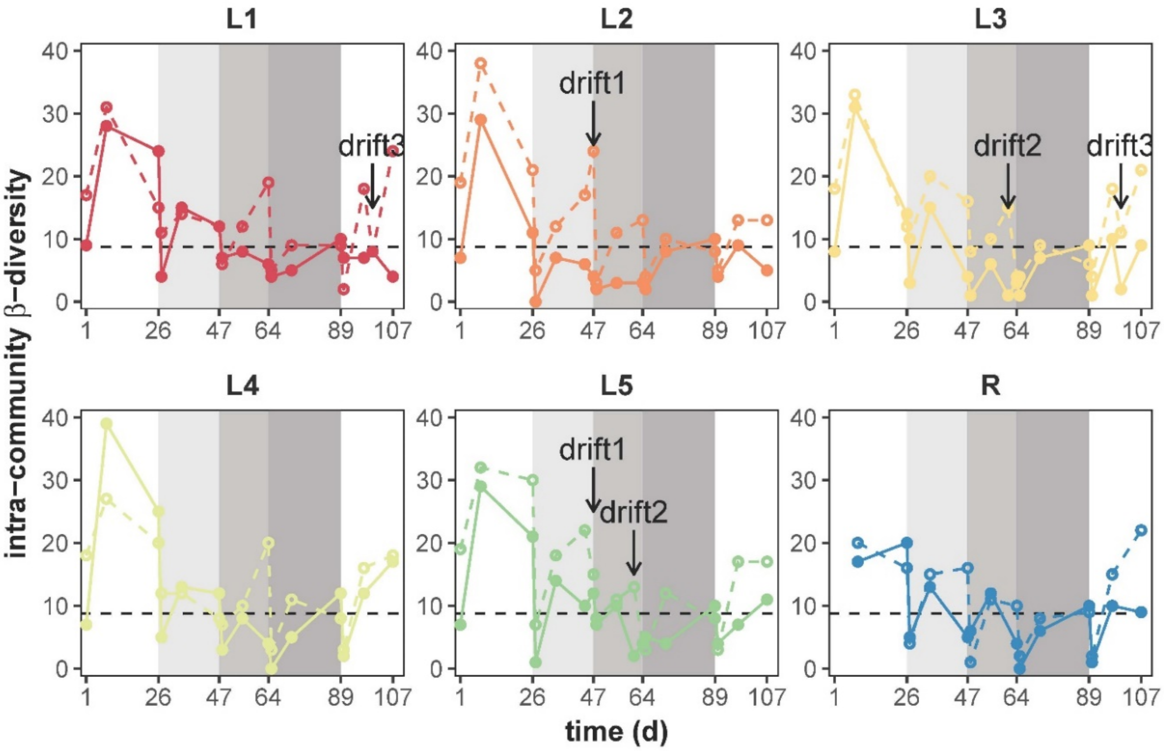


**Figure S9.3** Drift events, detected by intra-community β-diversity variations based on flow cytometric data (SCs, open circles and dashed lines) and based on 16S rRNA amplicon sequencing data (genus, closed circles and closed line). The intra-community β-diversity was calculated as the number of unshared dominant genera (relative abundance $>$ 0.63) or dominant SCs (relative cell abundance $>$ 1.25) by pairwise samples from successive time points per community. The dashed black line marks the threshold determining drift (8.76), and the labels and arrows indicate three drifts from which cells were chosen for 16S rRNA analyses. The following drifts were chosen: drift1: day 47 in L2 and L5, drift2 day 61 in L3 and L5, and drift 3: day 100 in L1 and L3. The shaded areas represent different phases with changed $RC$.

#### Step 2: Taxonomic assignment of sorted SCs

Cells of gates were cytometrically sorted only when their abundance was above 2%. Out of 35,840 SCs a total of 51 SCs were selected for cell sorting and taxonomic analysis based on four specific features: 1) *mass transfer*, to confirm the redistribution of SCs from R to L1-L5 (e.g. G5, G12, G14 and G33) during $RC_{80}$, which were also correlated with $RC$ (Fig. S12.1 and Fig. S8.3); 2) *netgrowth rate*, to investigate the taxonomy of cells in SCs, which grew (i.e. $\mu' SC_x > 0$ in $\geqslant$ 4 reactors) at $RC_{80}$ in L1-L5 (Fig. 4): G1, G5, G9, G13, G18, G25, G33. In addition, we were interested in cells that did not grow but were rescued from R at $RC_{80}$ in L1-L5 (i.e. $\mu' SC_x = 0$ in $\geqslant$ 2 reactors, Fig. 4): G2, G4, G11, G12, G14; 3) *nestedness*, to investigate the taxonomy of cells in the most nested SCs, that is, at $RC_{50}$: G2, G3, G4, G8, G9, G11 and at $RC_{80}$: G5, G12, G14 (Table S8.4); and 4) *drifts*, to investigate the taxonomy of cells benefitting from drifts (drift1: G3, G16, G22, G33, G79; drift2: G5, G7, G12, G18; drift3: G3, G4, G5, G7, G18, G24) or being disadvantaged by drifts (drift1: G1, G7, G24, G26; drift2: G19, G24; drift3: G12, G20, G38; Table S8.2). The sorted SCs are listed according to sample time and reactors in Table S9.2. The position of the sorted SCs in the gate template is shown in Fig. S9.4. The taxonomic composition of sorted SCs is shown at the genus level in Fig. S9.5 (Dataset S9.2). A total of 26 classes and 186 genera were identified over all sorted SC samples. Generally, most sorted SCs were dominated by only one or two classes, namely, Alphaproteobacteria, Gammaproteobacteria, Bacteroidia or Actinobacteria. At the genus level, most SCs were mono-dominant (marked in colour, Table S9.2). Only a few SCs comprised different genera (i.e., 047_G79_L2[4], 061_G5/G12_L3[4], 085_G5_L1/L3/R[1,2,3], 085_G14_L1[1,2,3], 086_G9/G25_L2[2]).

**Table S9.2** Fifty-one cytometrically sorted SCs from respective time points (day) and reactors for taxonomic analysis. Each sample is named as 'time_gate_reactor'. The number labelled as superscript beside the sample name indicates the subjects it involved: 1, mass transfer; 2, netgrowth rate; 3, nestedness; 4, drifts. Colour shades mark SCs dominated by *Azospirillum* …., *Leadbetterella* …., an unassigned genus from *Sphingobacteriaceae* (family) …., an unassigned genus from PeM15 (order) …., *Pseudacidovorax* ….,

*Azospira* …., *Acidovorax* …., *Reyranella* …., *Stenotrophomonas* …., *Brevundimonas* …., *Ochrobactrum* …., *Sphingopyxis* …. and *Microbacterium* …..

| ,time (d) | gate | reactor | | | | | |
|---|---|---|---|---|---|---|---|
| | | L1 | L2 | L3 | L4 | L5 | R |
| 26 | G33 | | 026_G33_L2[1] | | | | |
| 44 | G1 | | 044_G1_L2[4] | | | | |
| 44 | G7 | | | | | 044_G7_L5[4] | |
| 44 | G24 | | 044_G24_L2[4] | | | | |
| 44 | G26 | | 044_G26_L2[4] | | | | |
| 47 | G3 | | | | | 047_G3_L5[4] | |
| 47 | G16 | | 047_G16_L2[4] | | | | |
| 47 | G22 | | 047_G22_L2[4] | | | | |
| 47 | G33 | | 047_G33_L2[4] | | | | |
| 47 | G79 | | 047_G79_L2[4] | | | | |
| 58 | G19 | | | 058_G19_L3[4] | | | |
| 58 | G24 | | | 058_G24_L3[4] | | | |
| 61 | G5 | | | 061_G5_L3[4] | | | |
| 61 | G7 | | | 061_G7_L3[4] | | | |
| 61 | G12 | | | 061_G12_L3[4] | | | |
| 61 | G18 | | | 061_G18_L3[4] | | | |
| 63 | G2 | 063_G2_L1[3] | | | | | |
| 63 | G3 | 063_G3_L1[3] | | | | | |
| 63 | G4 | | | 063_G4_L3[3] | | | |
| 63 | G8 | | | 063_G8_L3[3] | | | |
| 63 | G9 | | | 063_G9_L3[3] | | | |
| 63 | G11 | 063_G11_L1[3] | | | | | |
| 71 | G2 | | | | | 071_G2_L5[2] | |
| 71 | G4 | 071_G4_L1[2] | | | | | |
| 71 | G11 | | | | | 071_G11_L5[2] | |
| 85 | G5 | 085_G5_L1[1,2,3] | | 085_G5_L3[1,2,3] | | | 085_G5_R[1,2,3] |
| 85 | G12 | 085_G12_L1[1,2,3] | | 085_G12_L3[1,2,3] | | | 085_G12_R[1,2,3] |
| 85 | G14 | 085_G14_L1[1,2,3] | | 085_G14_L3[1,2,3] | | | 085_G14_R[1,2,3] |
| 85 | G33 | 085_G33_L1[1,2] | | 085_G33_L3[1,2] | | | 085_G33_R[1,2] |
| 86 | G1 | | 086_G1_L2[2] | | | | |
| 86 | G9 | | 086_G9_L2[2] | | | | |
| 86 | G13 | | | | 086_G13_L4[2] | | |
| 86 | G18 | | 086_G18_L2[2] | | | | |
| 86 | G25 | | 086_G25_L2[2] | | | | |
| 99 | G12 | | | 099_G12_L3[4] | | | |
| 99 | G20 | 099_G20_L1[4] | | | | | |
| 99 | G38 | 099_G38_L1[4] | | | | | |
| 100 | G3 | 100_G3_L1[4] | | | | | |
| 100 | G4 | 100_G4_L1[4] | | | | | |
| 100 | G5 | | | 100_G5_L3[4] | | | |
| 100 | G7 | 100_G7_L1[4] | | | | | |
| 100 | G18 | | | 100_G18_L3[4] | | | |
| 100 | G24 | 100_G24_L1[4] | | | | | |

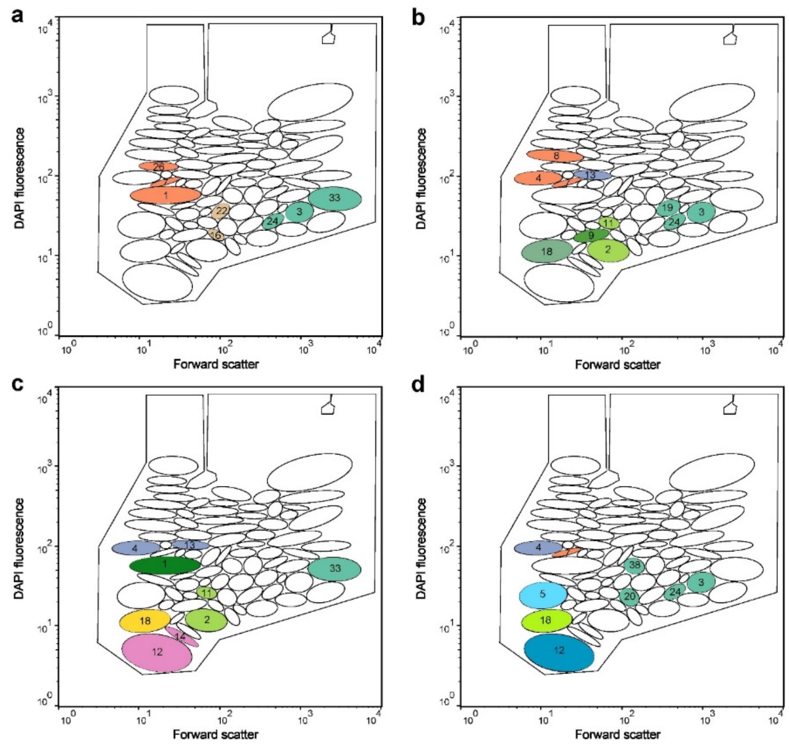


**Figure S9.4** The sorted SCs from a: phase $RC_{10}$ (days 26, 44, and 47), b: $RC_{50}$ (days 58, 61 and 63), c: $RC_{80}$ (days 71, 85 and 86) and d: Insular II (days 99 and 100) are coloured according to their mono-dominance of *Azospirillum* …., *Leadbetterella* …., an unassigned genus from *Sphingobacteriaceae* (family) …., an unassigned genus from PeM15 (order) …., *Pseudacidovorax* …., *Azospira* …., *Acidovorax* …., *Reyranella* …., *Stenotrophomonas* …., *Brevundimonas* …., *Ochrobactrum* …., *Sphingopyxis* …. and *Microbacterium* ….. The numbers (x) inside gates label the corresponding gate name: 'Gx'.

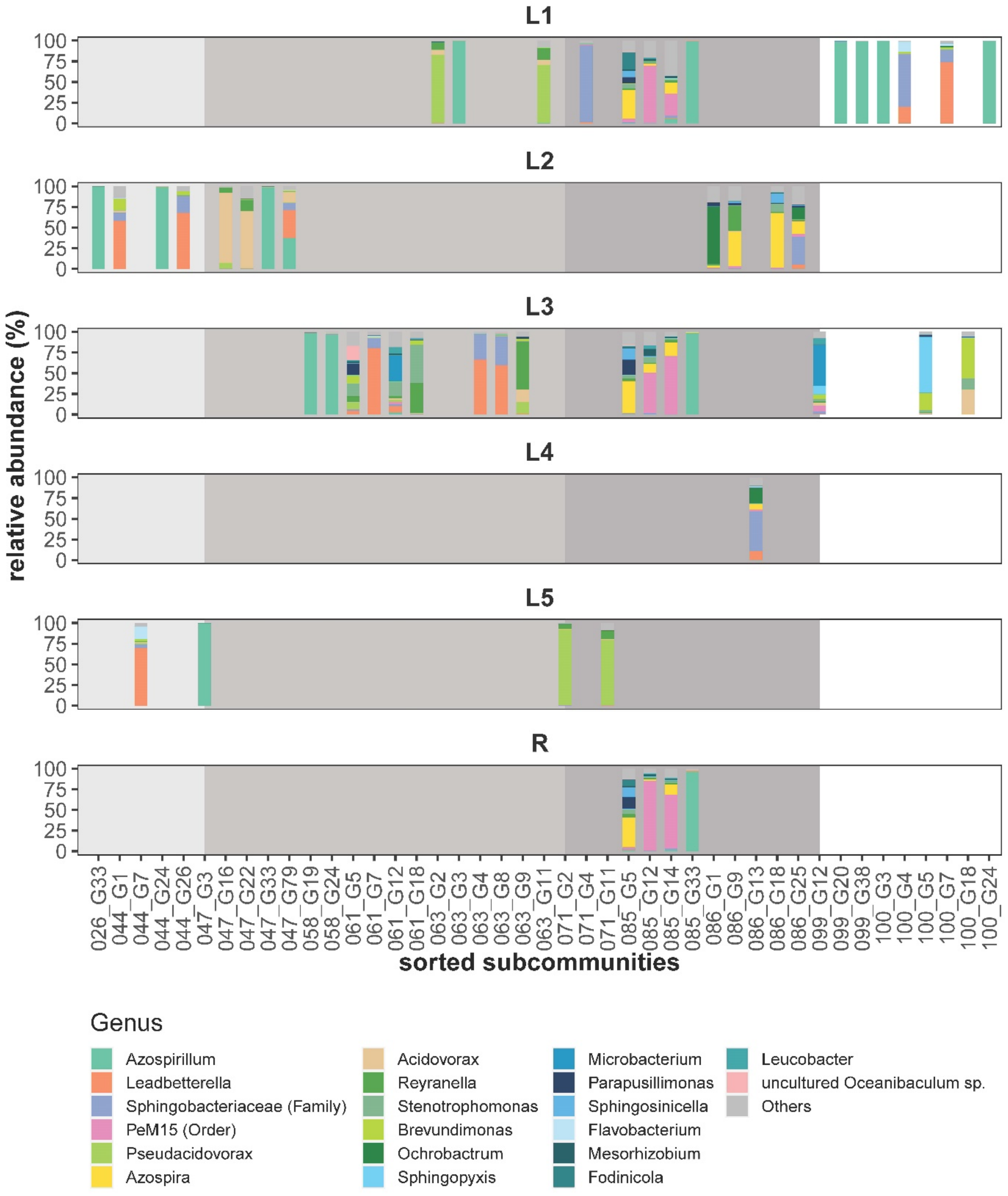


**Figure S9.5** The taxonomic composition in relative abundance (%) of sorted SCs at the genus level. In total, 186 genera were identified. The top 20 most abundant genera are colour-coded and stacked from bottom to top per column in decreasing order of mean relative abundance over all SCs samples. The unassigned genera are labelled with the respective family or order. The relative abundances of genera other than the top 20 are summed up and represented as 'Others'. The shaded areas represent different phases with changed $RC$.

The genus assignments of cells in these SCs are discussed under the four specific features mentioned above:

Mass transfer & netgrowth rate: Mass transfer supported the persistence of SCs in local communities, either by enabling their local growth or by rescuing non-growing SCs by their growth in the regional pool R. For example, at $RC_{80}$, *Azospirillum* (G33), *Azospira* (G9, G18), and *Ochrobactrum* (G1, Fig. S9.5, presenting only SCs > 2% cell abundance) followed the former mechanism ('growth'), and grew also at lower abundance in a set of local communities (Fig. 4), while an unassigned genus from PeM15 (order, G12 and G14), *Sphingobacteriaceae* (family, G4) and *Pseudacidovorax* (G2, G11, Fig. S9.5) followed the latter mechanism ('rescue') and did not grow in local communities in $RC_{80}$, but were rescued by growth in the regional pool R (Fig. 4). The mono-dominant SCs that showed netgrowth and that were rescued can be assumed to be superior local competitors. Some SCs that are also among the growing SCs (G5, G9, G25, Table S9.2) were multi-dominant and it can be assumed that the genera in each of the SCs can cooperate under $RC_{80}$ conditions.

Nestedness: Generally, nested SCs were those that persisted in the different phases of the 80-generation experiment. For phase $RC_{50}$, we found for the flow cytometrically determined nestedness the SCs G2, G3, G4, G8, G9, and G11 (Table S8.4). Nearly the same gates were calculated as nested ones from the 16S rRNA sequencing data: genera *Azospirillum* (G3), *Leadbetterella* (G4, G8), *Pseudacidovorax* (G2, G11) and *Reyranella* (G9).

In $RC_{80,}$ flow cytometrically determined nested SCs were G2, G4, G5, G9, G12, and G14 (Table S8.4). Sequencing data found PeM15 (order, G12, G14) and *Pseudacidovorax* (G2) nested, while G5 and G9 were mostly multi-dominant and changed their proportions of contained genera from $RC_{50}$ to $RC_{80}$.

In general, mass transfer supported genera such as G2, G12 and G14, which were nested and rescued by their ability to accumulate in the regional pool R (e.g., the unassigned genus from PeM15).

Drifts: Drifts are of stochastic in nature when growth conditions are balanced. Three drift events occurring during 80 generations of cultivation were taxonomically investigated (Table S9.1). We found a flow cytometrically determined drift1 in $RC_{10}$ where G1, G7, G24 and G26 decreased and G3, G16, G22, G33 and G79 increased in cell abundance (Table S8.2). 16S rRNA sequencing revealed the loss of *Leadbetterella* (G1, G7, G26) and the gain of *Acidovorax* (G16, G22).

Meanwhile, *Azospirillum* was lost from G24 and reappeared in G3 and G33. Drift2 ($RC_{50}$, day 61) showed cytometrically determined gates with decreased (G19, G24) and increased cell abundance (G5, G7, G12, G18). 16S rRNA sequencing revealed the loss of *Azospirillum* (G19, G24) and the gain of *Leadbetterella* (G7) and *Stenotrophomonas* (G18). Drift3 (Insular II, day 100) showed cytometrically determined gates with decreased (G12, G20, G38) and increased cell abundance (G3, G4, G5, G7, G18, G24). 16S rRNA sequencing revealed the loss of *Microbacterium* (G12) and the gain of an unassigned genus from *Sphingobacteriaceae* (G4), *Sphingopyxis* (G5), *Leadbetterella* (G7) and *Brevundimonas* (G18). *Azospirillum* was lost from G20 and G38 and was gained in G3 and G24.

*Azospirillum* showed its physiological flexibility by switching between different SCs, probably due to cell cycling states[22], instead of going extinct. Other genera also showed that flexibility. These findings support the lower intra-community β-diversity, calculated at the genus level, in comparison to the SCs calculated based on flow cytometric data (Fig. S9.3). This flexibility of some of the dominant genera that were also nested and that showed netgrowth or were rescued by mass transfer appears to have abolished the stochastic events occurring in the balanced periods under $RC$.

#### S10: Calculation of the netgrowth rate µ’

In this study, the biomass was quantified by counting cell numbers in local communities L1-L5 and the regional pool R (Fig. S5.2). The change of the total cell number during an interval $\Delta t$ (d) was determined by cell numbers coming from the influent, cell numbers lost by the effluent and the netgrowth rate µ’ of cells within $\Delta t$ (d).

$$Change = Netgrowth + Influent - Effluent \qquad \text{Eq. S10.1}$$

The reactors of the five local communities L1-L5 and the regional pool R were fully mixed. Therefore, the cell numbers of the influent into local communities L1-L5 and into the regional pool R ($influent_{Lin}$ or $influent_{Rin}$) were equal to the cell numbers of the reactors from which the influent was pumped. In addition, the cell numbers in the effluent ($effluent_{Lout}$ or $effluent_{Rout}$) were equal to the cell numbers in that same reactor (L1-L5 or R). The netgrowth rate $\mu'$ of cells was calculated by the following equations:

$$\Delta n_{L,R} \cdot V_{L,R} = \mu' \cdot V_{L,R} \cdot n_{L,R} \cdot \Delta t + Q_{in} \cdot n_{in} \cdot \Delta t - Q_{eff} \cdot n_{eff} \cdot \Delta t \qquad \text{Eq. S10.2}$$

$$\mu' = \frac{\Delta n_{L,R}}{\Delta t \cdot n_{L,R}} - \frac{Q_{in} \cdot n_{in}}{V_{L,R} \cdot n_{L,R}} + \frac{Q_{eff} \cdot n_{eff}}{V_{L,R} \cdot n_{L,R}} \qquad \text{Eq. S10.3}$$

with:

| symbol | description |
|---|---|
| $n_{L,R}$, $n_{in}$ & $n_{eff}$ | cell numbers (cells mL$^{-1}$) per reactor [local communities ($n_L$) and regional pool ($n_R$)], per influent ($n_{in}$) or per effluent ( $n_{eff}$) |
| $\Delta n_{L,R}$ | difference of cell numbers (cells mL$^{-1}$) between successive samples in the respective reactors during an interval $\Delta t$ (d) |
| $\overline{n_{L,R}}$ | average of cell numbers (cells mL$^{-1}$) between successive samples in the respective reactors during an interval $\Delta t$ (d) |
| $V_{L,R}$ | working volume (mL) per reactor |
| $Q_{in}$ | rate of influent volume per reactor (i.e., influent flow rate, mL d$^{-1}$), which is manually adjusted per phase in this study (Table S2.1) |
| $Q_{eff}$ | rate of effluent volume per reactor (i.e., effluent flow rate, mL d$^{-1}$) |
| $\mu'$ | netgrowth rate (d$^{-1}$) of cells in a local communities L1-L5 or the regional pool R |

The difference in cell numbers ($\Delta n_{L,R}$) between successive samples on day $i$ and day $i + \Delta t$ was determined during interval $\Delta t$ (Eq. S10.4). The average of cell numbers $\overline{n_{L,R}}$ based on the cell numbers of successive samples and per day were calculated (Eq. S10.5) and used for the calculation of μ' during the interval $\Delta t$. This was performed for the cell numbers in local communities L1-L5 as well as the regional pool R ($n_{L,R}$).

$$\Delta n_{L,R} = n_{L,R\ at\ i+\Delta t} - n_{L,R\ at\ i} \qquad \text{Eq. S10.4}$$

$$\overline{n_{L,R}} = (n_{L,R,at\ i} + n_{L,R,at\ i+\Delta t})/2 \qquad \text{Eq. S10.5}$$

#### Calculation of the recycling rate $\boldsymbol{RC}$

Calculation of the recycling rate $RC$ for the local communities was based on the given dilution rate $D = Q_{eff} / V_{L,R}$. In this study, $D$ was 0.72 d$^{-1}$. With fixed working volume $V_{L,R}$, $Q_{eff}$ was balanced with the sum of the flow rates of the influent$_{\text{Lin}}$ from the regional pool R ($Q_{in}$) and the influent from the medium ($Q_{medium}$). The influent$_{Lin}$ from the regional pool R ($Q_{in}$) contained cells. On each day, 576 mL was exchanged per reactor from which, during times of mass transfer, 10, 50, or 80 was taken from the regional pool R (Table S2.1). In the local communities L1-L5, with a given $D$ and a given effluent$_{Lout}$ flow rate $Q_{eff}$, the influent$_{Lin}$ flow rate $Q_{in}$was dependent on the recycling rate $RC$.

$$Q_{in} = RC \cdot Q_{eff} \qquad \text{Eq. S10.6}$$

#### Relationship between netgrowth rate μ' and mass transfer rate ***M***

In completely mixed reactors, $n_{eff}$ was considered equal to $n_{L,R}$, and for a local communities L1-L5, $n_{in}$ was considered equal to $n_R$. For a local communities, Eq. S10.3 was simplified:

$$\mu' = \frac{\Delta n_L}{\Delta t \cdot n_L} - D \cdot RC \cdot \frac{n_R}{n_L} + D \qquad \text{Eq. S10.7}$$

The mass transfer rate $M$ was quantified by the daily cell number entering the local community ($Q_{in} \cdot n_{in}$) in relation to the cell number present in the respective local community ($V_L \cdot n_L$):

$$M = \frac{Q_{in} \cdot n_{in}}{V_L \cdot n_L} = D \cdot RC \cdot \frac{n_R}{n_L} \qquad \text{Eq. S10.8}$$

By this, $M$ was determined by the dilution rate and the recycling rate ($D \cdot RC$) and the relative difference of cell numbers between the regional pool and respective local community ($\frac{n_R}{n_L}$).

Then, Eq. S10.7 was replaced as:

$$\mu' = \frac{\Delta n_L}{\Delta t \cdot n_L} + (D - M) \qquad \text{Eq. S10.9}$$

The netgrowth rate $\mu'$ is thus dependent on the difference between dilution rate $D$ and mass transfer rate $M$, which characterises the influence of emigration or immigration and the relative increase or decrease in cell numbers $\frac{\Delta n_L}{\Delta t \cdot n_L}$.

The ideal relationship among $\mu'$, $D$ and $M$ is shown for a local community in the case of an increase of $RCs$ (Fig. S10.1). In an ideal relationship, the netgrowth rate $\mu'$ decreases when the mass transfer rate $M$ increases at a given $D$ (from $M1$ to $M3$). Under these circumstances, the increase in $M$ would initially cause an increase in cell numbers (Fig. S10.2). After adaptation and when $\mu'$ is equal to the newly adjusted $(D - M)$ under balanced growth conditions [i.e. $\mu'$ –

$(D - M) = 0$], a successive phase-to-phase decrease of $\mu'$ will occur with further increase of $M$. Instead, if there is no $M$ and $D$ remains unchanged, the cell numbers will remain stable with $\mu' = D$ after an adaptation period. If $M$ is stopped after periods of various mass transfer rates, the netgrowth rate will equal $D$ again ($\mu' = D$), which will also lead to a decrease in cell numbers to their original values (Fig. S10.2). In comparison to hypothesised patterns of cell number change (Fig. S10.2), the experimentally counted cell numbers (Fig. S5.2) showed a less clear pattern at phases 1-3. However at $RC_{80}$ and Insular II, cell number changed drastically in the adaptation periods while fluctuating around a plateau in the balanced periods, which fitted our hypothesis (Fig. S5.2).

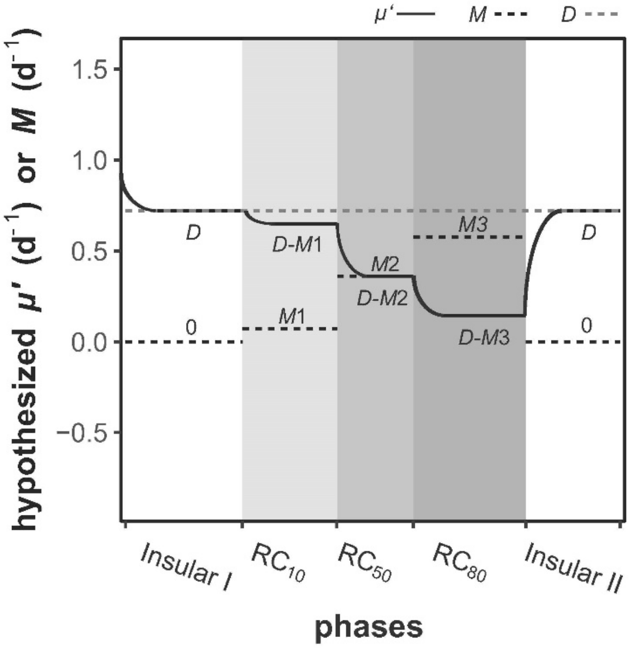


**Figure S10.1** Hypothesised ideal dynamics of the netgrowth rate $\mu'$ **(**the black line) and the mass transfer rate $M$ (dotted black line) after increases in recycling rates $RC$ in a local community. $D$: dilution rate (dashed grey line). Under ideal balanced conditions with unvarying cell numbers ($\Delta n_L = 0$) and if the cell number of the influent from the regional pool R ($n_R$) is equal to the cell number in the respective local community ($n_L$), then the transfer rate is $M = D \cdot RC$ and the netgrowth rate $\mu' = D - M$. $M$ can be calculated for known $RC$ values: $M1 = 0.1D$ ($RC$ 10), $M2 = 0.5D$ ($RC$ 50), $M3 = 0.8D$ ($RC$ 80). The shaded areas represent different phases with changed $RC$.

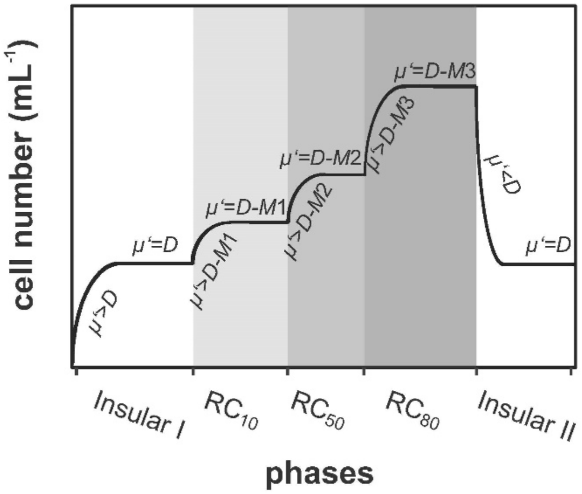


**Figure S10.2** Hypothesised ideal dynamics of the cell number (cells $mL^{-1}$) in a local community after increases in recycling rates $RC$. $\mu'$ : netgrowth rate, $D$: dilution rate, $M$: mass transfer rate. The cell numbers increase during the adaptation period when an increase in the mass transfer rate ($M$) causes an imbalance with $\mu' > D - M$. The netgrowth rate $\mu'$ will equal $(D - M)$ after the adjustment to balanced conditions $\mu' = D - M$. Under balanced conditions, cell numbers do not change ($\Delta n_L = 0$). The shaded areas represent different phases with changed $RC$.

In our study, the following ideal values for $M$ were expected for the proposed experimental setup when conditions return to balanced situations $(\Delta n_L = 0)$ and cell numbers of $\text{influent}_{\text{Lin}}$ (i.e. regional pool R) are equal to cell numbers in the local community $(n_R = n_L)$. With changing $RCs$, $M$ will also change, such as for $M1 = 0.1D = 0.072$ $d^{-1}$ ($RC_{10}$); $M2 = 0.5D = 0.36$ $d^{-1}$ ($RC_{50}$) and $M3 = 0.8D = 0.576$ $d^{-1}$ ($RC_{80}$). Corresponding to Eq. S8.9, $\mu' = D - M$, the netgrowth rate can theoretically be calculated to be 0.648 $d^{-1}$ ($RC_{10}$), 0.36 $d^{-1}$ ($RC_{50}$) and 0.144 $d^{-1}$ ($RC_{80}$). For comparison, the average values of experimental $M$ of L1-L5 were for $RC_{10}$ 0.06 ± 0.02 $d^{-1}$; for $RC_{50}$ 0.30 ± 0.06 $d^{-1}$; and for $RC_{80}$ 0.59 ± 0.05 $d^{-1}$, while the average values of calculated $\mu'$ were 0.69 ± 0.27 $d^{-1}$, 0.51 ± 0.42 $d^{-1}$ and 0.20 ± 0.18 $d^{-1}$ (Table S10.1, Dataset S10). Thus, the experimental $M$ and $\mu'$ values nearly reached the hypothesised values of Fig. S10.1.

#### Produced cell numbers in local communities L1-L5

The absolute cell numbers increased from 1.28 ± 0.74 $\times 10^{12}$ cells per 800 mL in Insular I phase to 4.23 ± 1.11 $\times 10^{12}$ per 800 mL in $RC_{80}$ (Table S10.1). To determine whether this increase in cell number was caused by mass transfer or by real cell production within reactors, the production of individual cells $(\text{PC})$ per reactor per day (cells $d^{-1}$) was calculated. For this calculation, the netgrowth rate μ′ per reactor was used. In addition, the term $\overline{n_{L,R}}$ was considered as the mean of the cell numbers in the same period for which μ’ was calculated.

$$\mathrm{PC} = \mu' \times V_{R,L} \times \overline{n_{L,R}} \qquad \text{Eq. S10.10}$$

The increase in $M$ and the decrease in μ′ in recycling phases $RC_{10}$-$RC_{80}$ did not markedly influence the number of produced cells per day and reactor (PC). Only in $RC_{80}$ was the PC value slightly lower. The data indicate that nutrients placed no limitation on growth in most phases. Only the ammonium concentration decreased to lower values in $RC_{80}$ and also in R (Fig. S3.1), which supports the lowered PC for $RC_{80}$. The values of PC, $M$ and μ′ are shown in Table S10.1 and Dataset S10.

Table S10.1 Summary of experimental M, μ’ and PC values per day and at reactors L1-L5. The mean ± sd values were all calculated among local communities L1-L5 during balanced periods. All negative values were set to zero before averaging.

| comparison L1-L5 | Insular I | $RC_{10}$ | $RC_{50}$ | $RC_{80}$ | Insular II |
|---|---|---|---|---|---|
| experimental M ($d^{-1}$) | 0 | 0.06 ± 0.02 | 0.30 ± 0.06 | 0.59 ± 0.05 | 0 |
| experimental μ’ ($d^{-1}$) | 0. 71 ± 0.27 | 0.69 ± 0.27 | 0.51 ± 0.42 | 0.20 ± 0.18 | 0. 79 ± 0.31 |
| cell numbers per reactor (800 mL, $\times 10^{12}$ cells $mL^{-1}$) | 1.28 ± 0.74 | 1.46 ± 0.52 | 1.97 ± 0.57 | 4.23 ± 1.11 | 1.26 ± 0.50 |
| PC per reactor (800 mL, $\times 10^{12}$ cells $d^{-1}$) | 0.94 ± 0.56 | 0.99 ± 0.49 | 1.01 ± 0.74 | 0.85 ± 0.72 | 1.01 ± 40.49 |

#### S11: Quantification of the netgrowth rate per subcommunity μ′ $SC_x$

Similar to the community netgrowth rate (μ′, Supplementary Information S10), the netgrowth rate was calculated for each SC. At the SC-level, the netgrowth rate ($\mu' SC_x$) of each SC per phase and reactor and time $\Delta t$ was calculated only for the balanced growth phases (Dataset S11). Overall, 2400 SCs were evaluated for each of the local and regional communities and per phase. In each phase, if an SC was never dominant ($\leqslant$ 1.25% relative cell abundance per SC), it was excluded from the evaluation. The criterion $\mu' SC_x \geqslant 0$ was chosen to differentiate the various degrees of netgrowth $\mu' SC_x$ of cells in the SCs (Fig. 11.1d). For the determination of $\mu' SC$, the absolute cell abundance per SC was used (Fig. S8.2).

The change in cell numbers per SC during interval Δt (d) was determined by cell numbers per SC originating from the influent, cell numbers per SC lost by the effluent and the netgrowth rate $\mu' SC$ of the cells per SC within Δt (d). According to the biomass balance shown in Eq. S11.1, the $\mu' SC$ can be calculated similar to $\mu'$ using Eq. S11.2 by specifying the values for $SCx$ (x = 1-80 for 80 SCs in total):

$$\Delta nSC_{x,L,R} \cdot V_{L,R} = \mu' SC_x \cdot V_{L,R} \cdot nSC_{x,L,R} \cdot \Delta t + Q_{in} \cdot nSC_{x,in} \cdot \Delta t - Q_{eff} \cdot nSC_{x,eff} \cdot \Delta t$$

Eq. S11.1

with:

| symbol | description |
|---|---|
| $nSC_{x,L,R}$, $nSC_{x,in}$ & $nSC_{x,eff}$ | absolute cell abundances in $SC_x$ (cells $mL^{-1}$) per reactor [local communities ($nSC_{x,L}$) and regional pool ($nSC_{x,R}$)], per influent ($nSC_{x,in}$) or per effluent ($nSC_{x,eff}$). |
| $\Delta nSC_{x,L,R}$ | difference of absolute cell abundances in $SC_x$ (cells $mL^{-1}$) between successive samples in the respective reactors during interval $\Delta t$ (d) |
| $\overline{nSC_{x,L,R}}$ | average of absolute cell abundances in $SC_x$ (cells $mL^{-1}$) between successive samples in the respective reactors during interval $\Delta t$ (d) |
| $\mu' SC_x$ | the netgrowth rate ($d^{-1}$) of cells in $SC_x$ in local communities L1-L5 or regional pool R |

The difference in cell numbers per $SC_x$ ($\Delta nSC_{x,L,R}$) between successive samples was determined for the interval $\Delta t$ (Eq. S11.2). The average cell numbers per $SC_x$ ($\overline{nSC_{x,L,R}}$) based on the cell numbers per SC of successive samples and per day were calculated (Eq. S11.3) and used for the calculation of $\mu' SC_x$ for interval $\Delta t$. This was done for the cell numbers of $SC_x$ in local communities L1-L5 and in the regional pool R ($nSC_{x,L,R}$).

$$\Delta nSC_{x,L,R} = nSC_{x,L,R\ at\ i+\Delta t} - nSC_{x,L,R\ at\ i} \qquad \text{Eq. S11.2}$$

$$\overline{nSC_{x,L,R}} = (nSC_{x,L,R\ at\ i} + nSC_{x,L,R\ at\ i+\Delta t})/2 \qquad \text{Eq. S11.3}$$

For a simplified determination of $\mu' SC_x$ in the local communities L1-L5, we assumed that the absolute cell abundance in the $\text{effluent}_{\text{Lout}}$ $nSC_{x,eff}$ is equal to the absolute cell abundance of the local communities L1-L5 $nSC_{x,L}$, and the absolute cell abundance in the $\text{influent}_{\text{Lin}}$ $nSC_{x,in}$ is equal to absolute cell abundance in the regional pool R $nSC_{x,R}$. For $SCx$ in a local community, Eq. S11.1 was simplified:

$$\mu' SC_x = \frac{\Delta nSC_{x,L}}{\Delta t \cdot nSC_{x,L}} - D \cdot RC \cdot \frac{nSC_{x,R}}{nSC_{x,L}} + D \qquad \text{Eq. S11.4}$$

For a simplified determination of $\mu' SC_x$ in the regional pool R, we assumed that the absolute cell abundance in the effluent$_{\text{Rout}}$ $nSC_{x,eff}$ is equal to the absolute cell abundance of the regional pool R $nSC_{x,R}$, and the absolute cell abundance in the influent$_{\text{Rin}}$ $nSC_{x,in}$ is equal to the absolute cell abundance in the local communities $nSC_{x,L}$. To calculate the cell numbers per SC entering the regional pool R $(nSC_{x,in})$ from five local communities (see setup of reactor in Fig. 1), the average of the cell numbers of $SC_x$ $[SUM(nSC_{x,L})/5]$ was used. Therefore, from Eq. S11.1, the calculation of $\mu' SC_x$ in the regional pool R was simplified:

$$\mu' SC_x = \frac{\Delta(nSC_{x,R})}{\Delta t \cdot nSC_{x,R}} - 5D \cdot \frac{SUM(nSC_{x,L})}{5 \cdot nSC_{x,R}} + 5D \qquad \text{Eq. S11.5}$$

#### Calculation of $\mu'$ $\boldsymbol{SC_x}$ on the basis of experimental data

In our study, $\mu' SC_x$ was determined on the basis of cell numbers per $SC_x$. The cell number per $SC_x$ was measured under the influence of the mass transfer at varying $RCs$ on a daily basis. The balanced periods of the different phases were chosen as $\Delta t$. For the calculation of $\mu' SC_x$ over $\Delta t$, the starting days $i$ (days 7, 33, 54, 71, 96) and ending days $i + \Delta t$ (days 26, 47, 64, 89, 107) were chosen. The detailed values for $\mu' SC_x$ per $SC_x$ during the balanced periods of the different phases are presented for the local communities and the regional pool in Dataset S11. For each phase, $\mu' SC_x$ (x = 1-80) was calculated for the dominant SCs (relative cell abundance > 1.25 in at least one sample during the corresponding periods) in L1-L5 and R (Fig. 4).

The $\mu' SC_x$ of most of the SCs showed positive growth in the local communities L1-L5. Others showed zero growth (blue dots in Fig. 4). Among those that showed zero growth were also those that were negative. The negative values originated from huge variations in cell numbers in the local communities L1-L5 compared to the relatively unchanged ones from the regional pool R (Table S5.1). To prevent an overestimation, we set all seemingly negative growth rates to zero.

Similar to μ' for the whole-community, $\mu' SC_x$ showed a decreasing trend with increased $RC$ (Fig. 4). During phases of insular growth, $\mu' SC_x$ must have been at least equal to $D$ in local communities; otherwise, SCs with $\mu' SC_x < D$ would run the risk of being washed out. Once the mass transfer $M$ had diminished the danger of a wash out due to back-cycling of cells, $\mu' SC_x$ was adjusted to the actual $D - M$. Thus, cells in $SC_x$ were present in the system with a $\mu' SC_x$ smaller than $D$. In the regional pool R, there was no influent of fresh medium, and most SCs showed no netgrowth ($\mu' SC_x$, blue circles in Fig.4) due to severe limitations in nutrients (e.g. ammonium decreased from 27.18 ± 11.22 mg N L$^{-1}$ in local communities to 12.46 ± 8.50 mg N L$^{-1}$ in the regional pool R). Cells entered R at high rates from the local communities L1-L5 where the nutrients were already limited. Consequently, there was minimal capacity for growth.

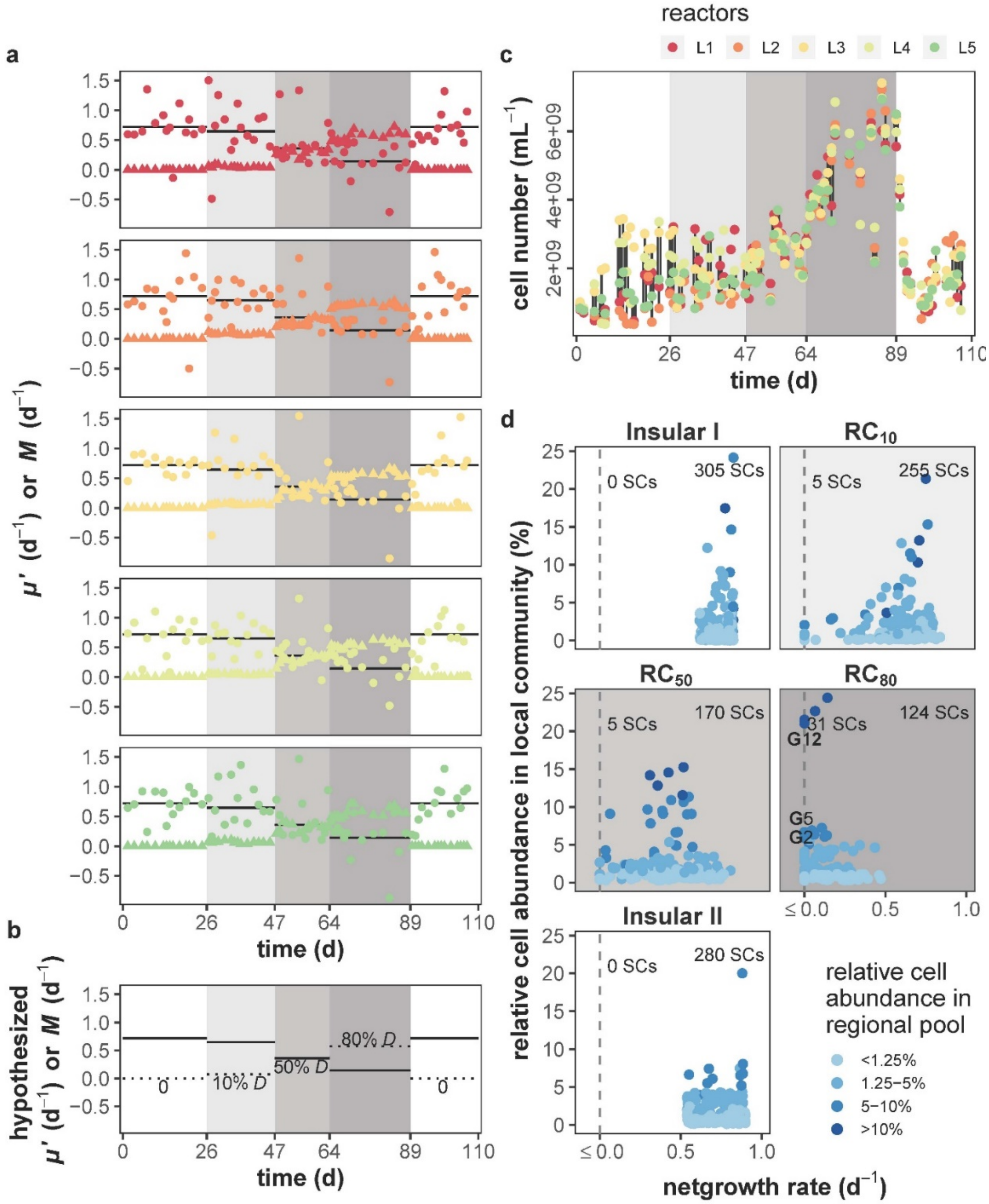

**Figure S11.1** Netgrowth rates of microbiomes and subcommunities during increasing recycling rates $RC$ and mass transfer rates $M$. **a)** Netgrowth rate $\mu'$ (closed circles) and mass transfer rate $M$ (closed triangle) of the local microbiomes L1-L5. The hypothesised values for $\mu'$ (black line) are also shown. **b)** Hypothetical $M$ (black dotted line) and $\mu'$ (black line) values calculated for the experimental setup. **c)** Absolute cell numbers for each of L1-L5. **d)** Netgrowth rates of all dominant SCs ($\mu' SC_x$) of L1-L5. Each point stands for one SC. Heights of points on the y-axis indicate relative average cell abundance per SCs during the balanced period per local community. The blue shades show the relative average cell abundance of the same SC in the regional pool R. The vertical grey dashed line indicates $\mu' SC_x = 0$. The numbers of dominant SCs with $\mu' SC_x = 0$ and $\mu' SC_x > 0$ are marked. With $\mu' SC_x = 0$, G12, G5 and G2 were the most abundant ones (relative average cell abundance $>$ 5%) in L1-L5 at $RC_{80}$. The grey shades in the background of all graphs indicate the five phases Insular I phase, $RC_{10}$, $RC_{50}$, $RC_{80}$ and Insular II phase.

#### S12: Effect of the recycling rate $RC$ on the presence of SC

To determine the influence of the recycling rate $RC$ on absolute cell abundance per SCs (cell number, $mL^{-1}$), the R package 'Hmisc'[23] and the Pearson's correlation test were used. All SCs from local communities L1-L5 and balanced periods were tested (Fig. S12.1). The test revealed a total of five SCs, which increased their absolute cell numbers to at least $10^8$ cell $mL^{-1}$ with increasing $RC$ and showed a high correlation coefficient IrhoI $\geqslant$ 0.55, $p \leqslant 0.05$. Theses gates were G5, G12, G13, G14 and G33. All of them were shown to be involved in netgrowth and rescue under mass transfer (Supplementary Information S11, step 2).

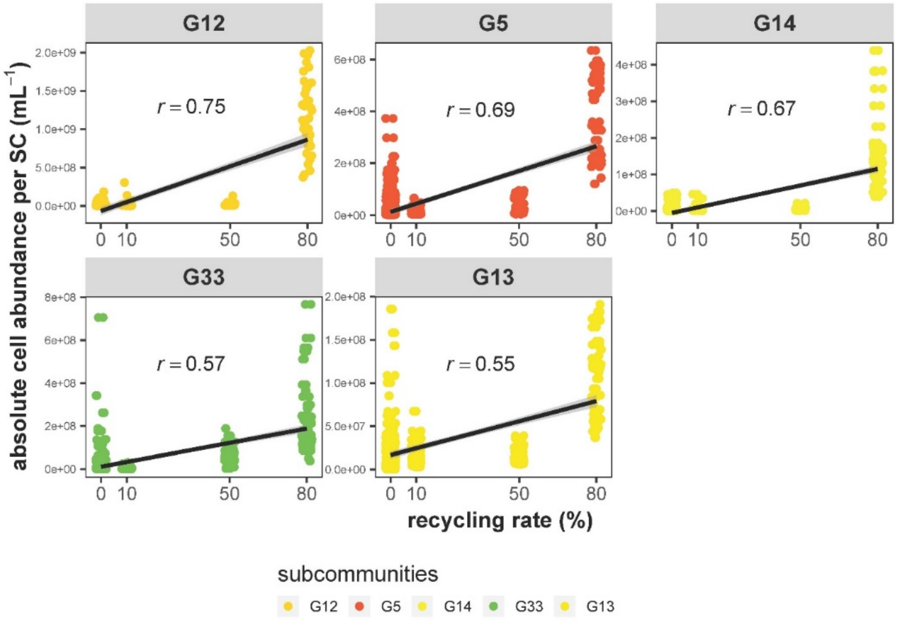

**Figure S12.1** The correlations between absolute cell abundance per SC and $RC$. Only samples from the balanced period were used for analysis, and only correlations with a Pearson's correlation coefficient ($r$) $>$ 0.55 and SCs with absolute cell numbers increased to at least $10^8$ cell $mL^{-1}$ are listed ($p \leqslant 0.05$). The line in each sub-plot indicates linear fitting, while the *r* value represents Pearson's correlation coefficient. The colours indicate the SCs.

In addition, the strength of the regional pool R to shape local communities L1-L5 in their different phases via mass transfer was tested for all communities including all SCs (from both the adaptation and balanced periods, Fig. S12.2). The data show that, as $RC$increased (particularly at $RC_{80}$), fewer SCs responded to mass transfer, but they did so with higher relative abundance per SC. At the same time, SCs tended to have similar relative cell abundances between L1-L5 and R. This is another indication that mass transfer synchronised L1-L5 via R. At Insular I and II phases, there was no correlation of cell abundance per SC between L1-L5 and R and an increase in the variety of dominant SCs. Clearly, the regional pool R shaped the local communities L1-L5.

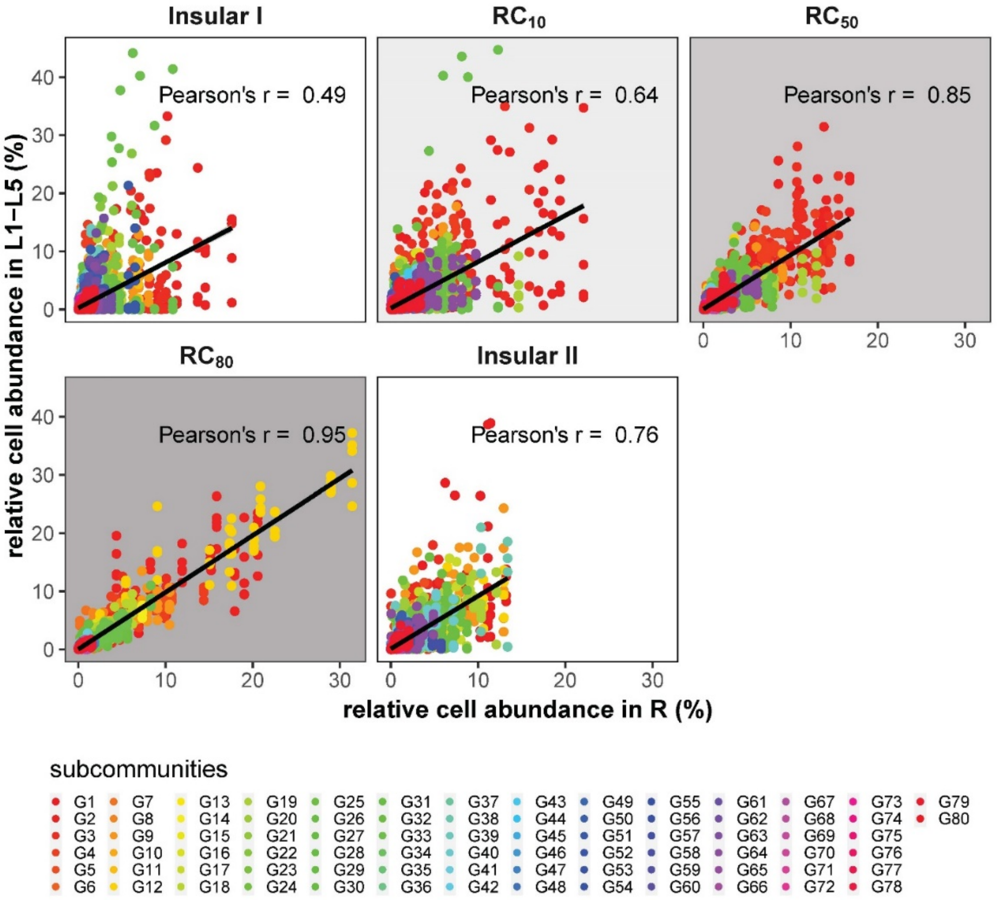

**Figure S12.2** Relationship between relative cell abundance per SC in local communities L1-L5 and the regional pool R. The line in each sub-plot indicates linear fitting, labelled with Pearson's correlation coefficient r. The shaded areas represent different phases with changed $RC$.

#### S13: Relationship between biotic and abiotic parameters

To analyse whether operational reactor conditions influenced the communities structures over 80 generations, the correlations of SCs versus biotic or abiotic parameters (SC vs. Para) but also between SCs (SC vs. SC) were tested. Spearman's rank order correlation coefficient was used for this purpose. For this analysis, the relative cell abundances of SCs from both the adaptation and the balanced periods were used. The following biotic and abiotic parameters were included in the analysis: pH, electrical conductivity (EC), ammonium, total phosphate (PHOt), total chemical oxygen demand (CODt), supernatant chemical oxygen demand (CODs), biological chemical oxygen demand (CODb), OD and cell number (Fig. S3.1). The lists of all strong correlations (Spearman's rank order correlation coefficient $|rho| \geqslant 0.75$, $p \leqslant 0.05$, corrected according to the Benjamini-Hochberg method[24]) are given as Dataset S13.1 for SC vs. SC correlations and Dataset S13.2 for SC vs. Para correlations. The numbers of those correlations are summarised in Table S13.1. Both analyses were performed with the R package 'Hmisc'[23].

**Table S13.1** The numbers of significant correlations of SC vs. SC and SC vs. Para are summarised for the local communities L1-L5 and the regional pool R per phase. Numbers of correlations in L1-L5 are averaged as mean ± sd. The significant correlations were estimated using Spearman's rank order correlation coefficient $|rho| \geqslant 0.75$, $p \leqslant 0.05$, corrected according to the Benjamini-Hochberg method[24]. Relative cell abundances were used for the analyses. SC: all subcommunities, Para: biotic and abiotic bulk parameters (Fig. S3.1, without DW).

| community | Insular I phase | | $RC_{10}$ | | $RC_{50}$ | | $RC_{80}$ | | Insular II phase | |
|---|---|---|---|---|---|---|---|---|---|---|
| | SC vs. SC | SC vs. Para. | SC vs. SC | SC vs. Para. | SC vs. SC | SC vs. Para. | SC vs. SC | SC vs. Para. | SC vs. SC | SC vs. Para. |
| L1 | 234 | 11 | 149 | 20 | 284 | 52 | 259 | 37 | 320 | 52 |
| L2 | 250 | 14 | 272 | 28 | 308 | 71 | 193 | 56 | 182 | 24 |
| L3 | 227 | 8 | 234 | 17 | 363 | 64 | 316 | 47 | 263 | 35 |
| L4 | 234 | 26 | 364 | 45 | 359 | 72 | 246 | 41 | 210 | 53 |
| L5 | 158 | 21 | 228 | 18 | 255 | 64 | 274 | 55 | 277 | 51 |
| mean ± sd | 221 ± 36 | 16 ± 7 | 249 ± 78 | 26 ± 12 | 314 ± 47 | 65 ± 8 | 258 ± 45 | 47 ± 8 | 250 ± 55 | 43 ± 13 |
| R | 283 | 69 | 292 | 30 | 295 | 68 | 335 | 44 | 319 | 99 |

To test whether the newly assembled community members also established an interconnected relationship and changed their interaction potential due to mass transfer, the numbers of significant correlations of SC vs. SC and between SC and the immediate environment of the microorganisms, SC vs. Para (biotic and abiotic parameters) were counted based on relative cell numbers in SCs. Generally, more significant correlations were found for SC vs. SC and of those more in the recycling phases RC (Table S13.2). The highest number of strong correlations for SC vs. SC in L1-L5 was found for RC$_{50}$ (mean = 314 ± 47) and RC$_{80}$ (mean = 258 ± 45; Wilcoxon test: pairwise comparison, $p$ = 0.012 [Insular I vs. RC$_{50}$], 0.222 [RC$_{10}$ vs. RC$_{50}$], 0.151 [Insular II vs. RC$_{50}$] and 0.143 [Insular I vs. RC$_{80}$], 0.691 [RC$_{10}$ vs. RC$_{80}$], 1 [Insular II vs. RC$_{80}$], respectively). The number of correlations was much lower for SC vs. Para; however, the trend was the same for RC$_{50}$ (mean = 65 ± 8) and RC$_{80}$ (mean = 47 ± 8; Wilcoxon test: pairwise comparison, $p$ = 0.112 [Insular I vs. RC$_{50}$], 0.012 [RC$_{10}$ vs. RC$_{50}$], 0.027 [Insular II vs. RC$_{50}$] and 0.008 [Insular I vs. RC$_{80}$], 0.032 [RC$_{10}$ vs. RC$_{80}$], 0.548 [Insular II vs. RC$_{80}$], respectively). These results show the highest interaction potentials of SC vs. SC and SC v. Para in phases RC$_{50}$ and RC$_{80}$, suggesting newly assembled communities by mass transfer.

A number of SCs profited exclusively from the increase in the recycling rates $RC$ in phases RC$_{50}$ and RC$_{80}$ (Fig. S12.1, linear correlation coefficient |rho| $\geqslant$ 0.55, $p \leqslant 0.05$). Of those, gates G5, G12, G14 and G33 showed the highest cell numbers (> $1\times10^{8}$ cells mL$^{-1}$ on average in RC$_{80}$, each) and among them G5, G12 and G14 were also those that were nested during RC$_{50}$ or RC$_{80}$ (Table S8.4). The cells of these SCs were sorted and analysed by 16S rRNA gene sequencing (Supplementary Information S9, step2). The number of interactions between the various SCs was found to be generally higher than the interactions between SCs and operational parameters. The low number of interactions between SCs and the operational parameters was probably caused by the continuous setup of the reactor system. Instead, the high number of interactions between SCs under these conditions is contrary to the expected unaltered cell states typical of continuous cultivation of pure cultures.

However, the connected netgrowth and rescue qualities of certain SCs under mass transfer, especially at RC$_{50}$ and RC$_{80}$ (Fig. S12.1), can be considered responsible for reinforcing their selection and, concomitantly, the (reversible, phase Insular II) decline of the other SCs (Fig. S12.2) can be considered a process that increased the interaction potential between the SCs.
